# Magnetoelectric nanodiscs diminish motor deficits in a model of Parkinson’s disease

**DOI:** 10.1101/2025.06.04.657885

**Authors:** Ye Ji Kim, Antoine De Comite, Emmanuel Vargas Paniagua, Sharmelee Selvaraji, Ethan Frey, Rajib Mondal, Nasim Biglari, Rebecca Leomi, Nidhi Seethapathi, Polina Anikeeva

**Author notes:** All correspondence should be addressed to: N.S. and P.A.

## Abstract

Magnetoelectric nanodiscs (MENDs) offer a wireless, minimally invasive route to neuromodulation by converting weak magnetic fields into electric polarization, but their therapeutic applications have remained unrealized. Here, we report the therapeutic development of MENDs for deep brain stimulation (DBS) in the subthalamic nucleus (STN) to alleviate motor deficits in a mouse model of Parkinson’s disease. To quantitatively assess therapeutic outcomes, we introduce GaitPattern, a computational pipeline that extracts DBS-induced alterations in salient gait features. Using GaitPattern, we demonstrate that MEND-mediated STN DBS induces motor improvements with precision comparable to clinical electrode-based DBS, while minimizing inflammatory responses associated with the implanted hardware. Notably, MEND-mediated DBS reduces oxidative stress in the brain, a key contributor to neurodegeneration. These findings position MEND-mediated STN DBS as an effective and minimally invasive neuromodulation strategy.

## Introduction

Parkinson’s disease (PD) characterized by the degeneration of dopaminergic (DA) neurons in the substantia nigra pars compacta (SNpc), a key node of the nigrostriatal motor pathway, is the fastest growing neurodegenerative motor disorder globally.^1–4^ The progressive degeneration of the DA neurons in the SNpc manifests in resting tremor, bradykinesia, rigidity, and postural instability – the hallmark features of PD.^4,5^ Early-stage PD is primarily treated with medications that diminish the motor symptoms, and their dosage increases as the disease progresses.^4–6^ However, following an initial “honeymoon” period, PD medications are often characterized by variable efficacy and side effects including dyskinesias.^4–7^ Notably, available medications are unable to halt the progressive degeneration of DA neurons.^8^

Deep brain stimulation (DBS) via electrodes implanted in the subthalamic nucleus (STN) is an effective therapy that alleviates motor symptoms associated with PD.^9,10^ However, the deployment of DBS poses risks associated with intracerebral implantation surgery, infection, and glial scar formation around the rigid millimeter-thick metal electrodes.^11^ Given these challenges as well as a limited effectiveness of DBS for non-motor symptoms, this therapy is currently available only to a small fraction of PD patients who meet the eligibility criteria outlined by the Core Assessment Program for Surgical Interventional Therapies in PD (CAPSIT-PD), which includes the average time from the diagnosis to STN DBS of ∼13 years.^12,13^

While electrode-based DBS remains a standard of care for late-stage PD, the technical risks associated with its invasiveness motivate the development of alternative neuromodulation strategies. Experimental approaches based on optogenetics,^14^ ultrasound,^15^ and magnetic fields^16^ aim to provide effective contactless neuromodulation without the need for implanted electrodes. Among these, magnetic field-based techniques offer an advantage of deep-brain penetration, unobstructed by skin or skull.^17,18^ Previous studies have leveraged biochemically benign magnetic nanomaterials as transducers to convert magnetic energy into heat,^19^ mechanical forces,^20,21^ and chemical stimuli.^22,23^ These stimulation strategies typically relied on specialized protein machinery, which had to be delivered via transgenic approaches with limited clinical potential. Recently, magnetoelectric transducers with dimensions ranging from nanometers to millimeters have enabled direct conversion of magnetic signals into changes in electric potential sufficient to excite neurons in vitro and in vivo, without requiring any genetic modification.^24,25^

This study explores the efficacy of neuromodulation mediated by synthetic magnetoelectric nanodiscs (MENDs, 250 nm diameter, 50 nm thickness) as an alternative approach to STN DBS in a mouse model of PD. Here, MENDs composed of a Fe_3_O_4_ core and consecutive shells of magnetostrictive CoFe_2_O_4_ and piezoelectric BaTiO_3_, generate electric polarization in response to biologically benign magnetic fields, enabling remote and transgene-free neuromodulation.^24^ In a hemi-Parkinsonian 6-hydroxydopamine (6-OHDA) mouse model, we compare the effects of MEND- and electrode-based STN DBS using computer-vision methods to analyze gait characteristics. Our observations suggest that MEND-based DBS could not only offer symptomatic relief but also pave the way for future minimally invasive approaches to improve PD patients’ motor deficits.

## Results and Discussion

### Neuromodulation mediated by magnetoelectric nanodiscs

In this work, we explore the effects of remote neuromodulation mediated by magnetoelectric nanodiscs (MENDs) in the context of a neurodegenerative disease (**Fig. 1a**). The synthesis of hexagonal magnetoelectric nanodiscs (MENDs) with a Fe₃O₄-CoFe₂O₄-BaTiO₃ double-core shell architecture and a diameter of 252±27 nm and thickness of 46±7 nm was performed as previously described (Methods).^24^ The elemental composition of MENDs was corroborated via electron diffraction spectroscopy (EDS) performed on high-angle annular dark-field scanning transmission electron microscopy (HAADF-STEM) images of the MENDs (**Fig. 1b,c** and Supplementary Fig. 1). Elemental mapping via EDS on scanning electron microscopy (SEM) images revealed the presence of MENDs as marked by Fe on neuronal cultures incubated with these particles at a concentration of 0.145 mg/ml in 24-well plates, resulting in a surface density of 0.75 µg/mm^2^ (**Fig. 1d**, Supplementary Fig. 2). The efficacy of the magnetoelectric conversion by the MENDs was quantified via a three-electrode electrochemical cell reported previously.^24^

**Fig. 1.**
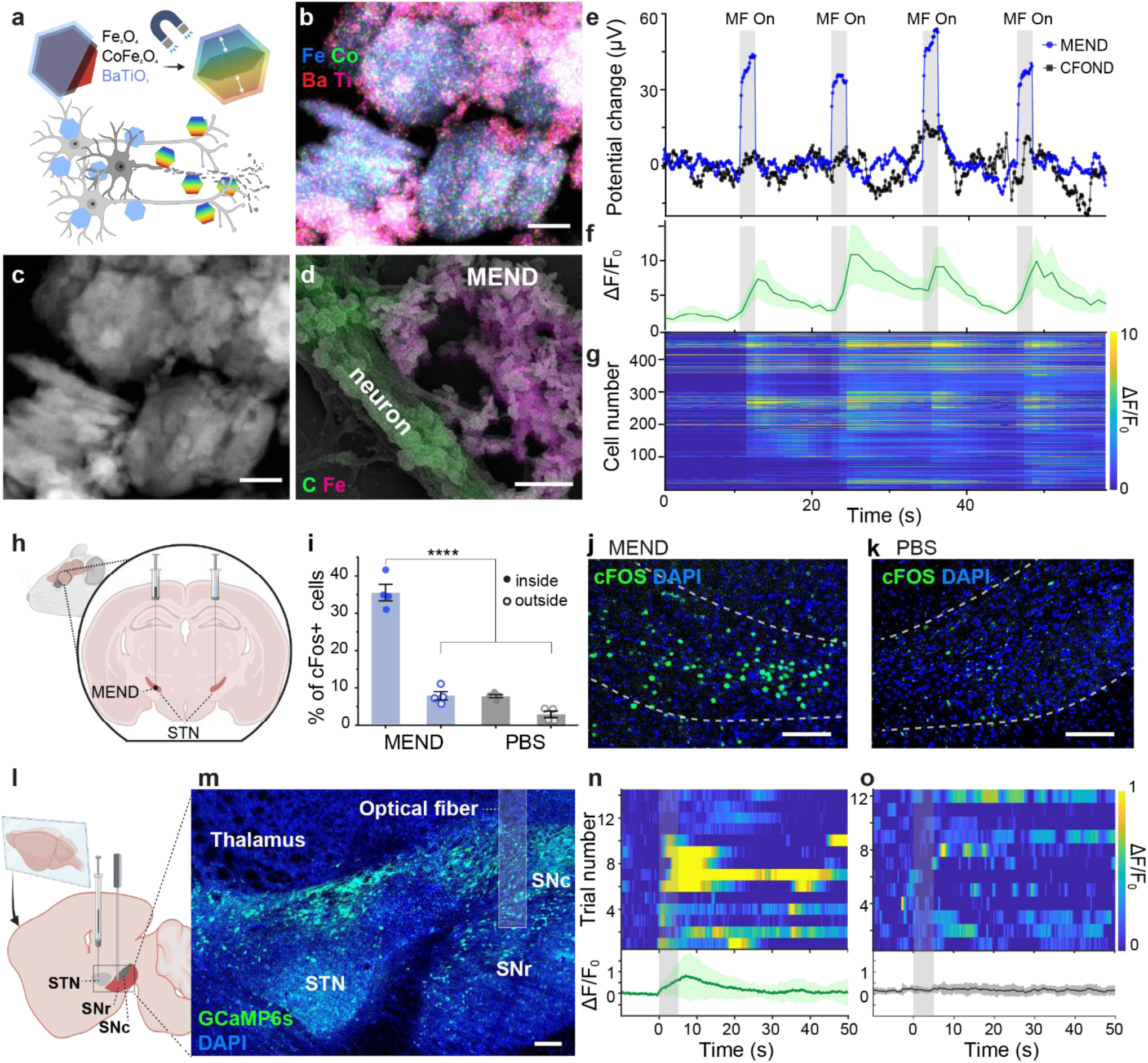
***In-vitro* and *In-vivo* neuromodulation mediated by MENDs. a** Schematic illustration of the MEND-mediated treatment on neurodegeneration. **b** EDS mapping of electron diffraction spectroscopy for Fe, Co, Ti, and Ba on **c** High-angle annular dark-field scanning transmission electron microscopy (HAADF-STEM) image of MENDs. Scale bars indicate 100 nm. **d** SEM image of neurons decorated with MENDs, overlayed with the EDS mapping. Scale bar is 1 𝜇m. **e** Potential changes in the electrochemical cell, including the electric polarization generated by MENDs under magnetic field application on the working electrode. **f** Average and **g** individual traces of GCaMP6s relative fluorescence changes on cultured hippocampal neurons. Grey shaded areas indicate the magnetic field (MF) epochs. (**f**) Line and shaded area represent mean and standard error of the mean (s.e.m.), respectively. **h** Schematic illustration for MEND injection in left STN and PBS injection in right STN. **i** Quantification of c-Fos expression in the STN (inside, solid data symbol) and surrounding regions (outside, open data symbol) from the images of (**j**) left and (**k**) right sidesafter the exposure the magnetic field. Scale bar, 100 µm. The normality of the dataset was confirmed with the Shapiro–Wilk test, and the statistical analysis was performed with one-way ANOVA followed by Tukey’s test; F(3,12)=123.4, p<0.0001.**** for p < 0.0001. The data is shown with mean±standard deviation (SD). The scale bars in (**j**) and (**k**) are 100 µm. **l** Schematic sagittal view of the left mouse brain illustrating injection of MENDs and AAV for GCaMP6s expression in the STN for photometry recording from the SNr. **m** Representative confocal image of a sagittal brain section from the left hemisphere showing GCaMP6s expression; green indicates GCaMP6s fluorescence and blue indicates DAPI nuclear staining. Scale bar, 500 µm. **n–o** GCaMP6s normalized fluorescence change (ΔF/F_0_) traces recorded from the left SNr following magnetic field stimulation (5 s; 220 mT OMF and 15 mT, 100 Hz AMF; gray bar). Top panels show individual trial traces, and bottom panels show the trial-averaged response for mice with (**n**) MEND injection and (**o**) PBS injection in the left STN. Grey shaded areas indicate MF epochs. Lines and shaded areas represent mean and s.e.m.

A combined magnetic field comprising a constant 220 mT offset magnetic field (OMF) and an alternating magnetic field (AMF) with a frequency of 100 Hz and amplitude of 15 mT was previously shown to elicit electric polarization in Fe₃O₄-CoFe₂O₄-BaTiO₃ MENDs which was sufficient to drive neuronal excitation.^24^ The selection of the OMF magnitude was motivated by extensive prior optimization, which identified 220 mT as the condition that yields the peak magnetoelectric coupling coefficient.^24^ For the AMF, it was previously shown that increasing its amplitude and frequency enhances the magnetoelectric coupling coefficient. However, a frequency of 100 Hz was chosen because it reliably elicits neural responses synchronized with the magnetic field application, while an amplitude of 15 mT represents the maximum achievable output of our experimental setup described in Supplementary Fig. 3. Here, these conditions similarly generated synchronized magnetically-evoked potential fluctuations in the electrochemical cell where the working electrode was coated with MENDs at a density of 2 µg/mm^2^ (**Fig. 1e**). Notably, no synchronized potential fluctuations were observed when a working electrode was coated with cobalt-ferrite coated magnetite nanodiscs (Fe₃O₄-CoFe₂O₄, CFONDs, 2 µg/mm^2^, Fig.1e).

The neuromodulatory efficacy of MENDs was then evaluated in primary hippocampal neurons expressing a fluorescent calcium indicator GCaMP6s (Supplementary Fig. 4). In neurons decorated with MENDs at a surface density of 1 µg/mm² a combined magnetic field (220 mT OMF; 15 mT, 100 Hz AMF) applied in 2s epochs separated by 10-second rest intervals evoked GCaMP6s fluorescence increase synchronized with the field exposure (**Fig. 1f,g**). Consistent with prior work, these results suggest the use of MENDs for temporally precise remote neuromodulation.

The effectiveness of MEND-mediated stimulation was then examined in the subthalamic nucleus (STN) of wild-type (WT, C57BL/6) mice. In these mice (n=4, 2 male, 2 female) MENDs (1.5 µl, 2 mg/ml) were injected into the left STN, while phosphate-buffered saline (PBS) was injected into the right STN to control for the potential effects of the magnetic field alone. Following a one-week recovery period, these mice were exposed to five 5-second epochs of a combined magnetic field (220 mT OMF; 10 mT, 100 Hz AMF) separated by 25-second intervals. We then applied immunohistology to analyze the expression of c-Fos across the STN and adjacent brain regions. The *c-fos* gene or its protein product c-Fos are widely used as markers of neuronal activation^26^, but increased c-Fos expression can be also be induced by inflammation or repair process following mechanical tissue injury following the intracranial injection^26–29^. To account for the potential effects of the injection procedure itself, c-Fos expression was compared between the MEND-injected and PBS-injected hemispheres within the same animals. We observed significantly higher expression of c-Fos in the left STN (injected with MENDs) as compared to right STN (injected with PBS control) (**Fig. 1i-k** and Supplementary Fig. 5 and 6).

Additionally, to assess the spatial precision of MEND-mediated stimulation in the STN, we compared c-Fos expression within and outside the STN (**Fig. 1j** and Supplementary Fig. 5 and 6). The expression of c-Fos was significantly higher within the STN in the MEND-injected hemisphere compared to the ipsilateral regions outside the STN as well as with both intra- and extra-STN regions on the contralateral PBS-injected hemisphere. Akin to prior observations, these findings indicate the utility of

MENDs and the combined magnetic field for driving neuronal excitation in the deep brain of mice. To ensure consistency, the MEND volume and concentration as well as magnetic field conditions (OMF, AMF, epoch, and rest-interval duration) described here were then applied throughout the manuscript unless otherwise specified.

To investigate how MEND-mediated stimulation engages neural circuits underlying motor behavior, we performed in vivo fiber photometry recordings in the substantia nigra pars reticulata (SNr), a downstream target of STN neurons, as prior studies have demonstrated that both electrical and optogenetic stimulation of the STN evoke robust responses in the SNr ^30^. Three weeks following injection of an adeno-associated viral vector carrying GCaMP6s gene under a pan-neuronal human synapsin promoter (AAV9-hSyn::GCaMP6s, 200 nL of 1×10¹³ vg/mL) together with MENDs (1.5 µg, 2 mg mL⁻¹) or PBS into the left STN, GCaMP6s expression was observed in the left SNr (**Fig. 1l,m**). A prominent increase in GCaMP6s fluorescence was observed in response to the applied magnetic fields in the MEND group, whereas no significant response was detected in the PBS control group (**Fig. 1n,o**).

Altered synaptic organization within basal ganglia circuits, including a weakened STN–SNr pathway and reduced excitatory input from the motor cortex, are commonly observed in models of PD^31–33^. Within the context of these dysregulated circuits, in addition to the excitation in the SNr evoked by STN DBS, we also observed increased c-Fos expression in the primary motor cortex (Supplementary Fig. 7). This increased c-Fos expression in the primary motor cortex potentially occurs through the recruitment of established basal ganglia-thalamocortical projection circuits^34,35^. These findings indicate that MEND-mediated neuromodulation is not restricted to the local cell bodies at the injection site and more broadly engages the motor circuits.

### Hemi-Parkinsonian mouse model

To assess the potential of MEND-mediated deep brain stimulation (MEND-DBS) as a therapeutic alternative to DBS via implanted electrodes, we employed a common unilateral mouse model of PD driven by an injection of 6-hydroxydopamine (6-OHDA, 3 µg per mouse, 15 mg ml^-1^) into the medial forebrain bundle (MFB), following previously published protocol^36^ and stereotactic coordinates^37^. This model manifests in a substantial loss of DA neurons in the SNpc in the 6-OHDA injected hemisphere, mimicking DA neurodegeneration in PD (**Fig. 2a**). 6-OHDA is a catecholamine-selective neurotoxin that is taken up intracellularly by the noradrenaline and DA transporters ^38–40^, leading to neurotoxic effects driven by reactive oxygen species (ROS) ^40,41^ and mitochondrial dysfunction ^42^ (**Fig. 2b**). Consistent with the prior reports of 6-OHDA PD models^36,38,43–45^, two weeks following a unilateral (left) MFB injection, a 45±14% of tyrosine hydroxylase (TH)-expressing DA SNpc neurons remained at the injected side, and the population decreased to 16±10% four weeks following surgery. We also observed significant loss of DA neurons in the ventral tegmental area (VTA), with the loss of TH-positive neurons from 76±14% to 40±6.1% during the same time period (**Fig. 2c-f**, Supplementary Fig. 8). It was previously shown that DA neuron degeneration in the SNpc largely plateaus by ∼3 weeks following 6-OHDA injection into the MFB^36,46^. Accordingly, here we selected time points before and after this observed degeneration plateau to capture both the ongoing disease-like progression as well as the advanced degree of pathology. This timeline provides insight into the progression of DA degeneration in the 6-OHDA PD mouse model, informing the potential therapeutic window to explore the effects of MEND-DBS on motor behaviors in this model.

**Fig. 2.**
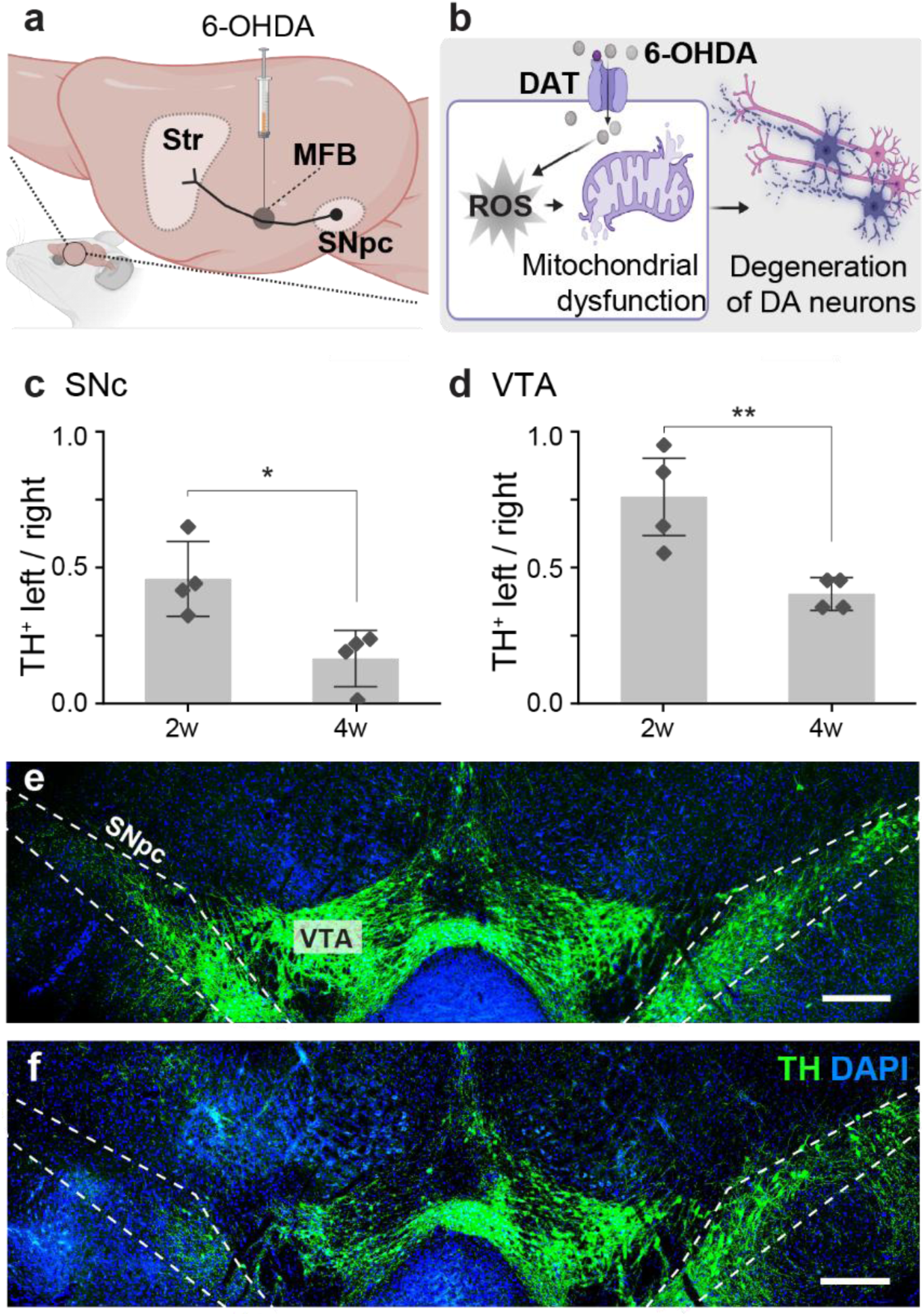
Unilateral 6-OHDA Parkinson’s disease (PD) model. **a** Schematic illustration of 6-OHDA PD mouse model. **b** Mechanisms underlying 6-OHDA-mediated degeneration of dopaminergic neurons. **c, d** Quantification of immunohitology of TH-stained dopaminergic neurons in (**c**) SNpc and (**d**) VTA. Scale bars, 500 µm. Markers represent individual subjects; bars and error bars represent mean±standard deviation (SD), respectively. The statistical analysis was performed with two sample t-test; (**c**) p=0.0145,t=3.3999, DF=6 (**d**) p=0.00361, t=0.6220, DF=6; * for p < 0.05, ** for p < 0.01. **e** 2 weeks and **f** 4 weeks after 6-OHDA injection surgery.

The unilateral 6-OHDA medial forebrain bundle (MFB) model was selected to ensure the robust evaluation of our MEND-modulation approach and validation of the behavioral analysis pipeline. This model exhibits robust motor impairments with relatively uniform lesion severity, enabling sensitive detection of treatment-induced behavioral changes, as compared to slower progressive models such as Mitopark or MPTP-based paradigms^47,48^, motivating its use for technology validation and direct comparison to electrode DBS “gold standard”.

### Evaluation of the effects of MEND-DBS on gross motor deficits in hemiparkinsonian mice

To compare the effects of MEND-DBS to traditional electrode-based DBS on PD-like motor symptoms in mice, we performed 6-OHDA injections into the left MFB with injections of MENDs, or PBS (negative control), or implantation of the electrodes (positive control) in the left STN during the same surgery. This combination of manipulations created three experimental groups that were evaluated for motor function at 4 (test #1) and 5 weeks (test #2) following the injection/ implantation surgeries (**Fig. 3a**): mice injected with 6-OHDA and PBS (Group 1, n=7); mice injected with 6-OHDA and implanted with an electrode (Group 2, n=8); and mice injected with 6-OHDA and MENDs (Group 3, n=10). An additional group (Group 4, n=11, 6-OHDA in MFB and MENDs in STN) was introduced to investigate the effects of earlier-stage MEND-DBS on the progression of DA neuron degeneration and motor function. In this group, motor symptoms were evaluated at 2 (test #1) and 5 weeks (test #2) following surgery (**Fig. 3a**). All experimental groups were compared to a naïve control group (Group 5, n=6) that were similarly evaluated for motor symptoms at 4 (test #1) and 5 weeks (test #2) (**Fig. 3a**). Except for the animals implanted with DBS electrodes (Group 3), all mice were exposed to the combined MEND-driving magnetic field, resulting in sham stimulation for Groups 1 (PBS) and 5 (naïve).

**Fig. 3.**
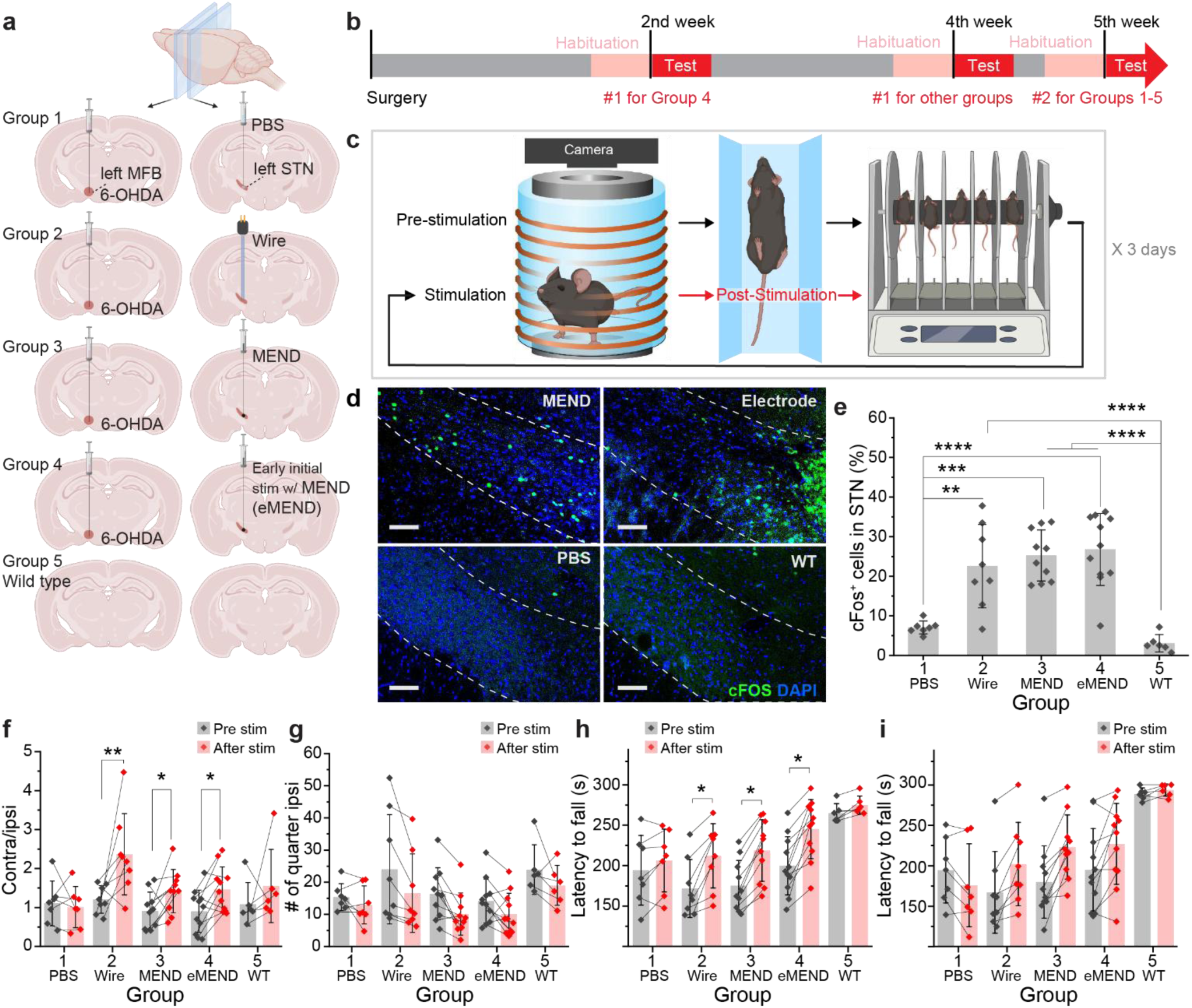
MEND-mediated STN DBS effects on motor functions in a unilateral 6-OHDA model. **a** Experimental Groups 1-5. Schematic illustration of **b** timeline for behavior assay and **c** sequence of each test, including cylinder test, gait analysis, and rotarod. **d** c-Fos expression in STN instilled with MENDs, electrodes, and PBS and STN of wild type. Scale bars, 100 µm. STN is marked with dashed line. **e** Quantification of c-Fos expression in STN of Groups 1-5. The normality of the dataset was confirmed with the Shapiro–Wilk test, and the statistical analysis was performed with One-way ANOVA followed by Tukey’s test; F(4, 37)=16.91, p<0.0001. **f** Number of quarters ipsilateral turns in the cylinder test and **g** ratio of number of contralateral turns to that of ipsilateral turn in the cylinder test. (**f, g**) Each data point represents the average of six days (three days from Test #1 and three days from Test #2 for each animal) of measurements per animal. The statistical analysis was performed with two-way repeated measures ANOVA; (**f**) F_Group_(4,37)=2.03, P_Group_=0.129, F_Stim_(1,37)=10.5, P_Stim_=2.31e-2, F_Interaction_(4,37)=1.85, P_Interaction_=0.159; (**g**) F_Group_(4,37)=0.796, P_Group_=0.542, F_Stim_(1,37)=19.9, P_Stim_=6.66e-3, F_Interaction_(4,37)=0.999, P_Interaction_=0.431, and Holm-Bonferroni post-hoc test confirmed pairwise pre vs post differences (**f**) Group 1: p=0.7574, Group 2: p=0.0076, Group 3: p=0.0498, Group 4: p=0.0397, Group 5: p=0.5328; (**g**) Group 1 p=0.6958, Group 2 p=0.0971, Group 3 p=0.3056, Group 4 p=0.2843, Group 5 p=0.3527. **h,i** Rotarod test result analyzed with change in latency to fall pre- and post-stimulation latency to fall, at (**h**) Test #1 and (**i**) Test #2. (H) Each data point reflects the mean of three days of data collected during Test #1 from each animal. (I) Each data point is the average of measurements per animal for three days during Test #2. The statistical analysis was performed with two-way repeated measures ANOVA; (**h**) F_Group_(4,37)=9.13, P_Group_=0.0118, F_Stim_(1,37)=70.2, P_Stim_=2.31e-2, F_Interaction_(4,37)=1.91, P_Interaction_=0.174; (**i**) F_Group_(4,37)=6.23, P_Group_=0.049, F_Stim_(1,37)=13.3, P_Stim_=3.54e-2, F_Interaction_(4,37)=2.90, P_Interaction_=0.13, and Holm-Bonferroni post-hoc test confirmed pairwise pre vs post differences (**h**) Group 1: p=0.573, Group 2: p=0.0408, Group 3: p=0.0415, Group 4: p=0.0458, Group 5: p=0.4602; (**i**) Group 1 p=0.6429, Group 2 p=0.1753, Group 3 p=0.1795, Group 4 p=0.7242, Group 5 p=0.8069. * for p < 0.05, ** for p < 0.01, *** for p < 0.001, **** for p < 0.0001, p > 0.5 is not indicated. (**e-i**) Bars and error bars represent mean±SD, respectively.

Locomotion assays were performed according to the experimental timeline outlined in **Fig. 3b**. Each test session spanned three days, with a standardized sequence of assays repeated daily to enhance statistical rigor (Methods). This sequence of assays comprised a 3 min pre-stimulation cylinder test to quantify rotations, a 15 min cat-walk assay for gait analysis, a rotarod test (5 min maximum) to assess motor coordination and balance, a 30-min rest in the home cage, followed by 3 min stimulation within the cylinder test, and concluded with post-stimulation 15 min cat-walk and rotarod (5 min maximum) assays (**Fig. 3c**). The animal was assumed to be exposed to a spatially uniform magnetic field generated by the solenoid coil wound around the cylindrical test setup, flanked by two flat-faced parallel permanent magnets separated by 8 cm gap comparable to the 7.5 cm diameter of the magnets^49,50^. Immediately following the behavioral assay sequence (90 min after the stimulation), the mice were sacrificed via transcardial perfusion with paraformaldehyde, and the expression of c-Fos was quantified in the STN to assess the extent of neuronal activation in each animal. Robust upregulation of c-Fos expression was observed in mice subjected to MEND-mediated (Groups 3 and 4) and electrode (Group 2) DBS as compared to control Groups 1 and 5 (**Fig. 3d,e** and Supplementary Fig. 9,10). The c-Fos–positive cells were defined as those where c-Fos immunofluorescence (marked by the secondary antibody labeled with Alexa Fluor 488, Methods) overlapped with a nuclear stain DAPI. C-Fos expression is confined to the nucleus, and any green fluorescence outside nuclear regions was interpreted as autofluorescence noise. The latter was particularly prominent within the regions damaged by the electrode implantation (Supplementary Fig. 10), and these autofluorescence signals were excluded from analysis because they did not colocalize with DAPI.

We then analyzed the effects of MEND- and electrode-DBS on motor function in all experimental groups. Notably, the mice that received MEND (Groups 3 and 4) or electrode (Group 2) DBS in the left STN showed an increase in the ratio of contralateral to ipsilateral turns compared to both the naïve (Group 5) and 6-OHDA and PBS injected (Group 1) animals (**Fig. 3f**), despite no significant reduction in the ipsilateral turns (**Fig. 3g**). The unilateral 6-OHDA injections into the nigral dopaminergic circuit were previously shown to yield ipsilateral rotations^51^, and the latter could be diminished following treatment with DA agonists or electrode DBS^52,53^. We previously observed that MEND-mediated left STN DBS in healthy WT mice led to an increase in contralateral rotations.^24^ In contrast, in hemiparkinsonian mice

MEND- or electrode-mediated STN DBS did not yield significant changes in contralateral turns (Supplementary Fig. 11a). Despite the absence of statistically significant changes in ipsilateral or contralateral turning when analyzed separately, the notable shift in their ratio indicates that MEND and electrode DBS significantly affect movement directionality in hemiparkinsonian mice, and the effects are distinct from those observed in healthy mice.

While our work does not employ any stimulants commonly used to unmask subtle phenotypic features in unilateral PD models^53,54^, our observations of ipsilateral rotational bias prior to stimulation in all 6-OHDA groups align with the prior findings (Supplementary Fig. 11b). Both MEND and electrode DBS in the STN were effective at reducing bias to ipsilateral rotations, indicating a partial rescue of the motor phenotype. However, the extent of reduction in ipsilateral rotations did not differ significantly between groups (Supplementary Fig. 11c-e).

To further evaluate the impact of MEND and electrode DBS on motor function, we assessed the performance of all groups in the rotarod test, which offers insights into balance, coordination, and physical condition based on the animal’s ability to remain on the rod.^44,55^ Although no significant differences in latency to fall were observed across the groups receiving 6-OHDA injections (Groups 1-4), all of these groups showed significant differences when compared to naïve mice (Group 5) at baseline, prior to stimulation (Day 1 of Test #1) (Supplementary Fig. 12a). Notably, the MEND- (Groups 3 and 4) and electrode-DBS (Group 2) groups exhibited improvements in latency to fall at subsequent post-stimulation time points (**Fig. 3h**) in Test #1, and significant inter-group differences were detected between MEND-DBS groups (Group 3,4) and naïve mice (Group 5) (Supplementary Fig. 12b). At Test #2, neither MEND-nor electrode-DBS showed significant differences between pre- and post-stimulation conditions (**Fig. 3i**). However, significant inter-group differences were observed in both electrode- and MEND-DBS groups (Group 2-4) as compared to sham-stimulation 6-OHDA controls (Group 1) (Supplementary Fig. 12c). These findings indicate that both electrode and MEND DBS improve motor functions evaluated via a rotarod.

### Evaluation of the effects of MEND DBS on movement velocity

In addition to the effects on movement directionality (cylinder test) and overall coordination (rotarod), we assessed the effects of MEND and electrode DBS on gait and velocity, critical parameters disrupted in PD. To extract salient movement features from cat-walk assays, we expanded upon computer-vision methods^56^ to develop an automated analysis pipeline (GaitPattern). The movement of mice inside a column (7.5 cm width × 8 cm height × 150 cm length) was first traced via DeepLabCut^56^ applied to videos recorded at 60 frames per second (Supplementary Video 1). The trace was then used in the GaitPattern pipeline to extract velocity, step length, step length asymmetry, and diagonal support, as these parameters have been previously shown to be affected by disrupted nigrostriatal signaling.^57–59^

First, we examined how velocity changes in the groups injected with 6-OHDA. The naïve mice (Group 5) exhibited significantly higher movement velocity as compared to the 6-OHDA-injected, sham-stimulated group (Group 1) in Test #1 (z = 15.4, p = 1.68 × 10⁻⁵³), and the difference became more pronounced in Test #2 (z = 23.1, p = 1.11 × 10⁻¹¹⁷) (**Fig. 4a**). The progressive decline in motor function following 6-OHDA injection was further corroborated via the bootstrap mean difference velocity (Group 5 – Group 1) 31.059 mm/s with 95% bootstrapped confidence interval (CI) [26.105, 35.984] mm/s and p=1.00×10^-4^ at Test#1 advancing to 51.689 mm/s with 95% bootstrap CI [46.842, 56.498] mm/s and p=1.00×10^-4^ at Test#2 (**Fig. 4a**).

**Fig. 4.**
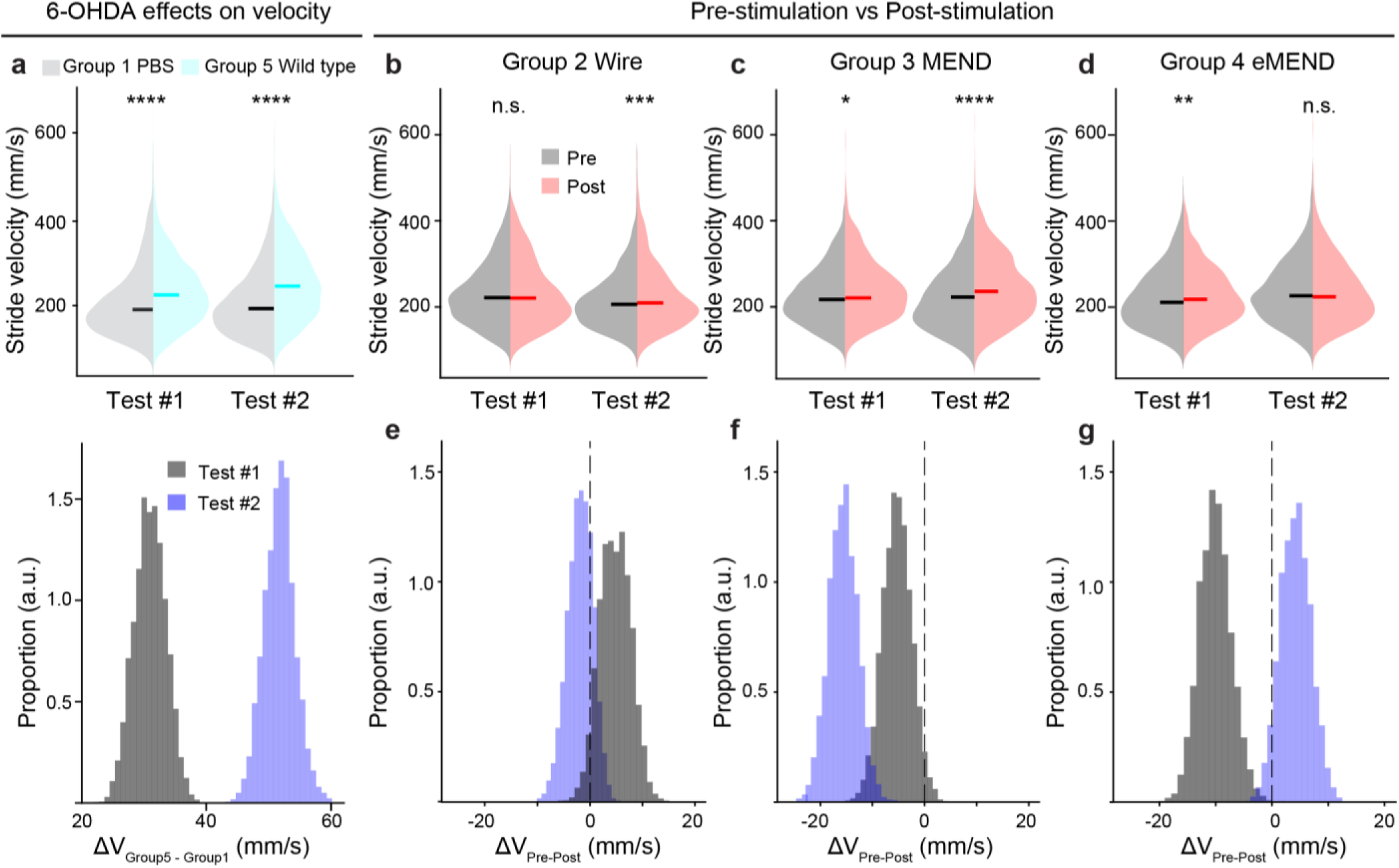
Automated analysis of velocity across stimulation and control conditions in a unilateral 6-OHDA model . **a** Velocity distribution (top) and bootstrap resample(bottom) for Group 1 (6-OHDA in MFB and PBS in STN) and Group 5 (naïve) mice at Tests #1 and #2. The data obtained by integrating all steps for both pre- and post-stimulation epochs. The statistical analysis for the histogram was performed via Wilcoxon rank sum test; Test #1: W= –15.1; Test #2: W=–22.2. For Test #1, the bootstrap mean difference (ΔV_m_, _Group5 – Group1_) = 31.1 mm/s with bootstrap 95% confidence interval (CI) = [26.1, 36.0] mm/s, and bootstrapped p=1.00×10^-4^. For Test #2, ΔV_m_, _Group5 – Group1_ = 51.7 mm/s with CI = [46.8, 56.5] mm/s, p=1.00×10^-4^. **b-d** Velocity distribution before and after stimulation in Group 2 (**b**), Group 3 (**c**), and Group 4 (**d**). The statistical analysis was performed via Wilcoxon rank sum test (**b**) Test#1 W= – 0.522, Test #2= –3.59; (**c**) Test #1 W=–1.08, Test #2= –6.46; (**d**) Test #1 W= –3.17, Test #2=1.01. (**a-d**) Each plot displays the integrated velocity per step across all animals within each group, based on data collected over three days for each test. **e-g** Bootstrap resampling of the velocity data shown in **b-d** of Group 2 (**e**), Group 3 (**f**), and Group 4 (**g**) at Test #1 (dark grey) and Test #2 (blue). (**e**) Test #1: ΔV_m, Pre-Post_ = 4.81 mm/s, CI = [–0.735, 10.4] mm/s, p = 2.19×10^-1^; Test #1: ΔV_m, Pre-Post_ = –1.86 mm/s, CI = [–6.84, 2.53] mm/s, p = 2.96×10^-5^; (**f**) Test #1: ΔV_m, Pre-Post_ = –5.29 mm/s, CI = [–10.8, 0.178] mm/s, p = 2.51×10^-1^; Test #2: ΔV_m, Pre-Post_ = –15.6 mm/s, CI = [–21.2, –10,2] mm/s, p = 1.81×10^-11^; (**g**) Test #1: ΔV_m, Pre-Post_ = –10.1 mm/s, CI = [–15.4, –4.26] mm/s, p = 2.15×10^-5^; Test#2: ΔV_m, Pre-Post_ = 4.18 mm/s, CI = [–0.995, 9.47] mm/s, 𝛽 = 0.822 mm/s, p = 6.94×10^-1^. In the bootstrap plots, black dashed line indicates 0.

These comparisons were performed by integrating the step data from 3 days for each of the tests (Methods). The integration was performed to account for daily fluctuations in velocity (Supplementary Fig. 13). For instance, in mice not receiving any stimulation (Groups 1 and 5) statistically significant yet opposite changes were observed following sham stimulation on different test days (Supplementary Fig. 13). These physiological fluctuations that may arise from minor (0.5-2 hr) misalignment of circadian cycles across test days and other environmental factors underscore the importance of considering the reliability of daily changes when analyzing other gait parameters.

As noted above, our study did not use stimulants commonly leveraged to amplify locomotor deficits in PD models. Our automated analysis revealed significant differences in velocity between Groups 1 (6-OHDA) and 5 (naïve) at the pre-stimulation phase of Test 1 (**Fig. 4a**), however comparisons between averaged velocities of individual animals did not show the significant difference (Supplementary Fig 14a). Notably, at Test #2 (week 5 post injection) naïve mice (Group 5) showed significantly higher average velocity than the 6-OHDA (Group 1) animals. Notably, Group 5 animals had higher velocity than the 6-OHDA-injected Groups 1, 2, and 3, even following the stimulation at Test 2. Interestingly, Group 4, which received the 6-OHDA injection and initial stimulation at week 2, showed no significant differences from the naïve group (Supplementary Fig. 14b).

To further examine these group-level differences, we conducted a step-level analysis by integrating all steps across the three test days for each animal. As shown in **Fig. 4b,c**, STN DBS using both electrodes and MENDs (Groups 2 and 3) significantly increased movement velocity immediately after stimulation in Test #2. In contrast, the 6-OHDA and PBS injected (Group 1) and naïve (Group 5) animals did not show significant changes at either Test #1 or Test #2 (Supplementary Fig. 15). Notably, the effects on velocity diverged between the two stimulation methods over time. While both MENDs and electrode-based DBS yielded increased velocity immediately following stimulation, the mice receiving electrode DBS showed a decrease in velocity between Test #1 and Test #2, whereas animals receiving MEND DBS did not exhibit such a decline (Supplementary Fig. 16).

Group 4 mice exhibited increased velocity following stimulation at Test #1 (2 weeks post 6-OHDA injection), however these effects were not observed at Test #2 (**Fig. 4d**), where these mice already showed velocities not statistically different from the naïve animals (Group 5, Supplementary Fig. 14b). Together, these observations suggest that MEND DBS offers greater benefits if initiated earlier in the progression of DA neuron degeneration.

To test whether the observed differences reflected effects of stimulation rather than biological data variability, we conducted bootstrap resampling and regression analyses. At Test #1, Groups 2 and 3 showed no reliable stimulation-related effects (**Fig. 4e, f**). In contrast, Group 4 exhibited a robust increase in velocity following stimulation at Test #1, with a bootstrap CI entirely below zero [−15.4, −4.4 mm/s] and a significant regression relationship (p = 2.10 × 10⁻⁵) (**Fig. 4g**). At Test #2, significant stimulation effects also emerged in Groups 2 and 3 (**Fig. 4e,f**). Group 2 showed a significant regression relationship despite a confidence interval that partially overlapped zero with small variability (CI: [−6.9, 2.5 mm/s], p = 2.96 × 10⁻⁵), while Group 3 demonstrated a strong and consistent reduction in velocity (CI: [−21.0, −10.1 mm/s], p = 1.81 × 10⁻¹¹). Group 4 did not exhibit stimulation-related changes at Test #2 (**Fig. 4g**).

To assess the potential influence of sex on stimulation effects, we examined STN DBS–induced changes in movement velocity separately in male and female animals across experimental groups. When stratified by sex, Group 2 (electrode DBS) showed no stimulation-associated change in velocity at Test #1 in either males or females. At Test #2, however, both sexes exhibited a stimulation-associated increase in velocity, reflected by negative bootstrap mean differences (female p = 5.53 × 10^−4^; male p = 1.15 × 10^−2^) despite the corresponding bootstrap CIs partially overlapping with zero (Supplementary Fig. 17a). In Group 3, MEND-mediated stimulation produced robust stimulation-associated increases in velocity in males at both Test #1 and Test #2, while significant stimulation effect was only observed in females at Test #2 (Supplementary Fig. 17b). In Group 4, which received early MEND-mediated stimulation, both male and female animals exhibited significantly higher velocity following stimulation at Test #1 (Supplementary Fig. 17c). Overall, sex-stratified analyses revealed patterns consistent with those observed in the pooled data, indicating that sex did not qualitatively alter the temporal or group-specific stimulation effects on movement velocity.

While sex differences in susceptibility to DA lesions have been reported in clinical PD and 6-OHDA models, with female animals often showing greater resilience, whether STN DBS efficacy differs by sex remains unclear^60–62^. Consistent with prior reports of comparable motor benefits of DBS in men and women despite sex-specific differences in non-motor outcomes, our observations indicate similar stimulation-induced improvements in velocity across sexes. Another factor that can affect the data variability is the duration of the behavioral assays. To probe this, we analyzed the effects of stimulation on velocity on a per-minute basis and found no significant time-dependent effects across the experimental groups (Supplementary Fig. 18).

The velocity has previously been studied alongside mobility or distance traveled in the open-field test to evaluate the effects of STN DBS. Our analysis of distance traveled during a 15-min gait assay revealed no significant differences between pre- and post-stimulation conditions (Supplementary Fig. 19). However, total movement distance does not capture step-level mobility, and comparisons of averaged velocities across individual animals failed to reveal significant differences even at baseline conditions (Supplementary Fig. 14a). In contrast, step-level velocity analysis detected statistically significant differences between 6-OHDA–injected mice (Group 1) and naïve mice (Group 5) (**Fig. 4a**). Previous studies also observed subtle increases in movement velocity or mobility post-stimulation when evaluating averages across individual animals^35,63,64^, although these changes were not highlighted because pre- versus post-stimulation differences were not statistically significant, whereas significant increases were observed during stimulation. Thus, our step-level analysis pipeline provides a more sensitive approach for detecting subtle changes in mobility that would otherwise be missed by approaches considering whole body movement alone.

### Evaluation of the effects of MEND DBS on gait symmetry

Step-level analysis using GaitPattern enables a more precise evaluation of gait, capturing subtle step-by-step features beyond integrated locomotor measures. Given the unilateral DA degeneration in our hemiparkinsonian model, we assessed the right-left gait asymmetry across groups. At both pre- and post-stimulation epochs of Tests #1 and #2, significant differences were observed between the right and left step lengths in all mice injected with 6-OHDA (Groups 1-4), while naïve controls (Group 5) exhibited comparable step lengths (**Fig. 5a,b** and Supplementary Fig. 20). To quantify the effects of stimulation on these lateral differences, we defined a parameter step length asymmetry (SLA) shown in Equation (1):

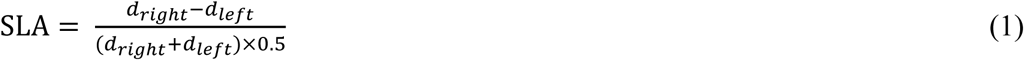

where 𝑑_𝑟𝑖𝑔ℎ𝑡_ and 𝑑_𝑙𝑒𝑓𝑡_ are the distances between the front and hind paws on the right and left sides, respectively.

**Fig. 5.**
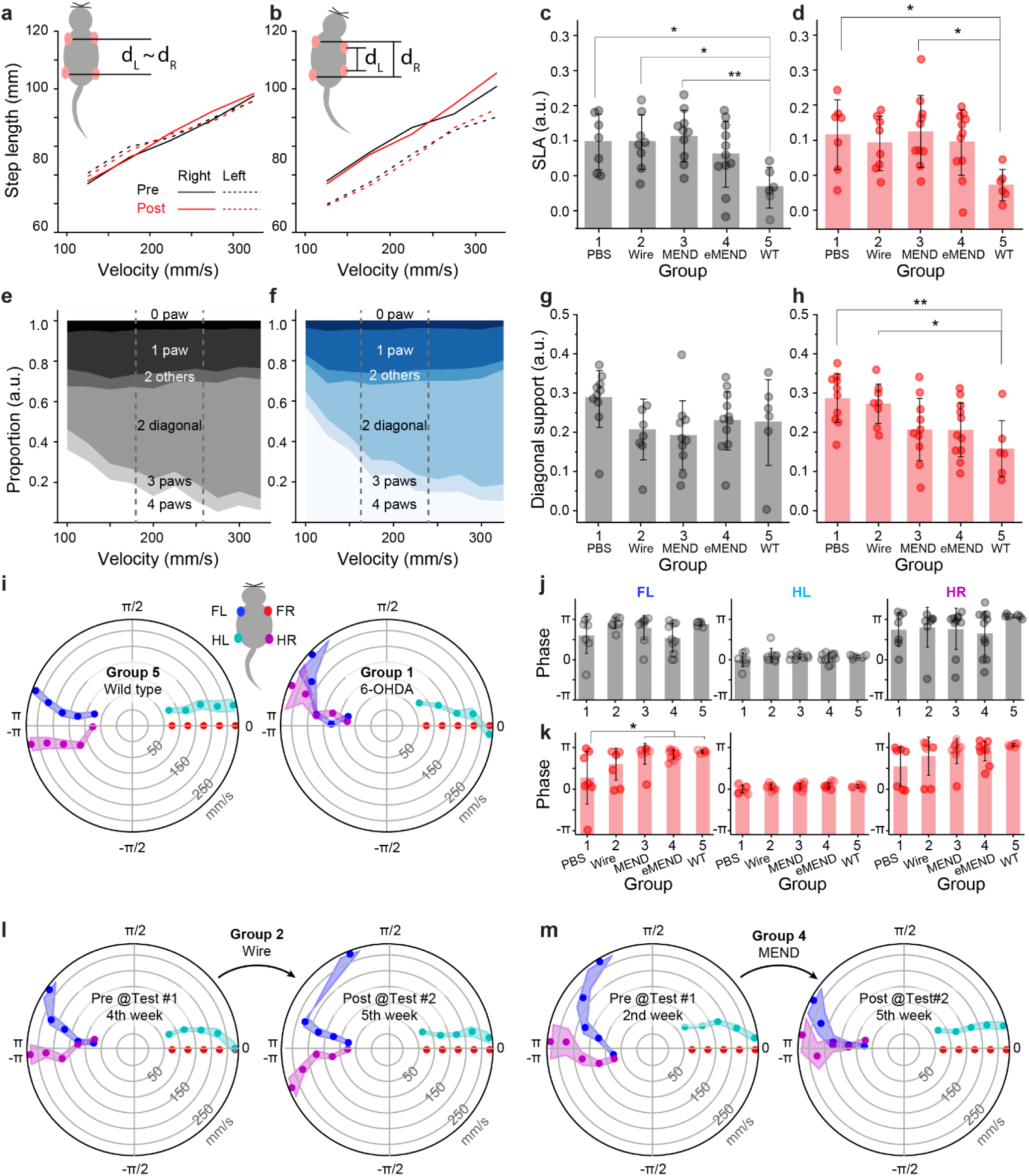
Automated analysis of gait changes in response to 6-OHDA injections and STN DBS. a,b. Right and left step lengths as function of velocity for Group 1(**a**) and Group 5 (**b**), averaging the data from each step across all animals within each group over three days for Test#2. **c** Baseline SLA measured prior to the initial stimulation at day 1 of Test #1. **d** SLA following stimulation at Test #2, where each data point represents the average measurement per animal during the post-stimulation session over three days in Test #2. The statistical analysis was performed via one-way ANOVA followed by Tukey’s test; (**c**) p=0.0109, F(4, 37)=3.80 (**d**) p=0.0254, F(4, 37)=3.14. **e,f** Support variation along with the movement velocity of (**e**) Group 1 and (**f**) Group 5, averaging measurement per animal during the post-stimulation session over three days in Test #2. **g** Baseline diagonal support at low velocity measured prior to stimulation at day 1 of Test#1. **h** The post-stimulation diagonal support averaged over three days during Test #2. (**g,h**) Black markers and error bars represent the mean and standard deviation for each group. The statistical analysis was performed with One-way ANOVA followed by Tukey’s test; (**g**) p=0.2027, F=1.56, and (**h**) p=0.00490, F=4.39. **i** Polar plots indicating the phase of the step cycle in which each limb enters stance relative to stance onset of FR paw for Group 1 (left) and Group 5 (right). Radial axis represents walking velocity. Limbs are color coded as shown in the inset; each data point represent averages across animals within each velocity bin. **j** Baseline (pre-stimulation at the first day of Test#1) and **k** post-stimulation at Test#2 averaged phase at speed 200-300 mm/s for each paw (FL first column, HL second column, and HR third column). The statistical analysis was performed with One-way ANOVA followed by Tukey’s test; (**j**) FL F(4,37)=2.12, p=0.098; HL F(4,37)=0.90, p=0.47; HR F(4,37)=0.73, p=0.58; (**k**) FL F(4,37)=3.94, p=9.19×10^-3^; HL F(4,37)=1.17, p=0.34; HR F(4,37)=2.06, p=0.11. * for p < 0.05, ** for p < 0.01, *** for p < 0.001, **** for p < 0.0001, n.s. for p > 0.05.(**c,d,g,h,j,k**) Black markers and error bars represent mean±SD. **l,m** Polar plots showing the phase of the step cycle at which each limb enters stance relative to the FR paw’s stance onset for (l) Group 2 (electrode STN DBS 6-OHDA) and (m) Group 4 (MEND early STN DBS, 6-OHDA) at baseline (left, Pre-stimulation on the first day of Test#1) and Post-stimulation at Test #2 (right). (**i, l, m**) Markers and shaded areas represent circular mean±s.e.m.

**Fig. 6.**
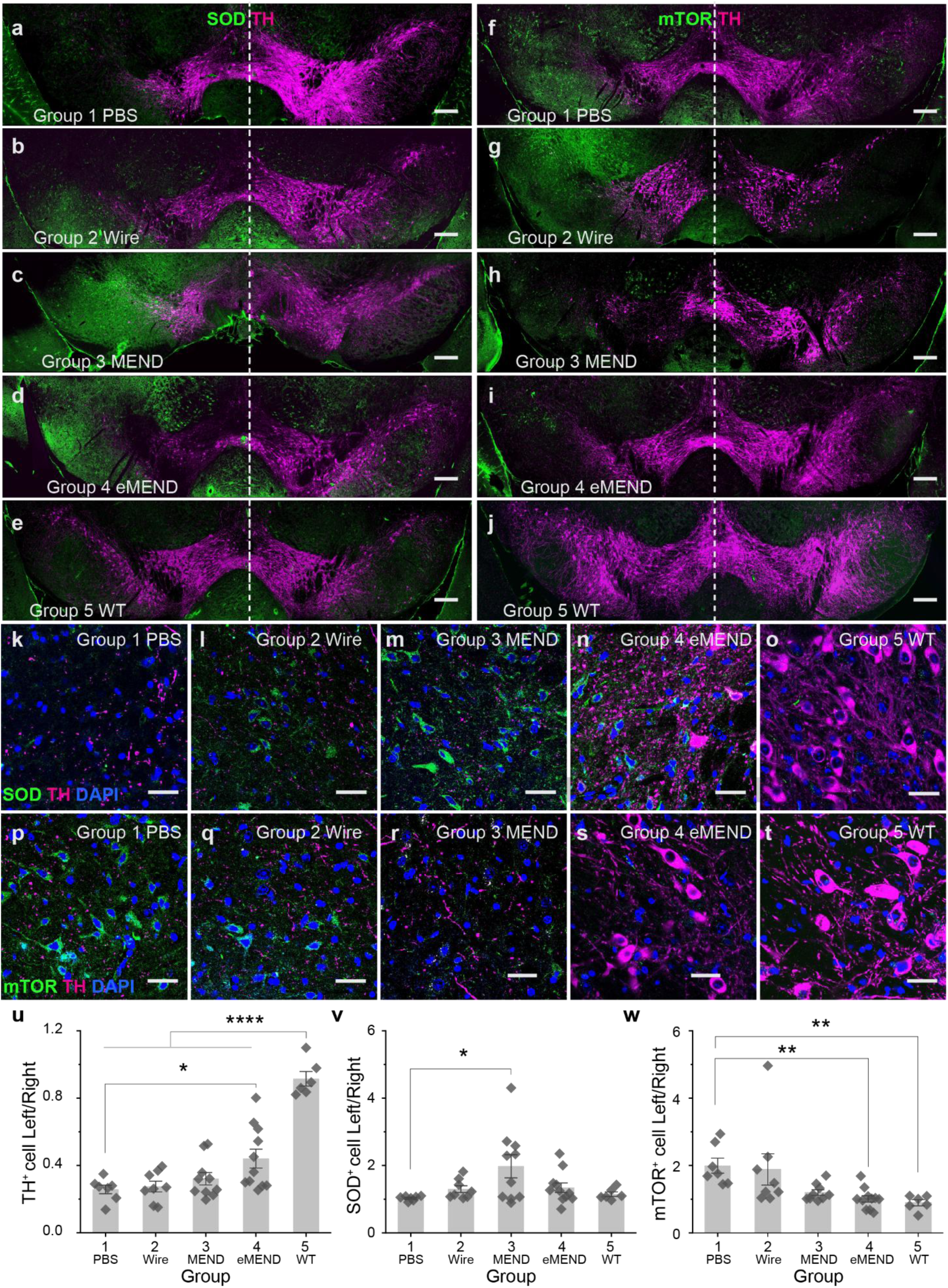
DA neurons degeneration and oxidative stress changes in response to MEND-mediated STN DBS. a-e. SOD and TH staining on SNpc of (**a**) Group 1, (**b**) Group 2, (**c**) Group 3, (**d**) Group 4, and (**e**) Group 5. **f-j** mTOR and TH staining on SNc of (**f**) Group 1, (**g**) Group 2, (**h**) Group 3, (**i**) Group 4, and (**j**) Group 5. (**a-j**) Scale bar is 200 µm. **k-o** Magnified view of SOD, TH, and DAPI staining on SNpc of (**k**) Group 1, (**l**) Group 2, (**m**) Group 3, (**n**) Group 4, and (**o**) Group 5. **p-t** Magnified view of SOD, TH, and DAPI staining on SNpc of (**p**) Group 1, (**q**) Group 2, (**r**) Group 3, (**s**) Group 4, and (**t**) Group 5. (**k-t**) Scale bars, 50 µm. **u-w** Quantification of (**u**) TH-, (**v**) SOD-, and (**w**) mTOR-expressing cell percentage in the left SNpc normalized to that on the right hemisphere. (**u-w**) The Shapiro–Wilk test was performed to test the normality of data distribution. (**u,v**) The statistical analysis was performed with one-way Anova followed by Tukey’s test; (**u**) F(4,37)=29.1, p<0.0001, (**v**) F(4,37)=3.27, p= 0.0214. As Group 2 in (**w**) rejected normality, the statistical analysis was performed with the Kruskal-Wallis test followed by post-hoc Dunn’s test; H=21.3, DF=4, p= 2.81e-4. * for p < 0.05, ** for p < 0.01, *** for p < 0.001, **** for p < 0.0001, n.s. for p > 0.05. (**u-w**) Makers represent individual subjects; bars and error bars represent mean±SD.

On day 1 of Test #1 – corresponding to week 4 for Groups 1–3 and 5, and week 2 for Group 4 – prior to stimulation all groups receiving 6-OHDA injections (Groups 1-4) exhibited greater SLA than the naïve Group 5 (**Fig. 5c**). Although Group 4, which was assessed at week 2 post-injection, also showed elevated SLA relative to Group 5, this difference did not reach statistical significance. This lack of significance suggests that the extent of DA degeneration two weeks post-6-OHDA injection was not sufficient to manifest in gait asymmetry.

We found that DBS (electrode or MEND-mediated) had no immediate effect on SLA in either Test #1 or #2 in any of the groups (Supplementary Fig. 21). However, we observed longer-term differences in SLA between the groups. At Test #2 post-stimulation, Groups 2 (electrode DBS) and 4 (early onset of MEND DBS) exhibited low SLA not statistically different from the naïve Group 5 (**Fig. 5d**). Meanwhile, Group 3 (late onset MEND DBS) exhibited significantly greater SLA than the controls. This may be attributed to the lower stimulation intensity of MEND-DBS, or the reduced efficacy of delayed intervention with nanomaterials which may be partially internalized by the tissue^24^ (as Group 3 received initial stimulation at 4 weeks post 6-OHDA injection). This suggests that electrode DBS reduces SLA associated with unilateral DA neuron degeneration, and MENDs-mediated stimulation offers comparable benefits when administered early in the degeneration progression.

We additionally assessed ipsilateral rotational bias in 6-OHDA lesioned animals during the catwalk assays. Consistent with the cylinder test results, the 6-OHDA-injected groups exhibited ipsilateral rotational bias whereas naïve controls showed no bias at both Test#1 and #2 (Supplementary Fig. 22 a,b). To account for inter-animal variability in total distance traveled the number of rotations was normalized to this distance for each subject. Notably, unlike in the cylinder test, where stimulation was applied during the assay, we did not observe differences in ipsilateral or contralateral turns during the catwalk tests when comparing pre- and post-stimulation conditions for either electrode or MEND DBS (Supplementary Fig. 22c-f). This indicates that while STN DBS does not provide lasting effects on the ipsilateral rotational bias after the stimulation, it provides benefits during active stimulation.

Next, we compared variations in diagonal support, a contact mode where the body weight is supported by two diagonally opposite limbs (for example, left front and right hind) during quadrupedal locomotion. Diagonal support provides stability and balance, allowing the other two limbs to move forward. The diagonal support increases with the movement velocity, as shown in the phase diagram for all animals in our behavioral assays (Supplementary Fig. 14), which contributes to a wider base of support. Using this diagram, the velocity range can be divided into three segments: low (100 – 183.3 mm s^-1^), medium (183.3 – 266.7 mm s^-1^), and high (266.7 - 350 mm s^-1^). The most notable differences in diagonal support between 6-OHDA injected and naïve subjects were observed in the low velocity range (**Fig. 5e,f** and Supplementary Fig. 23).

We observed that the effects of unilateral 6-OHDA injections on the diagonal support emerged later as compared to SLA. On day 1 of Test #1, no significant differences in diagonal support were observed across all groups and between stimulation conditions at low velocities (**Fig. 5g**). Similar observations were made on days 2 and 3 of Test #1 as well. However, by day 2 of Test #2, 6-OHDA injected and sham-stimulated (Group 1) and electrode-stimulated (Group 2) mice showed significantly larger diagonal support at low velocities as compared to naïve controls (Group 5) at pre-stimulation time points (Supplementary Fig. 24). In addition, there was no significant change in diagonal support immediately after the stimulation at either Test #1 or Test #2 (Fig. S25). Given that at low velocities the differences between the sham-stimulated hemiparkinsonian animals and naïve controls were most pronounced at the latest experimental timepoint, Test #2 post stimulation epoch (**Fig. 5h**).

Beyond diagonal support, temporal analysis of interlimb coordination during symmetrical trot patterns across a wide range of walking speeds in naïve mice^65^ was assessed by examining the phase of the step cycle at which each limb enters a stance (i.e. time when the limb contacts with the floor) relative to a stance onset of a reference limb. The front right (FR) limb was designated as the reference, and step-cycle phases were analyzed using polar plots across movement-speed bins (50–100, 100–150, 150–200, and 250–300 mm/s) for front left (FL), hind right (HR), and hind left (HL) limbs. Naïve mice (Group 5) exhibited a characteristic trot pattern in which diagonal limb pairs moved synchronously (i.e., near-zero phase difference in FR–HL and FL–HR pairs) and alternated with the opposite diagonal pair (approximately ± π phase difference in FR–FL or FR–HR pairs) (**Fig. 5i**). This interlimb coordination was markedly disrupted in hemiparkinsonian 6-OHDA mice, particularly at higher walking speeds (**Fig. 5j**).

To quantify inter-group differences and stimulation effects, we averaged the phase of each paw for each animal within the high-velocity range (200–300 mm/s). Consistent with other gait metrics (e.g., step-length asymmetry and diagonal support), neither electrode nor MEND DBS produced an immediate improvement in interlimb coordination at Test #1 or Test #2 following stimulation (Supplementary Fig. 26). However, MEND-mediated stimulation altered the longitudinal progression of gait-phase coordination.

At baseline (**Fig. 5j**) and at Test #1 post-stimulation (Supplementary Fig. 27), no significant inter-group differences were observed. By Test #2 post-stimulation (**Fig. 5k**), FL steps in both Group 3 (MEND DBS at 4 weeks) and Group 4 (MEND DBS at 2 weeks) showed a significantly improved trot pattern compared to Group 1 (sham-stimulated 6-OHDA), with effect sizes comparable to the difference between naïve mice (Group 5) and Group 1 (Supplementary Fig. 28). Interestingly, electrode DBS failed to restore gait-phase coordination toward the naïve-like pattern (**Fig. 5l**). In contrast, MEND-mediated stimulation shifted gait-phase relationships toward those observed in healthy naïve controls, with the strongest effects observed when stimulation was initiated earlier (**Fig. 5m**).

Our observations suggest that distinct motor deficits in a 6-OHDA model emerge over different timescales. For instance, velocity (Fig. 4a) and SLA (Fig. 5a) were significantly affected by 6-OHDA by 4 weeks post-injection, whereas diagonal support (Fig. 5g) and gait-phase (Fig. 5j) remained relatively unaffected over the same period. This dissociation suggests that different components of locomotor control exhibit differential vulnerability to DA loss and may rely on partially separable circuit dynamics. Furthermore, while both electrode and MEND DBS resulted in comparable improvements in velocity, their effects on long term gait pattern outcomes diverged. Notably, neither stimulation modality induced immediate changes in diagonal support or interlimb coordination following stimulation (Supplementary Fig. 21 and 26). Instead, stimulation effects on these gait parameters evolved over time; significant differences were observed between 6-OHDA-injected sham-stimulation group (Group 1) and naïve animals (Group 5) by Test#2 (Fig. 5h,k). Groups receiving MEND stimulation did not show significant differences relative to naïve animals, whereas groups receiving electrode stimulation developed motor deficits comparable to those in the sham-stimulation group.

Although some DBS effects on gait asymmetry, such as diminished ipsilateral rotational bias (shown in Fig. 3f), appear to dissipate rapidly once the stimulation is turned off (Supplementary Fig. 22), our data support an alternative pattern that emerges over longer timescales, potentially driven by plastic changes within motor circuits. Such plasticity-dependent mechanisms may underlie the gradual emergence of inter-group differences observed in this study, particularly for locomotor parameters related to coordination and bilateral gait symmetry. These behavior-specific and temporally distinct effects of stimulation are consistent with clinical observations from PD patients receiving DBS. While STN DBS produces sustained improvements in tremor, rigidity, and bradykinesia, its effects on gait and posture appear transient^66–73^. These differences are hypothesized to stem from disparate underlying mechanisms, with rapid and reversible desynchronization of pathological subcortical activity contrasting with slower, experience-dependent reorganization of distributed motor networks^74^. Our findings in the 6-OHDA model align with this framework, supporting the idea that DBS-mediated motor improvements arise from a combination of immediate network modulation and longer-term plastic adaptations that vary across motor circuits.

### MEND stimulation suppresses oxidative stress

Given the diverging trends in motor deficits between groups, we investigated the extent of DA neurodegeneration across all conditions. Despite receiving the same dose of 6-OHDA, the extent of DA neuron depletion, as marked by a decrease in the relative number of TH-expressing cells in the left SNpc, varied between Groups 1-4 (**Fig. 6a-u**). Group 1 (sham stimulation; **Fig. 6a,f,k,p**) exhibited significantly greater DA depletion in the left SNpc compared with Group 4 (2-week–onset MEND-DBS; Fig. **6d,j****,n,s,u**). In contrast, Groups 1–4 (sham stimulation, electrode DBS, 4-week–onset MEND-DBS, and 2-week–onset MEND-DBS) all showed significant reductions in TH-positive neurons in the left SNpc relative to the naïve control group (Group 5; **Fig. 6u**).

Although the mechanisms underlying DA neuron degeneration in PD remain vigorously investigated, changes in dopamine metabolism, immune activation, mitochondrial dysfunction, oxidative stress, and impaired autophagy have been associated with the disease progression motivating further inquiry into the effects of DBS on these processes.^75–78^

For instance, the level of superoxide dismutase (SOD), an antioxidant enzyme, appear to correlate negatively with the severity of PD symptoms.^79^ Treatments targeting reactive oxygen species (ROS), such as CuxO nanoparticle clusters (which mimic SOD activity) ^80^ or iron chelation with Deferiprone,^81^ have been shown to reduce oxidative stress and improve motor function. Following these insights, we compared SOD expression in the substantia nigra (SN, divided into SNpc and SN pars reticula, SNr) across the five experimental groups in our study (**Fig. 6a-e,k-o,v**, Supplementary Fig. 29, 30). The SNpc is a primary DA-producing region in the basal ganglia motor circuit, and SNr is the major area receiving excitatory projections from STN. Therefore, the entire SN area was assessed for changes in the SOD levels in response to the STN DBS. All DBS Groups (2 – electrode, 3 and 4 – MEND) exhibited higher relative levels of SOD expression between the left (stimulated) and right (unstimulated) SN as compared to sham-stimulated (Group 1) and naïve (Group 5) animals. Although the increases in Groups 2 and 4 were not statistically significant relative to the sham and naïve groups, the effect was significant for Group 3 (MEND DBS) (**Fig. 6v**).

In addition to antioxidant enzymes, autophagy processes, particularly mitophagy, have been suggested to play a role in regulating oxidative stress, and impaired mitochondrial function has been linked to PD progression heterogeneity.^81,82^ Therefore, here we assessed the effects of DBS on the expression of mechanistic target of rapamycin (mTOR), which regulates mitophagy (**Fig. 6f-j,p-t,w**, Supplementary Fig. 29, 31). Unilateral injection of 6-OHDA yielded significantly higher expression of mTOR in the left (injected) hemisphere in sham-stimulated mice (Group 1) as compared to laterally symmetric and negligible expression observed in naïve (Group 5) mice (**Fig. 6f, j, m**). Interestingly, significantly lower activation of mTOR was observed in MEND-injected Group 4 (**Fig. 5i, s, w**).

Finally, to compare the biocompatibility of MEND-mediated DBS with electrode DBS, we assessed the markers of astrocytic (glial fibrillary acidic protein, GFAP)^83^ and glial activation (ionized calcium-binding adaptor molecule 1, Iba1; cluster of differentiation 68, CD68) as proxies for neuroinflammatory response. ^84,85^ Consistent with prior reports of foreign-body response to rigid brain implants,^86–89^ significantly higher expression of GFAP, Iba1, and CD68 was observed in the vicinity of DBS electrodes as compared to the MEND injection sites (Supplementary Fig. 33). The reduced mechanical invasiveness of 1.5 µl of 2 mg/ml MEND injections may, in turn, potentially contribute to the lower mTOR activation. Further inquiry into biophysical pathways may enable refinement of MEND-mediated DBS and help guide the optimal timing of its deployment in preclinical and clinical studies.

Although MEND-mediated STN DBS was associated with lower levels of oxidative stress and neuroinflammatory responses compared with electrode-based stimulation in this study, the acute nature of the 6-OHDA model limits interpretation of these findings in the context of long-term neuroprotection or disease modification. Future studies using more physiological and progressive PD models, such as Mitopark or α-synuclein accumulation bases models^90,91^, will be necessary to evaluate whether MEND-mediated stimulation can influence pathology progression over longer timescales.

## Conclusion

Our study demonstrates the potential of MEND-based neuromodulation as a less invasive therapeutic alternative to electrode-DBS for mitigating motor deficits in a mouse model of PD. Following 1.5 µl (2 mg/mL) injections into the STN, MENDs mediated remote magnetoelectric excitation of neuronal activity. In a unilateral 6-OHDA mouse model of PD, MEND-DBS improved motor functions. Notably, when applied earlier in a disease model progression, MEND-DBS reduced the extent of DA neuron degeneration and preserved motor function to a greater degree than the late-stage MEND-DBS, potentially due to reduced oxidative stress. Akin to electrode-DBS, MEND-DBS increased expression of the antioxidant enzyme SOD. However, unlike electrode-DBS, animals receiving MEND-DBS did not upregulate mTOR expression, which is typically elevated following 6-OHDA lesioning. Furthermore, the low mechanical invasiveness of MEND-DBS led to a reduction of inflammatory markers. Together, our findings illustrate the potential for less-invasive DBS to reduce oxidative stress while relieving motor symptoms in PD. Although this work was conducted in a murine model, the magnetic field parameters and apparatus designs readily align with previously reported human-scale implementations^24^.

## Materials and Methods

### Synthesis of MENDs

MENDs consist of a Fe_3_O_4_-CoFe_2_O_4_-BaTiO_3_ double-core-shell structure. The procedure described here follows the Methods of Ref. ^24^ and is provided here for clarity and convenience. The innermost core Fe_3_O_4_ magnetic nanodiscs (MNDs) were synthesized by reducing hematite nanodiscs that was produced by heating a uniform mixture of 0.273 g of FeCl_3_·6H_2_O (Fluka), 10 ml ethanol, 600 μl of deionized (DI) water, and 0.8 g of anhydrous sodium acetate (Sigma Aldrich) in a sealed Teflon-lined steel vessel at 180°C for 18 h. After washing the hematite nanodiscs with DI water and absolute ethanol 1:1 solution for three to five times, the nanodiscss are dispersed in 20 ml of trioctyl-amine (Sigma-Aldrich) and 1g of oleic acid (Alfa Aesar) and then reduced to magnetite at 370 °C (20 °C min^−1^) in H_2_ (5%) and N_2_ (95%) atmosphere for 30 min, connected to an evacuated Schlenk line. The second layer CoFe_2_O_4_ was synthesized with 257 mg cobalt acetylacetonate (Co(acac)_2_, Aldrich) and 706 mg iron acetylacetonate (Fe(acac)_3_, Aldrich) precursors dispersed in 20 ml diphenyl ether (Aldrich), 1.90 ml oleic acid (Sigma-Aldrich), and 1.97 ml oleylamine (Aldrich). After mixing the solution with dried MNDs, it was connected to the Schlenk line to evacuate and heat at 100 °C (7 °C min^−1^) for 30 min in an N_2_ atmosphere while magnetically stirring at 400 rpm. After closing the N_2_ line, the temperature was increased to 200 °C (7 °C min^−1^), maintained for 30 min, and then increased to 230 °C (7 °C min^−1^) and maintained for 30 min. After cooling the solution to room temperature, the CFONDs were washed three times with n-hexane and ethanol. The thickness of the 5 nm CoFe_2_O_4_ layer was achieved by repeating the organometallic synthesis and washing steps three times. The outermost layer, BaTiO_3_, was formed by the sol-gel method after coating CFONDs with poly(vinylpyrrolidone) (PVP, Sigma-Aldrich). Oil separation was performed on a mixture comprising 16 mg of CFONDs dispersed in n-hexane, 30 ml of DI water, 6 ml of ethanol, and 2g of PVP. After drying the solution in the vacuum at 80 °C, the amber-coloured gel was redispersed in a solution consisting of 0.5 g citric acid (Sigma-Aldrich) and 24 µl titanium isopropoxide (Aldrich) dissolved in 15 ml of ethanol and 0.1 g citric acid and 0.0158 g barium carbonate (Aldrich) dissolved in DI water. The solvents are evaporated at 80 °C for 12–14 h. The powders were heated at 600 °C for 2 h, 700 °C for 2 h, then 800 °C for 1 h, sequentially. The synthesized MENDs exhibit a magnetoelectric coupling coefficient of 92.2 mV mT^-1^ cm^-1^ under a magnetic field combining 220 mT OMF with 10 mT 100 Hz AMF.

### Electrode fabrication

The electrodes are placed in thermally drawn fiber. With CNC mill, two polycarbonate (PC) layers (McMaster, 8574K45) were machined to have dimensions of 10.7 × 16.7 mm^2^ and a 9.5 × 11.9 mm^2^ channel and dimensions of 6 × 16.7 mm^2^ and two 4.7 × 4.7 mm^2^ channels, respectively. A hot press at 185 °C and 5 psi for 60 minutes was applied to consolidate the PC layers. For 45-55 ratio of size reduction, the fiber was drawn in a vertical tower with a 3-zone furnace having temperatures of 140, 260, and 80 °C for the top, middle, and bottom furnaces, respectively. During the draw, spooled 100 µm stainless steel (SS) wires (Amazon, B0CB6C1X5Y) were fed into the pre-form and converged into the fiber. The drawn fiber was cut into 7 mm sections, and electrical connections were made to the SS wires. The wires were soldered onto 3D-printed copper traces connected to male header pins.

### Calcium imaging in cultured hippocampal neurons

Animal procedures were approved by the Massachusetts Institute of Technology Committee on Animal Care (protocol #2305000529). Hippocampus extracted from neonatal rat pups (P1, Sprague-Dawley, 001) were dissociated with Papain (Worthington Biochemical). The cells were then seeded on Matrigel (Corning)-coated glass slides (5 mm diameter) in 24-well plates at a density of 112,500 cells ml^−1^. The cells were incubated in 1 ml Neurobasal medium (Invitrogen) with glial inhibition using 5-fluoro-2’-deoxyuridine (FO503 Sigma) 3 days after seeding. Four days after seeding, the neurons were transduced with 1 µl of an adeno-associated virus serotype 9 (AAV9) carrying a fluorescent calcium ion indicator GCaMP6s (AAV9-hSyn::GCaMP6s, Addgene viral prep #100843-AAV9, >1 × 10^13^ IU ml^−1^).

1h before imaging, 50 µg of MENDs dispersed in DI water (2.9 mg/ml) was mixed with the neural basal in the well plate incubating the neurons. After the incubation, the glass coverslips entailing the neurons were then transferred into a custom sample holder containing 200 µl of Tyrode’s solution for imaging at an inverted microscope (Olympus IX73, 20X objective lens) integrated with the electromagnet applying magnetic field to the neurons (220 mT OMF combined with 10 mT 100 Hz AMF). The GCaMP6s fluorescence changes were recorded at imaging rate of 1 fps. The relative fluorescence changes Δ*F*/*F*_0_ was calculated with *F*_0_ average of fluorescence intensity (*F*) during 30s pre-stimulation and Δ*F* (*F*-*F*_0_).

### Surgical procedures for nanoparticles and 6-OHDA injections and electrode implantation

All experiments were carried out and approved by the Massachusetts Institute of Technology Committee on Animal Care (Protocol #2506000815). Adult male and female C57BL/6J mice (4-6 weeks old) were used in this study. Animals were housed under standard laboratory conditions (12-hour light/dark cycle; food and water ad libitum).

During the surgery, the mice were anesthetized using isoflurane (0.5 – 2.5% in Oxygen) using an anesthesia machine (VET EQUIP), and their eyes were covered with ophthalmic ointment. The animals’ core temperature was maintained with a heating pad. The fur on the top of the head was removed with depilatory cream after fixing the head into the stereotaxic surgery stage using ear bars. Following the sterilization of the skin with ethanol and betadine, a midline incision was made along the scalp. The injection or implantation was performed with the coordinates in the Mouse Brain Atlas^92^.

Parkinsonian symptoms were induced by unilateral injections of 0.2 µl of 15 mg/ml 6-hydroxydopamine (6-OHDA, 6-hydroxydopamine hydrobromide, Sigma-Aldrich 162957) into the left medial forebrain bundle (MFB, anterio-posterior (AP): –1.2 mm, medio-lateral (ML): –1.1 mm, dorso-ventral (DV): –5 mm) at a rate of 0.1 µl/min (3 µg total). To minimize the reflux of the 6-OHDA solution, the syringe had to be lifted at a speed ≤0.5 mm min^-1^. The three mice, in which the syringe was lifted at a faster speed did not show degeneration of TH-stained DA neurons (Fig. S14). During the same surgery, PBS, MENDs (1.5 µl of 2 mg/ml), or electrodes were administered in the left subthalamic nucleus (STN coordinates: AP −2.06 mm, ML −1.50 mm, DV −4.50 mm) using a microinjection apparatus (10 µl Hamilton syringe #80308, UMP-3 syringe pump and Micro4 controller; all from World Precision Instruments). The behavioral assays were conducted at two time points: 2 and 5 weeks after surgery for Group 4, and 4 and 5 weeks for all other groups. The mice were divided into five experimental groups:

1. Group 1: 6-OHDA PBS (n = 7, Female = 3, Male = 4)
2. Group 2: 6-OHDA electrode (n = 8, Female = 3, Male = 5)
3. Group 3: 6-OHDA MENDs (n = 10, Female = 6, Male = 4)
4. Group 4: 6-OHDA MENDs early stimulation (n = 11, Female = 6, Male = 5)
5. Group 5: Naïve control (n = 6, Female = 4, Male = 2)

### Behavioral assays

All experiments were carried out and approved by the Massachusetts Institute of Technology Committee on Animal Care (Protocol # 2506000814). Behavioral assays were conducted to assess motor functions at two time points: 2nd and 5th week after surgery for Group 4, and 4th and 5th weeks for Groups 1-3. Each test was performed over a 3-day period with the following sequence:

1. Pre-stimulation cylinder assay 3 min
2. Pre-stimulation gait analysis 15 min
3. Rotarod test 5 min at maximum
4. 30-minute resting period in the home cage
5. Stimulation session inside the cylinder assay 3 min
6. Post-stimulation gait analysis 15 min
7. Post-stimulation rotarod test 5 min at maximum

Every 3-day test period was performed after 3-day habituation period. During the habituation, the whole sequence was performed equally without stimulation.

### Cylinder test inside a magnetic field apparatus

The cylinder test was conducted in an enclosed arena consisting of an acrylic cylinder (inner diameter: 71 mm, height: 80 mm). A copper wire coil around the cylinder served as an electromagnet during the stimulation session, generating an alternating magnetic field (AMF). Additionally, a static offset magnetic field (OMF) was applied using two O-shaped permanent magnets (outer diameter: 7.5 mm, inner diameter: 4 mm; K&J Magnetics, Inc., RZ0X84) positioned at the top and bottom of the cylinder.

Each mouse’s rotational behavior was recorded using a camera placed directly above the cylinder, aligned with the top O-shaped magnet. The test included two sessions; for the pre-stimulation session, mice were placed in the cylinder without any applied stimulation, and for the stimulation session, mice were exposed to either a magnetic field or an electrode-based stimulation.

The stimulation session was performed with either magnetic field or electrode-based DBS. The applied field consisted of a static offset of 220 mT combined with a 10 mT, 100 Hz alternating magnetic field. Stimulation was delivered in five epochs, each lasting 5 seconds, with 25-second intervals between epochs. Otherwise, custom-made fibers containing electrodes (diameter: 100 µm) were used to deliver a 100 Hz, 20 ± 2 µA sine-wave stimulation. Stimulation followed the same temporal pattern as the magnetic field condition (five 5-second epochs with 25-second intervals). For this setup, the top O-shaped magnet was removed to allow electrode connection to a potentiostat (Gamry Interface 1010E).

Mouse rotation (ipsilateral and contralateral turns) was quantified from video recordings using custom-written code.

### Rotarod

The rotarod test was performed to assess motor coordination, balance, and physical condition. Mice were placed on a rotarod apparatus (Ugo Basile, Italy), which accelerated from 4 to 40 RPM over a 5-minute period. Only at the last day of habituation period, the mice were placed on the rotarod apparatus with the constant-speed loading mode. For the other former two-days of habituation period, the rotarod was skipped from the sequence. The time (latency) before falling from the rod was recorded. Each mouse underwent two trials with 10 min interval per session, and the average latency to fall was used for analysis.

### Gait analysis

Gait analysis was performed using a custom pipeline, which was developed to track and measure key gait parameters. Mice were placed inside a column (7.5 cm width × 8 cm height × 150 cm length) with a transparent bottom, and a camera placed underneath recorded their movements at 60 frames per second. The feet, nose, and tail positions were tracked across frames using markerless motion capture software^56^ and low pass filtered using a 6^th^ order double pass Butterworth filter. In the subsequent gait-analysis pipeline, we detected locomotion bouts based on a movement speed threshold of at least 75 mm/s for 0.5s, following a previous study^65^. Acceleration was not analyzed due to the requirement of the double differentiation that causes substantially increased rate of numerical errors and poor reliability in data rendering^93^. To identify the timing of foot contact, we computed the distance between the fore-aft position of the foot marker and the fore-aft position of thenose markers: the minima and maxima of this distance respectively correspond to the beginning and end of contact^94,95^. The contact location was then defined as the average location of the foot marker during the stance phase determined by these beginning and end of contact. Using this contact information, we extracted the following parameters: step length, step length asymmetry (SLA), diagonal support, and interlimb coordination. The movement velocity was defined as the average movement velocity of the fore-aft position of the nose marker, computed using 4^th^ order centered finite difference schemes, during one gait cycle (defined as the time interval between two successive contact initiations of the front right foot). The step length of a given pair of feet was defined as the fore-aft distance between the contact locations of these feet. The step length asymmetry was obtained by comparing the step length of the front right – hind right (𝐿_𝑟𝑖𝑔ℎ𝑡_) and front left – hind left (𝐿_𝑙𝑒𝑓𝑡_) pairs of feet as follows: 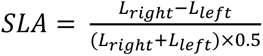. The contact modes, including the diagonal support, were computed as the percentage of time for which any number of feet, or only two diagonally opposed feet (front right-hind left or front left-hind right) for diagonal support, were in contact with the floor. Interlimb coordination was captured by computing the relative contact timing of each limb within the gait cycle defined as the time interval between two successive contacts of the front right limb. These relative contact timings were then expressed in radians with a relative contact timing of 1 corresponding to π radians.

### Immunohistochemistry

All experiments were carried out and approved by the Massachusetts Institute of Technology Committee on Animal Care (Protocol # 2506000814). After the last stimulation session during the behavior assays, each mouse was sacrificed by a lethal intraperitoneal injection of sodium pentobarbital (Fatal-plus, 20 mg ml^-1^, dose 100 µl per mouse). After checking no response with toe pinch, transcardial perfusion was performed with 4% paraformaldehyde (PFA) solution in PBS. The brain was then extracted and kept in 4% PFA solution overnight and moved to PBS. After storing the brain in 4 °C PBS overnight, the 50 µm coronal brain slices were prepared with vibrating blade microtome (Leica VT1000S). The permeabilization was done on the slices for 30 min at room temperature with 0.3% v/v Triton X-100 solution in PBS, and the media was changed to 0.3% v/v Triton X-100 and normal donkey serum 5% v/v blocking serum solution in PBS for 1h blocking on the orbital shaker. In the case using mouse primary antibody, the brain slices were additionally incubated in anti-mouse IgG (H+L) Fab fragment antibody (0.1 mg ml^-1^, Jackson ImmunoResearch 115-007-003) for 1 hour at room temperature. After PBS washing three times, the media was changed to first antibody solution for overnight incubation at 4 °C. Following PBS washing three times, at room temperature in dark condition, the brain slices were immersed in a secondary antibody solution for 2 hour on the orbital shaker. After three times PBS washing, 4′,6-diamidino-2-phenylindole (DAPI) staining was performed for 30 min. After PBS washing, the brain slices were transferred onto glass slides with mounting medium (Fluoromount G, Southern Biotech).

As primary and secondary antibody pairs, rabbit anti-c-Fos (1:500, Cell Signaling Technology, 2250s) paired with donkey anti-rabbit Alexa Fluor 488 (1:1,000, Invitrogen, A21206), rabbit anti-tyrosine hydroxylase antibody (1:500, Novus Biologicals NB300-109) paired with donkey anti-rabbit Alexa Fluor 647 (1:1,000, Invitrogen A31573), mouse SOD1 antibody (1:100, ThermoFisher MA1-105) and mouse mTOR antibody (1:100, ThermoFisher AHO1232) paired with donkey anti-mouse Alexa Fluor 488 (1:1,000, Invitrogen A21202), and three pairs of primary (goat anti-Iba1 antibody (1:500, Abcam ab107159), goat anti-GFAP antibody (1:1,000, Abcam ab53554), rabbit anti-CD68 antibody (1:250, Abcam ab125212)) and secondary (donkey anti-goat IgG (H + L), Alexa Fluor 633 (1:1,000, Fisher Scientific A-21082) and donkey anti-rabbit Alexa Fluor 488 (1:1,000, Invitrogen, A21206)) antibodies were used.

For imaging of the STN, coronal sections were collected at an anterior–posterior (AP) coordinate of −2.1±0.2 mm. For VTA and SNr analyses, coronal sections were collected at AP −3.1±0.2 mm. These coordinates were selected to ensure consistent sampling across animals.

To quantify TH-positive neurons, images were processed using a custom analysis pipeline (Supplementary Note 1). The noise-removal step was optimized to exclude dendritic structures prior to cell counting, thereby preventing the misidentification of somata located adjacent to dendrites as separate cells.

### Statistical analyses

Origin 2024b Pro was used for statistical assessments for following analysis; to compare c-Fos expression in the MEND-injected left and PBS-injected right subthalamic nucleus (STN) of naïve animals, data normality was assessed using the Shapiro–Wilk test, and a paired t-test was then conducted to evaluate statistical differences between the two regions. To compare the TH-positive ratio (left-to-right side of the brain) at 2 and 4 weeks after 6-OHDA injection, normality was assessed using the Shapiro–Wilk test, followed by a two-sample t-test for statistical analysis. For comparisons among Groups 1–5, normality was tested with the Shapiro–Wilk test. Statistical analysis was conducted using one-way ANOVA, followed by Tukey’s post-hoc test. For comparison among Groups 1-5 in cylinder, rotarod assays, and SLA, the normality of the dataset was assessed with the Shapiro–Wilk test, and for the dataset rejecting normality, the paired sample Wilcoxon signed ranks test was performed for statistical analysis. The statistical analysis for the other dataset was performed with a paired t-test. For quantitative comparison of TH, SOD, and mTOR staining, Shapiro–Wilk test was performed to assess normality of data distribution.

As some data reject normality, the statistical analysis was performed with the Kruskal-Wallis test followed by a post-hoc Dunn’s test. For quantitative comparison of the immune responses with Iba1, GFAP, and CD68, the Kolmogorov-Smirnov test was performed to test the normality of data distribution. As all datasets have a normal distribution, the statistical analysis was performed with One-way Anova followed by Tukey’s test.

For the violin plots of velocity and stride length, statistical analysis was performed using the Wilcoxon rank-sum test in Python (NumPy library). While some plots depict pre- and post-stimulation data from the same group, the gait analysis pipeline automatically filters out animals with minimal movement. As a result, the pre- and post-stimulation groups were treated as independent. The polar plots for the interlimb coordination analysis were obtained by using the circular definition of mean and standard deviation.

## Acknowlegements

This work was funded in part by the Director’s Pioneer Award from the National Institutes of Health and National Institute for Complementary and Integrative Health (DP1-AT011991), McGovern Institute for Brain Research at MIT, K. Lisa Yang and Hock E. Tan Center for Molecular Therapeutics at MIT, K. Lisa Yang Brain-Body Center at MIT, and Pappudu Sriram and Dr. Rajesh Venkataramani Parkinson’s Disease Fund at MIT. Y.J.K. is a recipient of the Mathworks Fellowship. E.F. is a recipient of the National Science Foundation Graduate Research Fellowship. S.S. is the recipient of the McGovern Institute 25^th^ Anniversary Fellowship. N.B. is a recipient of the Human Frontier Science Program: Long-term Postdoctoral Fellowship.

## Author contributions

Y.J.K. and P.A. designed the study. Y.J.K. synthesized and characterized the materials and performed all in vitro experiments. Y.J.K., A.D.C., N.S. developed the gait analysis pipeline. Y.J.K. and E.V.P. performed behavioral assays and immunohistology. Y.J.K. and S.S. performed the surgeries. E.F. fabricated the fiber used for the electrode-based DBS. R.M. assisted with the setup for the gait-recording assay. N.B. contributed to the analyses and figures. R.L. contributed to figure preparation. All authors contributed to the writing of the manuscript.

## Code availability

The code for GaitPattern is available via the Github repository at https://gitshare.me/repo/bedb9ac3-ea8a-4200-b352-cdd49e62ce21

## Data availability

All data supporting the conclusions of the manuscript are available within the manuscript or the Supplementary Information. Small quantities of physical samples of the magnetoelectric nanodiscs are available from the corresponding authors upon reasonable request.

## Supplementary Information

### Supplementary Note 1

MATLAB code to quantify cells expressing tyrosine hydroxylase (TH+)

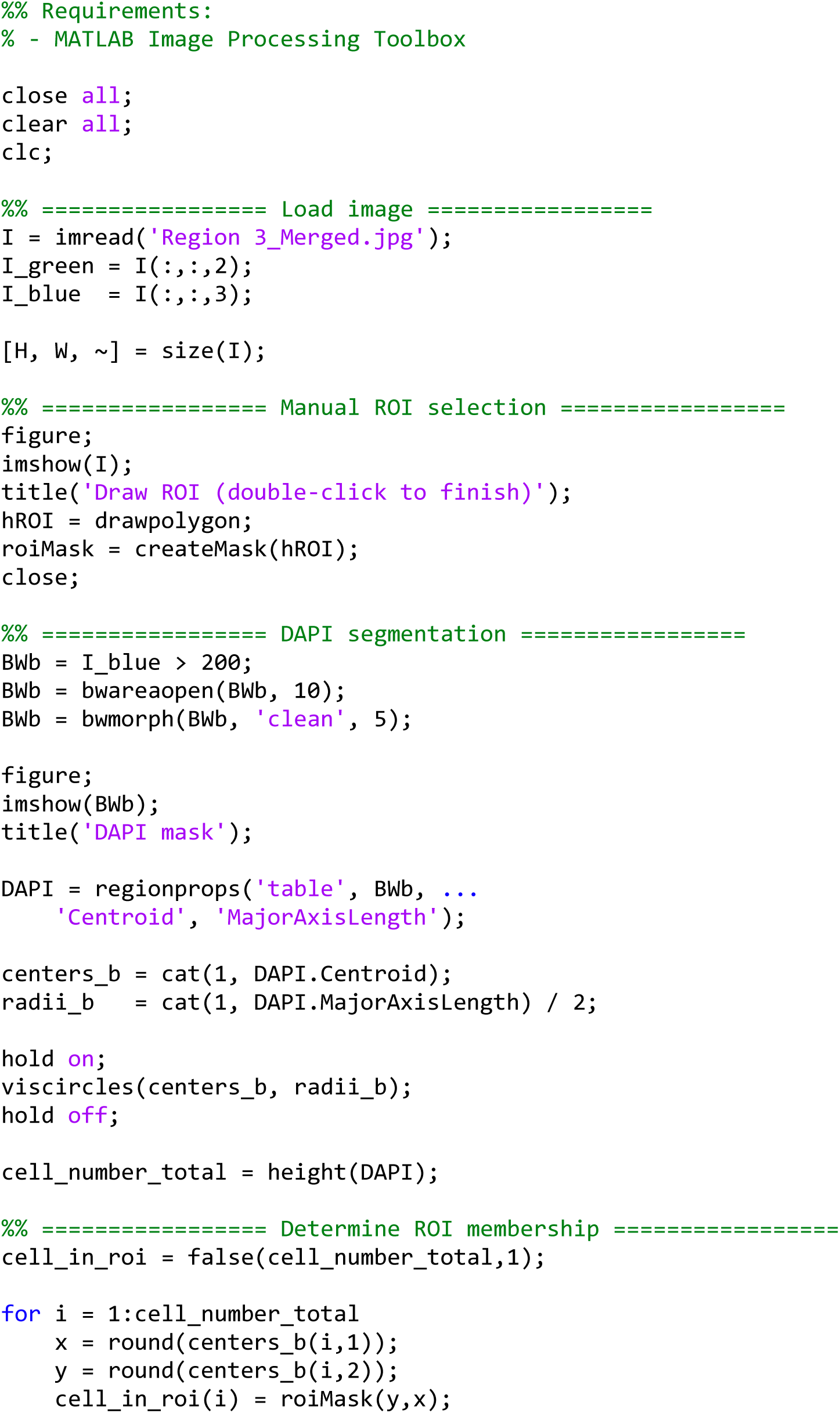
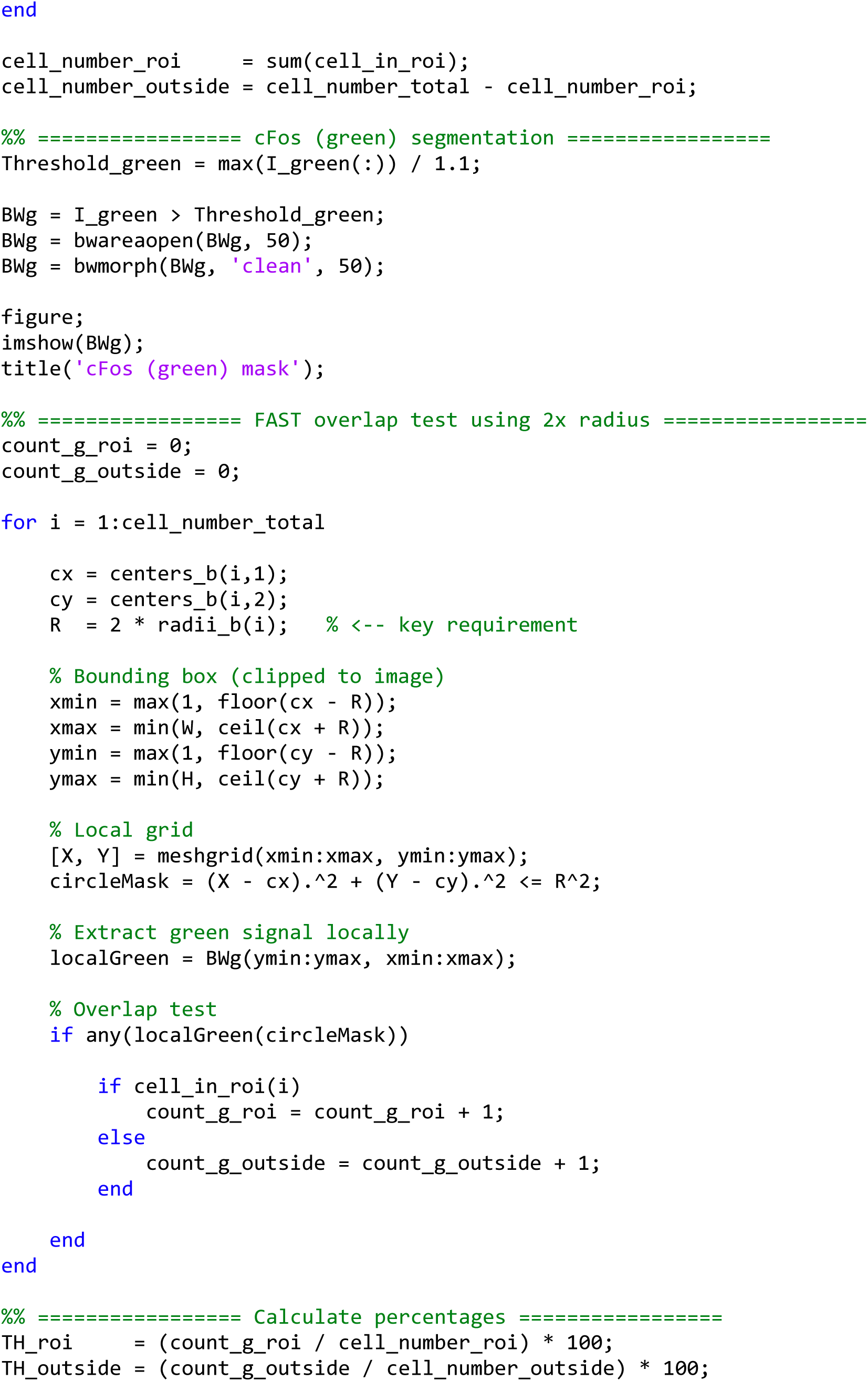
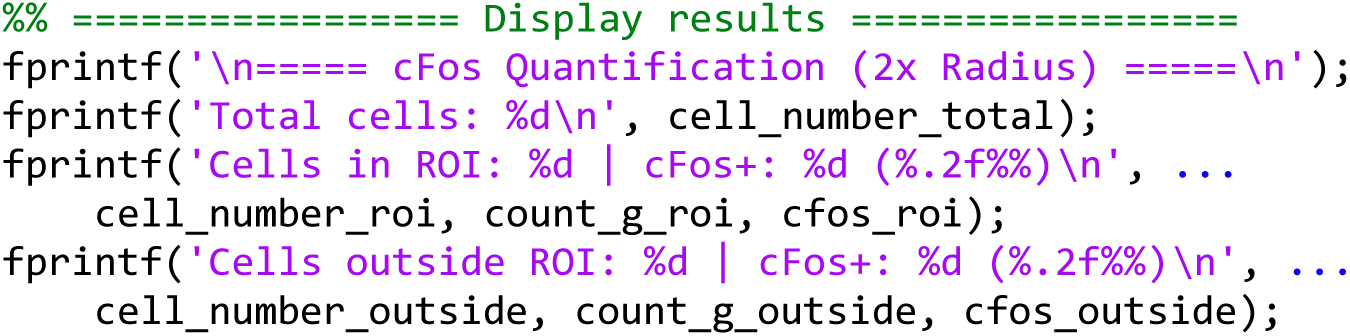

**Supplementary Fig. 1.**
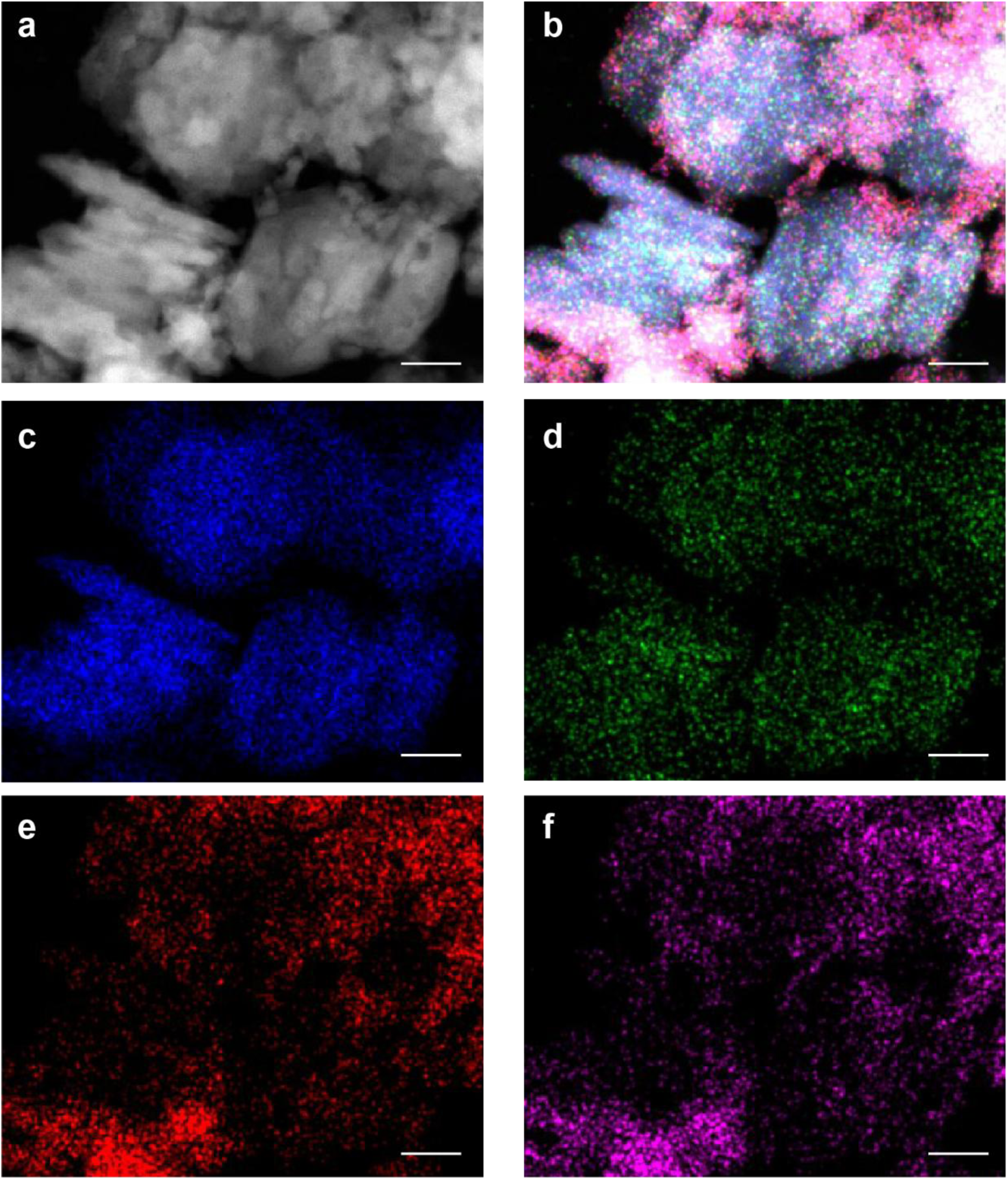
MEND images and elemental mapping. **a** High-angle annular dark-field scanning transmission electron microscopy (HAADF-STEM) image of MENDs. **b** Overlay of Energy Dispersive X-ray Spectroscopy (EDS) mapping on HAADF-STEM image. EDS mapping was performed for **c** Fe, **d** Co, **e** Ba, and **f** Ti. Scale bars = 100 nm.

**Supplementary Fig. 2.**
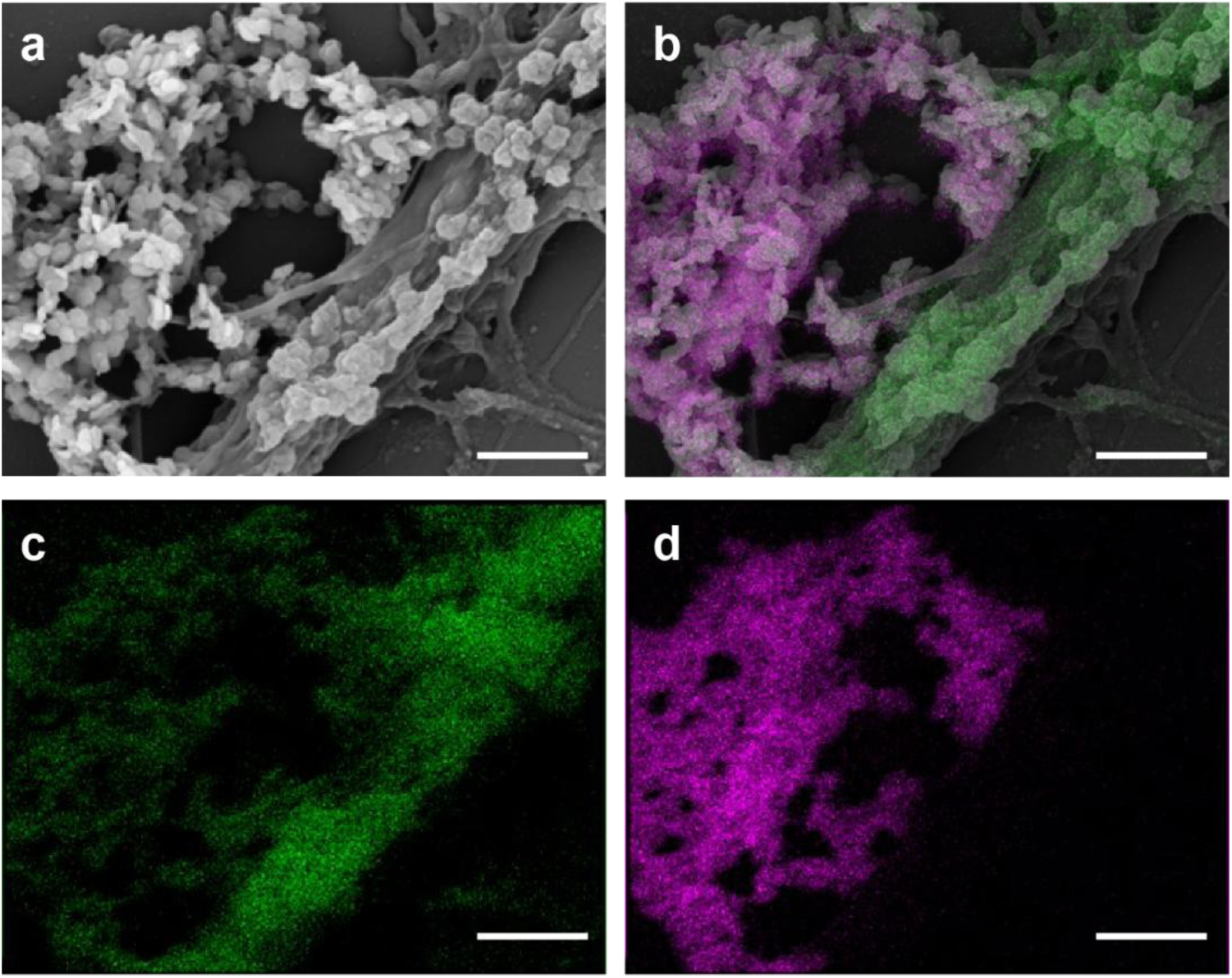
Neurons decorated with MENDs. **a** Scanning electron microscope (SEM) image of the neurons decorated with MENDs. **b** Overlay of elemental mapping on the SEM image. Elemental mapping was performed for **c** C, implying neurons, and **d** Fe, indicating presence of MENDs. Scale bars = 1 µm.

**Supplementary Fig. 3.**
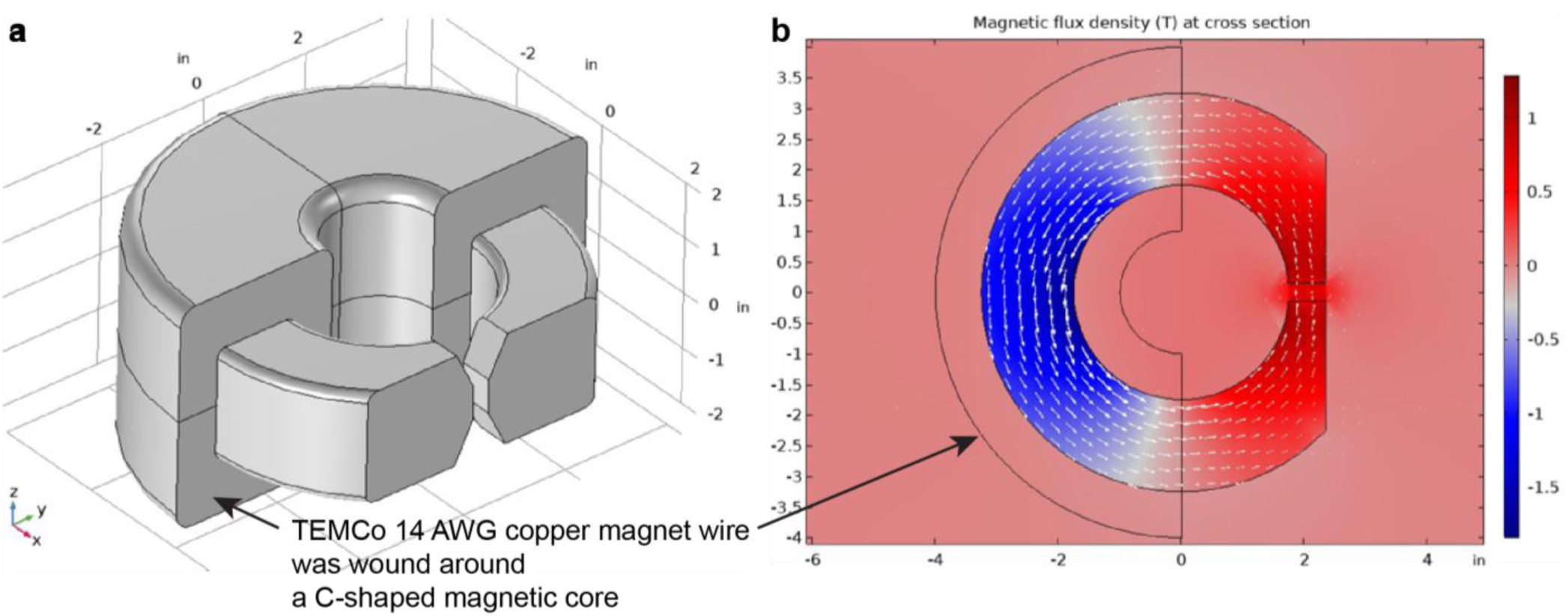
Electromagnet coil for ME coefficient measurement and calcium imaging. **a** Schematic illustration of the C-shaped magnetic core wound by 14 AWG copper wire. **b** Top view of the magnetic flux density from the coil simulated via COMSOL.

**Supplementary Fig. 4.**
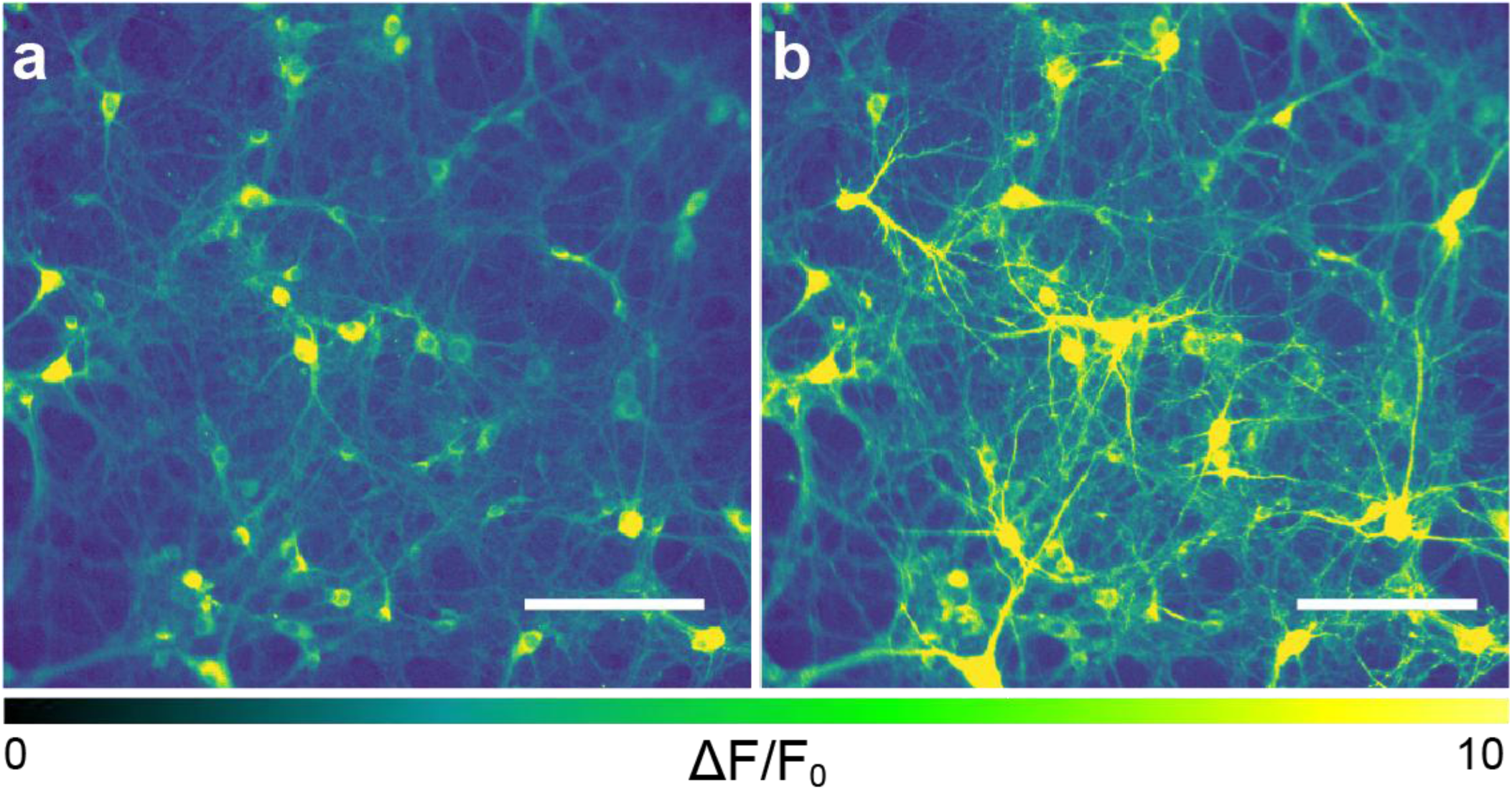
Ca^2+^ ion concentration transients in response to MEND-mediated stimulation. The relative GCaMP6s fluorescence change (Δ*F*/*F*_0_) in hippocampal neurons decorated with MENDs **a** before and **b** after MF application (2 s, OMF 220 mT; AMF 100 Hz, 10 mT). Scale bars = 150 µm.

**Supplementary Fig. 5.**
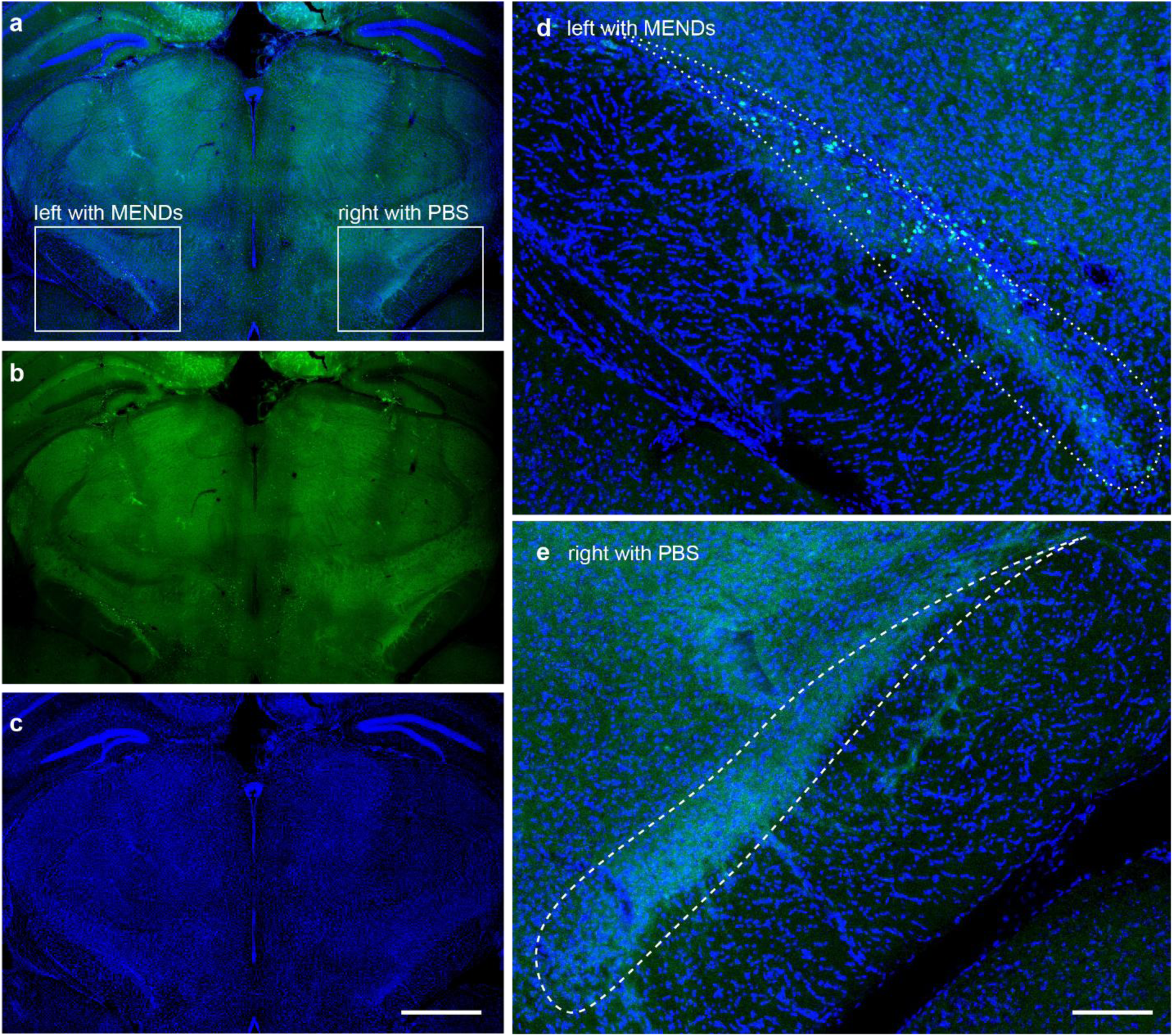
Image of the brain slice including the STN injected with MENDs in the left side and PBS in the right side. The injected MEND bolus is not obtainable, so spatial specificity of EMDNs can be implicated by the c-Fos expression after the exposure to the magnetic field. **a-c** the whole slice of the brain having injection of MENDs in the left STN and PBS in the right STN: **a** overlay of **b** c-Fos and **c** DAPI. Scale bar = 1 mm. Magnified view of **d** left and **e** right STN. Blue is DAPI, and green is c-Fos. Scale bar = 100 µm.

**Supplementary Fig. 6.**
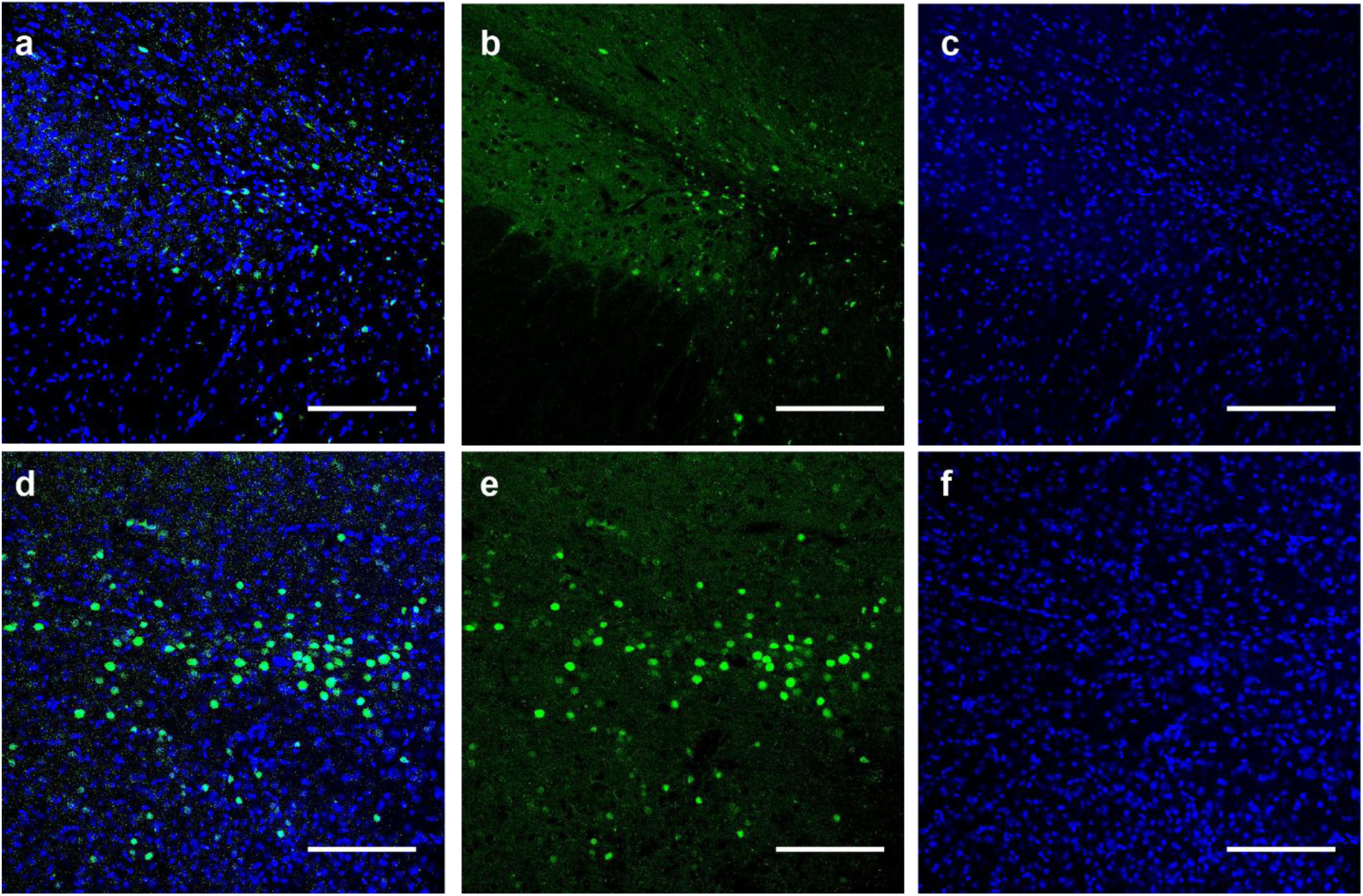
c-Fos expression in response to MEND-mediated stimulation. **a** Overlay of **b** c-Fos and **c** DAPI in the right STN, where PBS was injected 1 week prior to the application of magnetic field. **d** Overlay of **e** c-Fos and **f** DAPI in the vicinity of MENDs injected into the right STN of the same animal. Scale bar = 100 µm.

**Supplementary Fig. 7.**
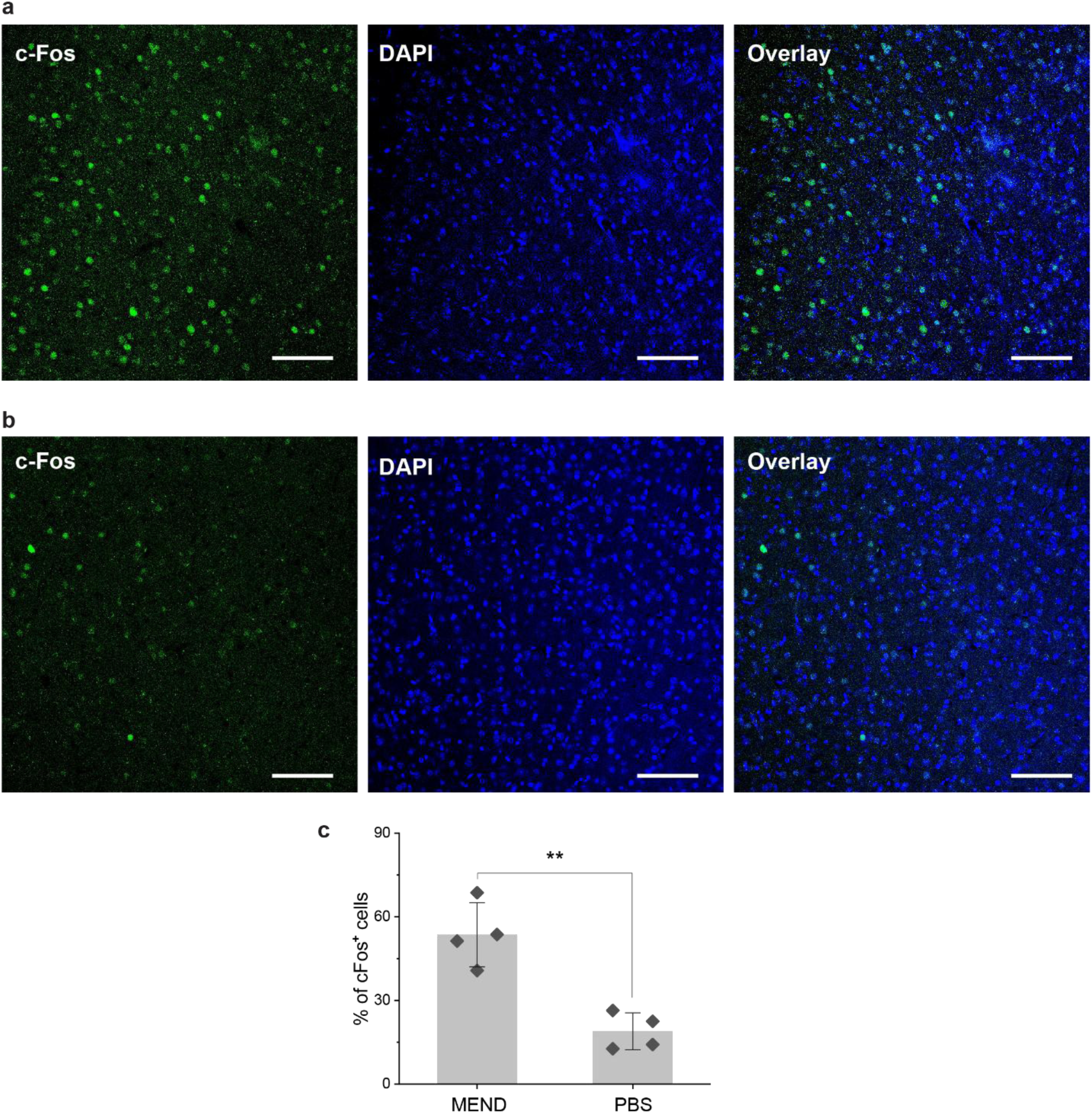
c-Fos expression in motor cortex. Immunofluorescence analysis for c-Fos expression on the motor cortex of the mice having **a** MENDs or **b** PBS in the left STN after the exposure to the magnetic field for calcium imaging. Scale bars = 100 µm. **c** Quantitative analysis on the number of c-Fos-positive cells out of number of DAPI cells in the motor cortex. Bars and error bars represent mean ± standard deviation (S.D.). As the data is fulfilling normal distribution, we performed two-sample t-test; p = 0.00198, t = 5.22, DF = 6.

**Supplementary Fig. 8.**
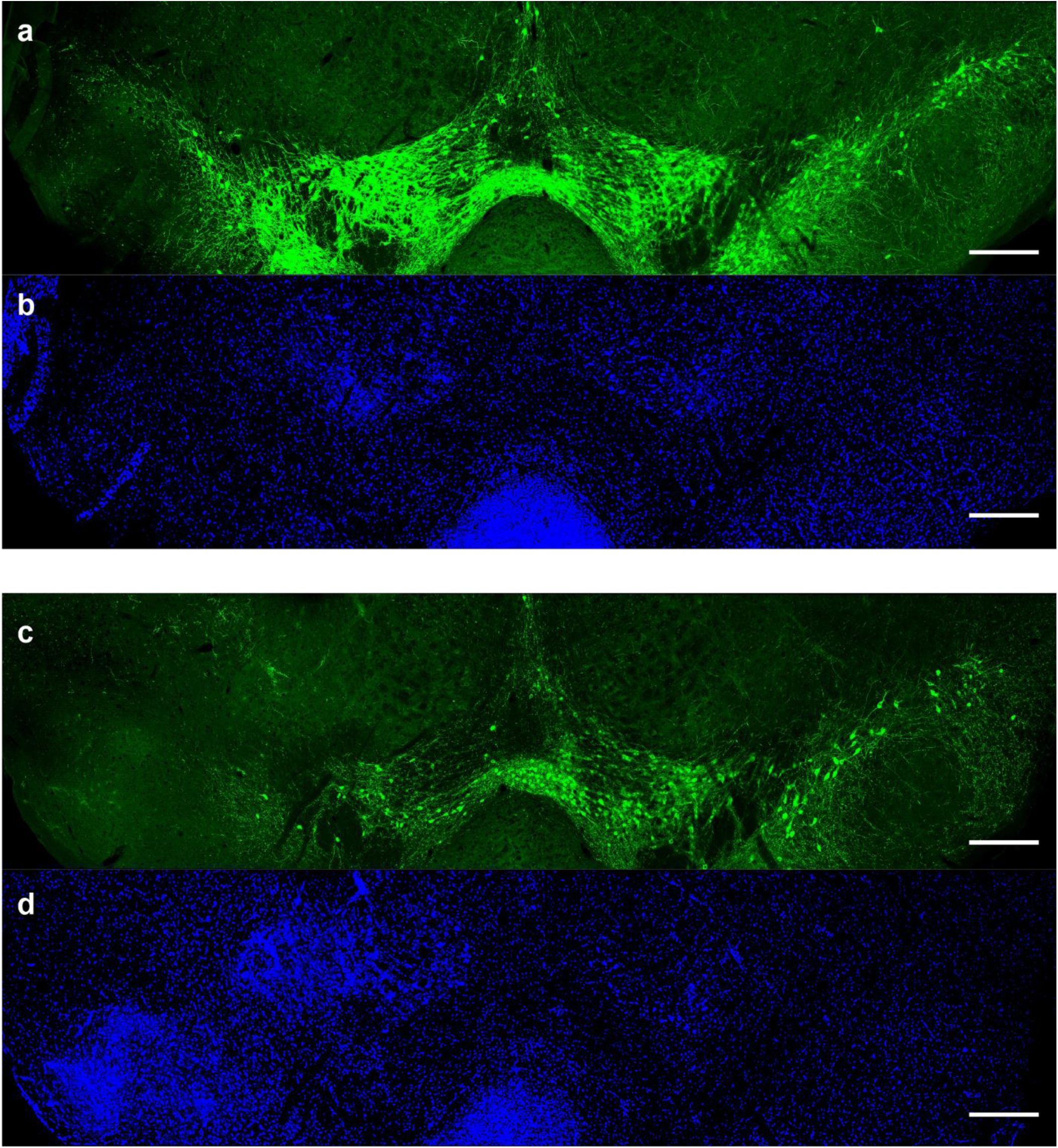
Degeneration of dopaminergic neurons in SNpc. **a** TH staining and **b** DAPI staining on the brain 2 weeks after the injection of 6-OHDA in left MFB. **c** TH staining and **d** DAPI staining on the brain 4 weeks after the injection of 6-OHDA in left MFB. Scale bars = 300 µm.

**Supplementary Fig. 9.**
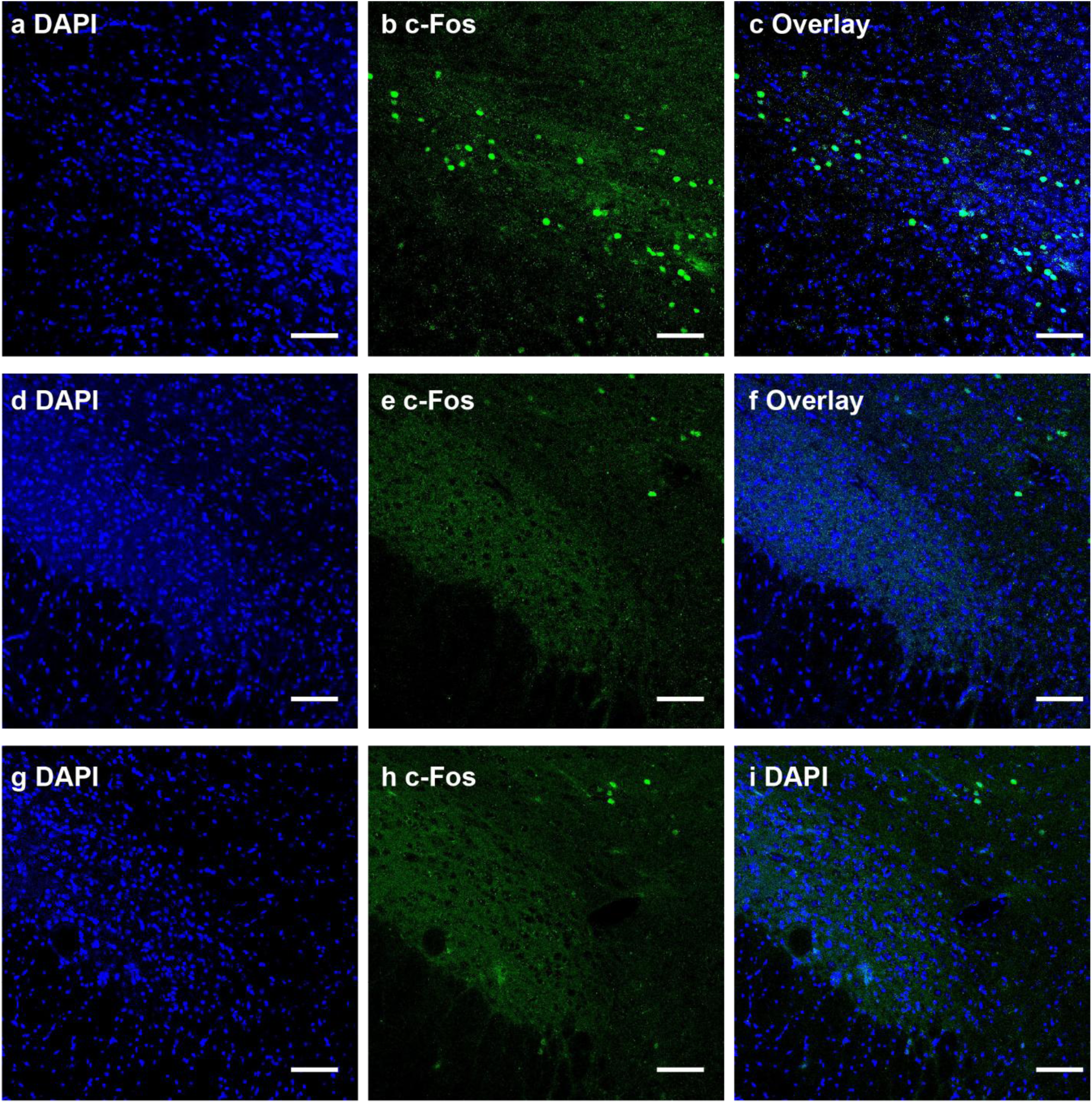
c-Fos expression in the STN. a-c. STN injected with MENDs, **d-f** STN injected with PBS, and **g-i** STN of a naive mouse. Fluorescent images of blue channel (**a,d,g** - DAPI), green channel (**b,e,h,** - c-Fos), and their overlay (**c,f,i**). Scale bar, 100 µm.

**Supplementary Fig. 10.**
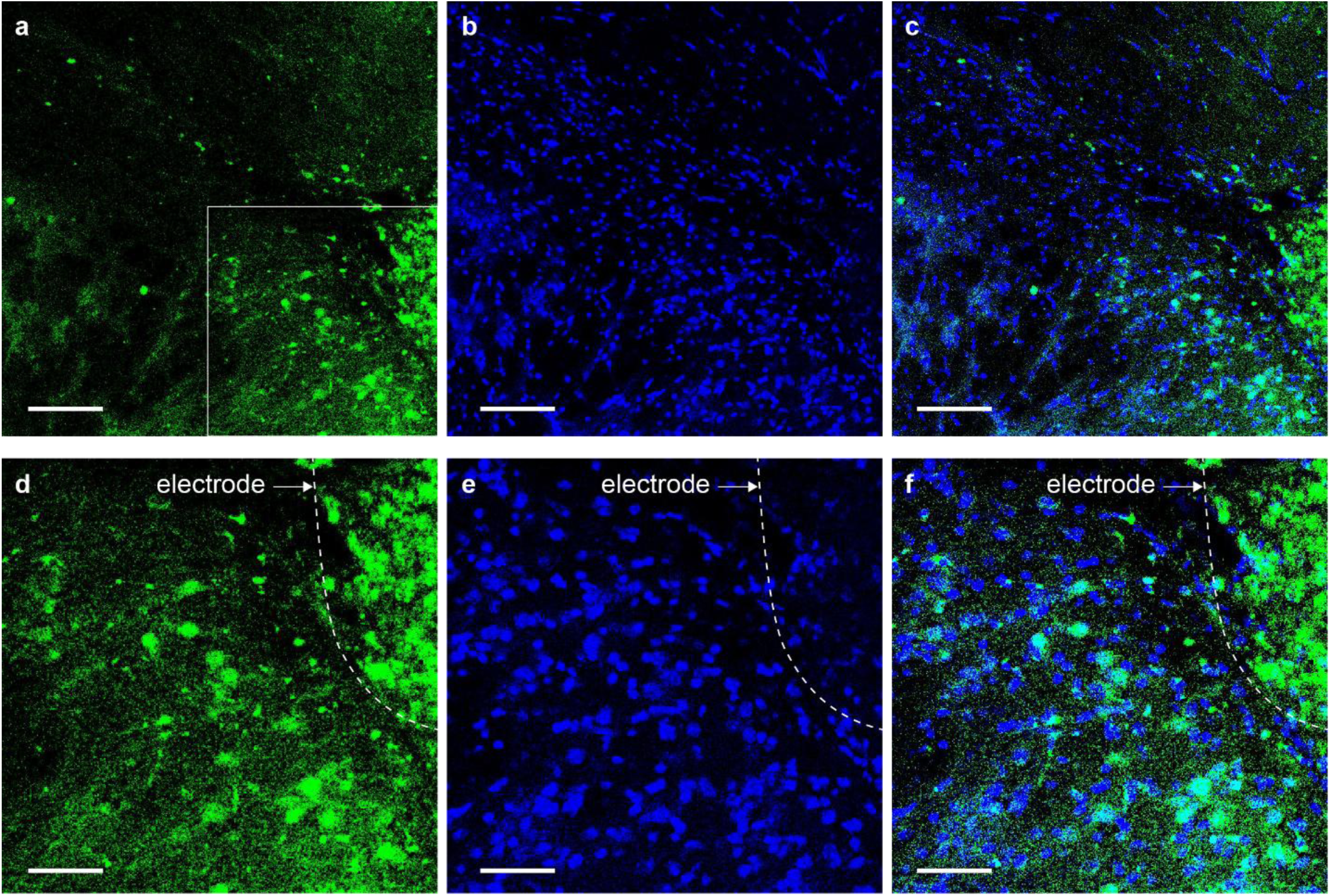
Magnified view of the STN after stimulation with an implanted electrode. a-c. Fluorescence images of **a** green channel (c-Fos), **b** blue channel (DAPI), and **c** their overlay. Scale bars = 100 µm. **d-f** Magnified views of the area indicated as white square in panel **a**; **d** green channel, **e** blue channel, and **f** their overlay. Scale bars = 25 µm.

**Supplementary Fig. 11.**
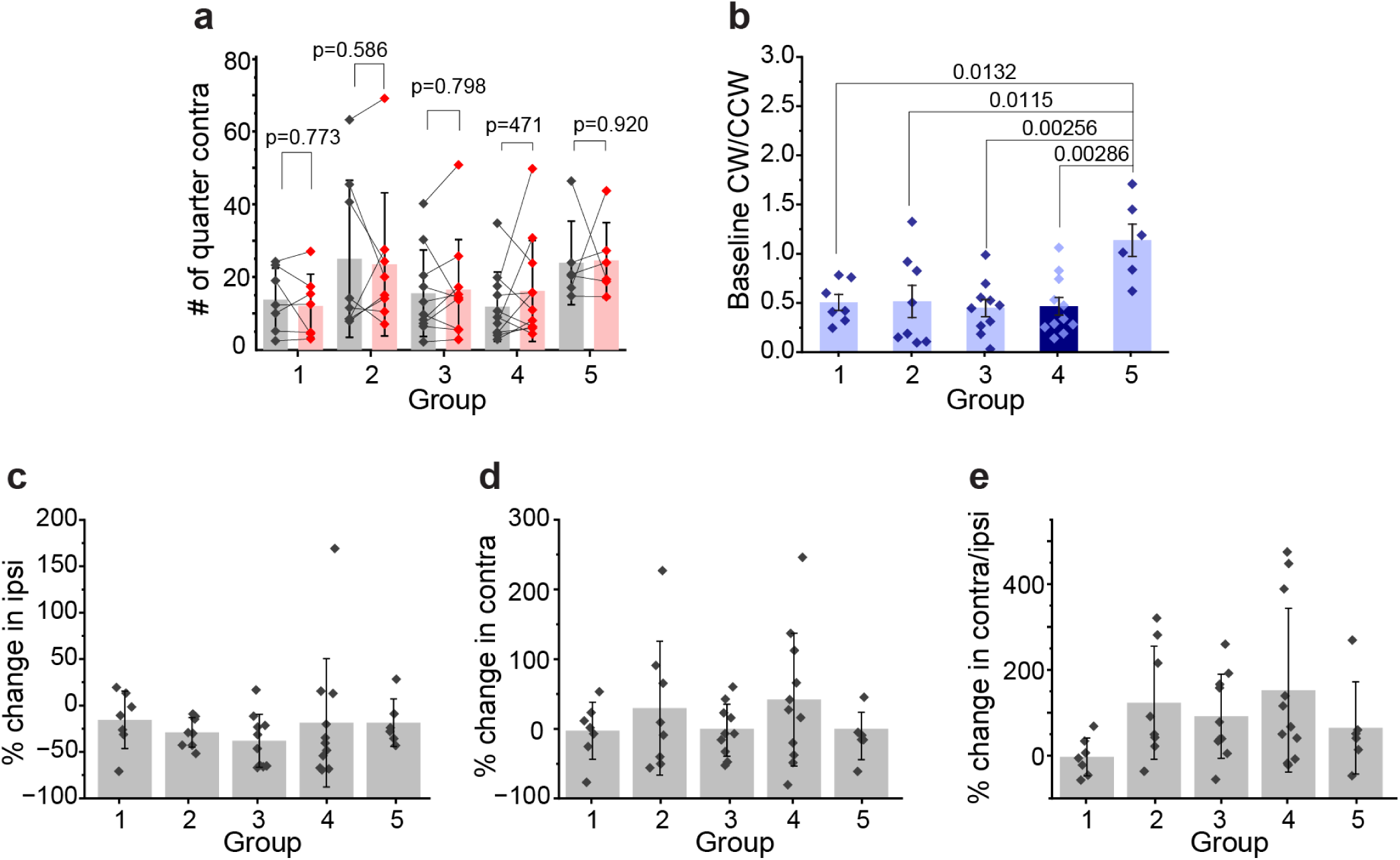
Cylinder test. **a** Number of quarter contralateral turns in the cylinder test. The statistical analysis was performed with two-way repeated measures ANOVA; (**f**) F_Group_(4,37)=0.834, P_Group_=0.519, F_Stim_(1,37)=0.0325, P_Stim_=0.864, F_Interaction_(4,37)=0.374, P_Interaction_=0.824, and Holm-Bonferroni post-hoc test confirmed pairwise pre vs post differences Group 1: p=0.773, Group 2: p=0.586, Group 3: p=0.798, Group 4: p=0.471, Group 5: p=0.920. **b** Number ratio of the contralateral turns to that of ipsilateral turns in the cylinder test before the stimulation on Day 1 of Test #1, corresponding to the baseline. The normality of the dataset was confirmed with the Shapiro–Wilk test, and the statistical analysis was performed with One-way ANOVA followed by Tukey’s test; F(4,37)=5.05, p=0.00238. **c-e** Inter-group comparison of the changes following the stimulation in **c** ipsilateral turns, **d** contralateral turns, and **e** the ratio of contralateral turns to the ipsilateral turns. The Shapiro–Wilk test was performed to test the normality of data distribution. The data in **c** fully follows the normal distribution, the statistical analysis was performed with One-way ANOVA followed by Tukey’s test; F(4,37)=0.94, p=0.45. As the data in plot **d** and **e** are not fully following the normal distribution, the statistical analysis was performed with the Kruskal-Wallis test followed by post-hoc Dunn’s test; **d** H=3.39, DF=4, p= 0.50; **e** H=7.21, DF=4, p= 0.1253. Bars and error bars represent mean ± S.D.

**Supplementary Fig. 12.**
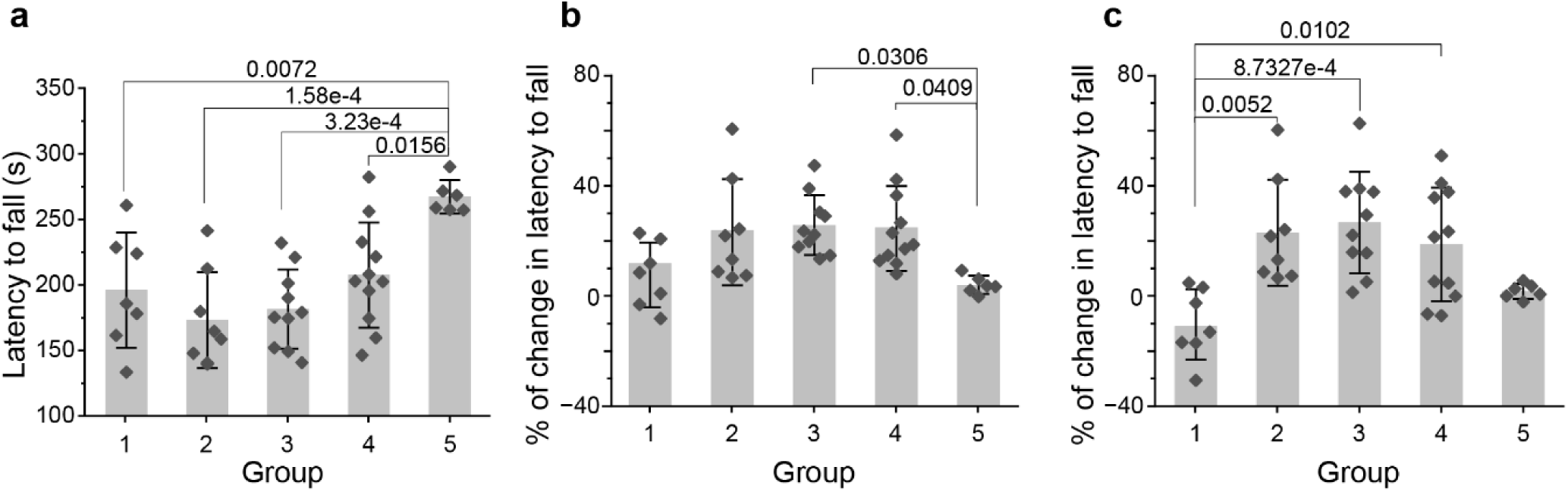
Rotarod test. **a** Latency to fall at Test #1 pre-stimulation. **b-c** Percent change in the latency to fall (100% × (Post-Pre)/Pre) at **b** Test #1 and **c** Test #2. The normality of the dataset was confirmed with the Shapiro–Wilk test, and the statistical analysis was performed with One-way ANOVA followed by Tukey’s test; (**a**) F=7.420, p=1.711×10^−4^, DF=4, (**b**) F(4,37)=4.241, p=6.03e-3, (**c**) F(4,37)=6.392, p=5.15e-4. Bars and error bars represent mean ± S.D..

**Supplementary Fig. 13.**
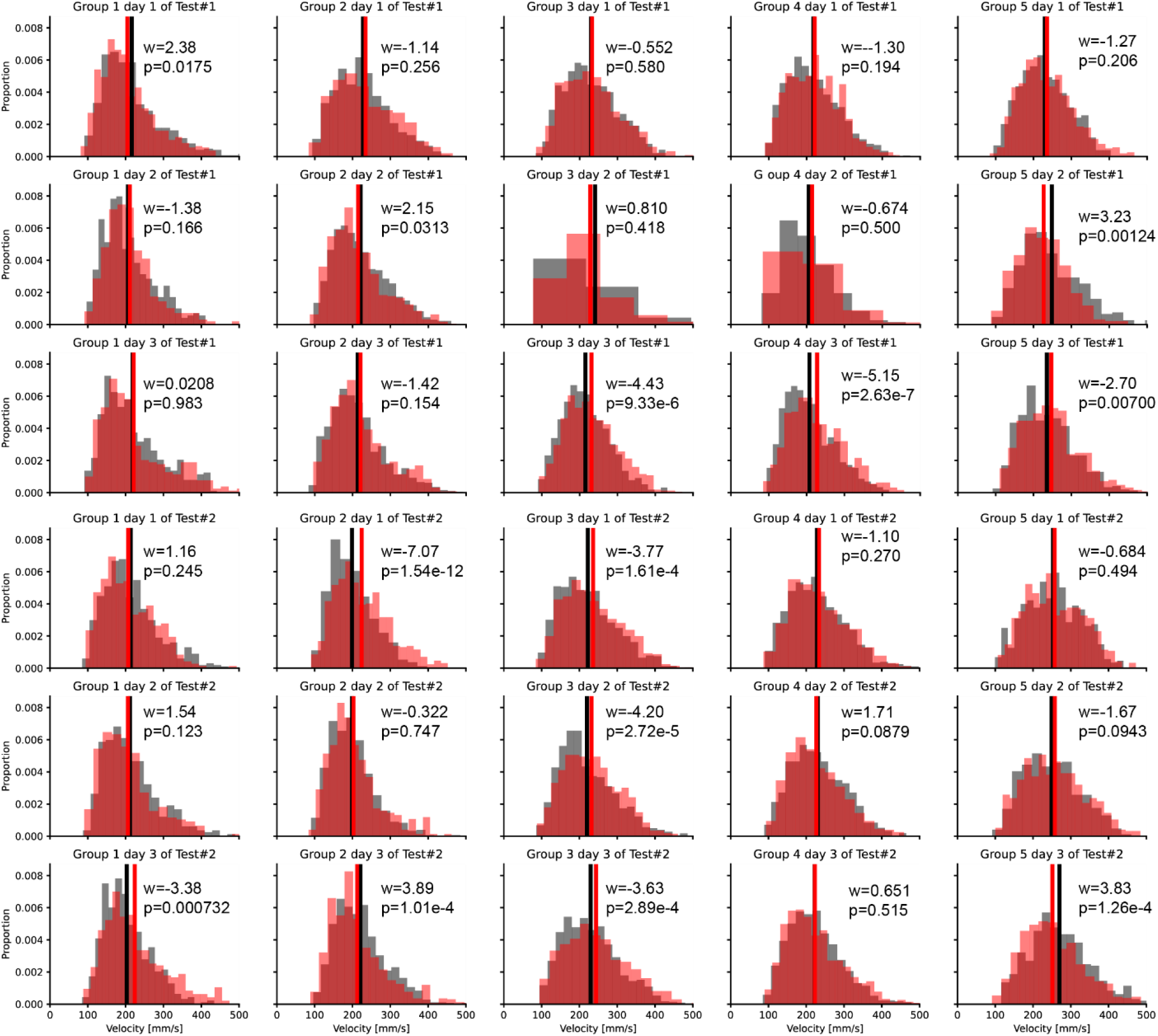
Movement velocity changes in response to the stimulation. Histograms of the movement velocity for each step of animals in Groups 1 to 5, presented left to right, from Day 1 of Test #1 (earliest time point) to Day 3 of Test #2 (latest time point), presented top to bottom. The statistical analysis was performed with the Wilcoxon Rank Sum Test.

**Supplementary Fig. 14.**
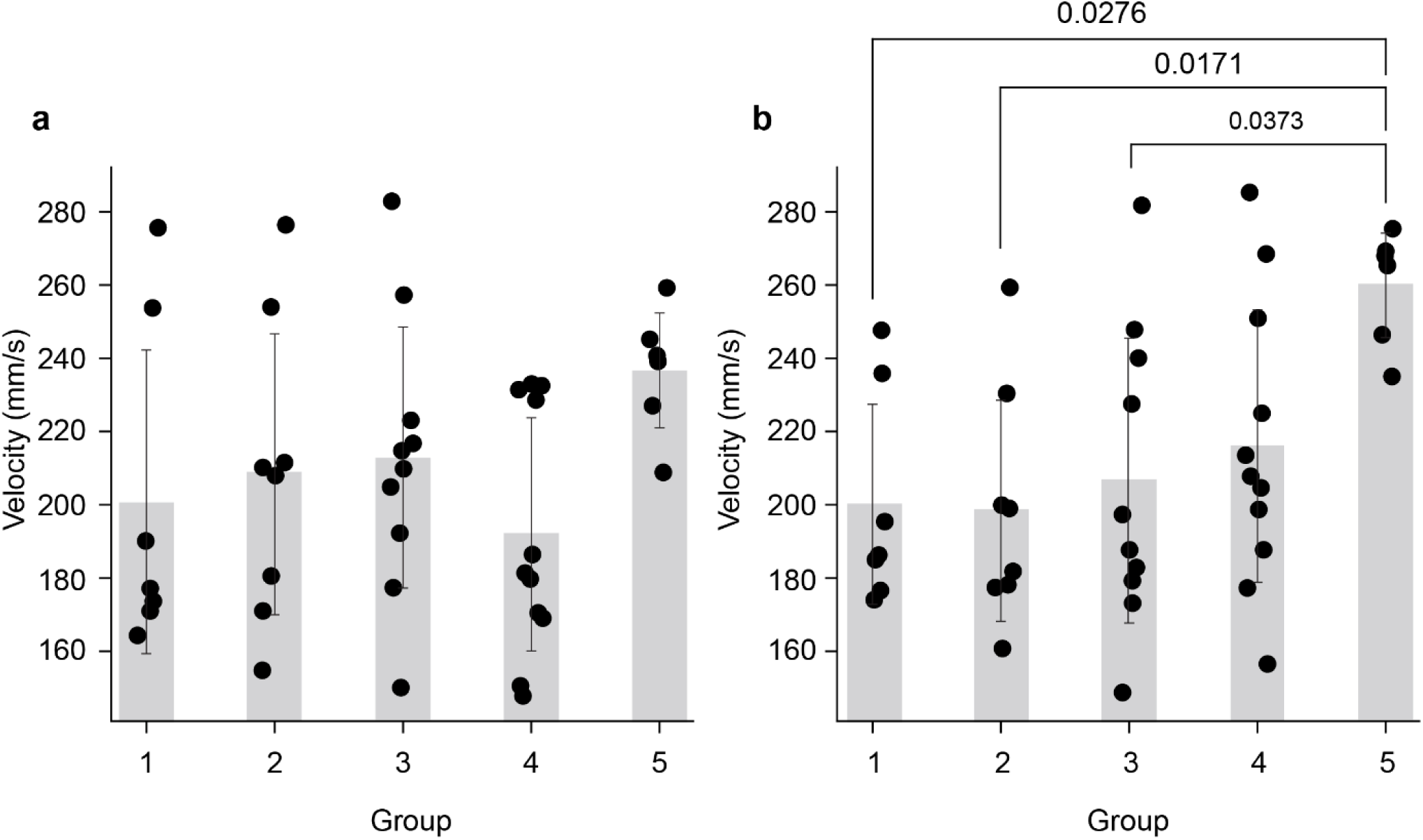
Inter-group comparison of velocity. Scatter plot of average velocity per animal at the pre-stimulation sessions over three days in (**A**) Test #1 and (**B**) Test #2. The statistical analysis was performed with One-way ANOVA followed by Tukey’s test; at Test #1 F(4,37)=1.58, p=0.200; at Test #2 F(4,37)=3.47, p=0.0167. Bars and error bars represent mean ± S.D.

**Supplementary Fig. 15.**
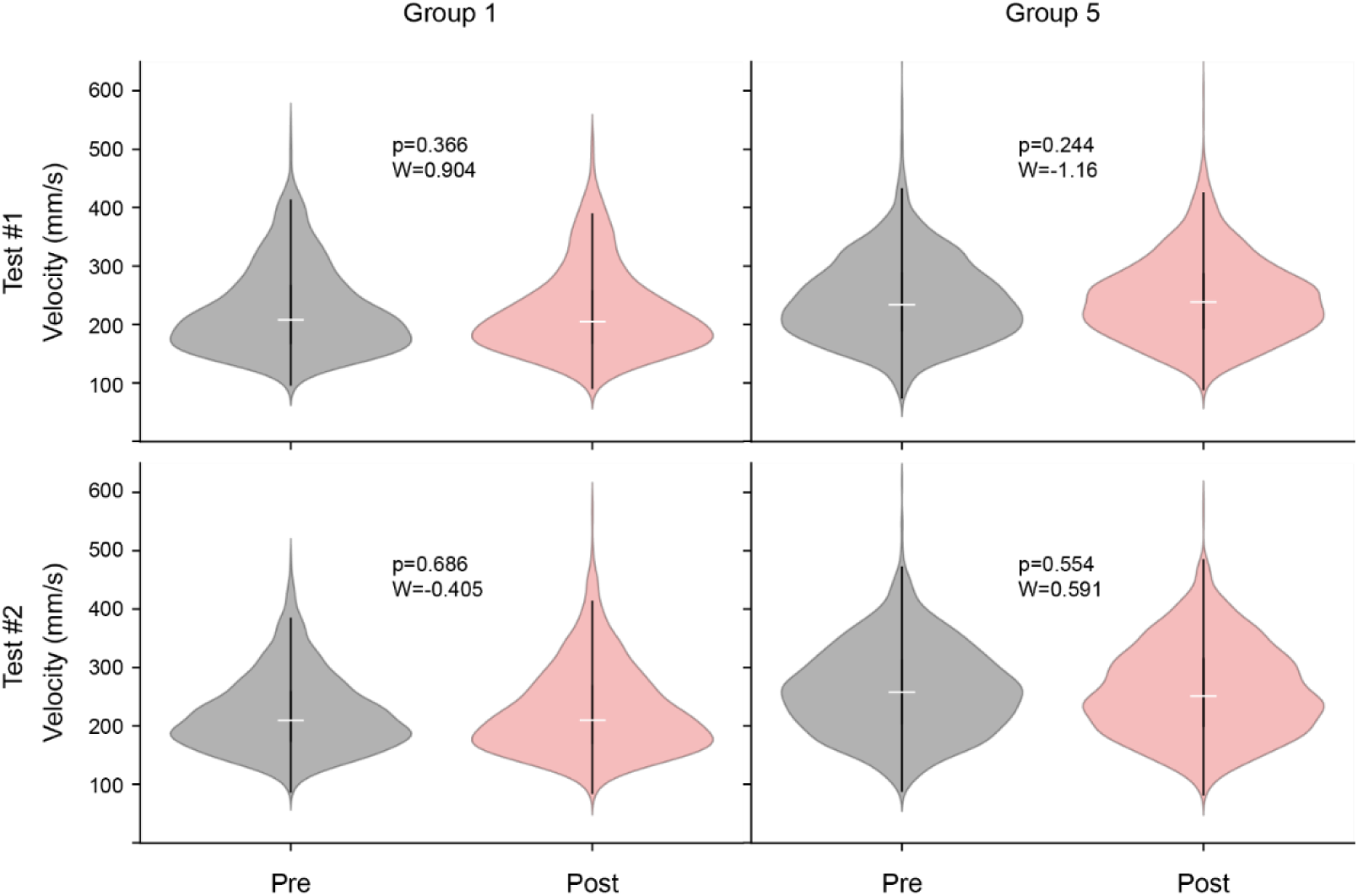
Velocity changes immediately after the stimulation for Groups 1 and 5. Pre-and Post-stimulation velocity of Group 1 and 5 (left and right) at Tests #1 and #2 (top and bottom). The statistical analysis was performed via Wilcoxon Rank Sum Test.

**Supplementary Fig. 16.**
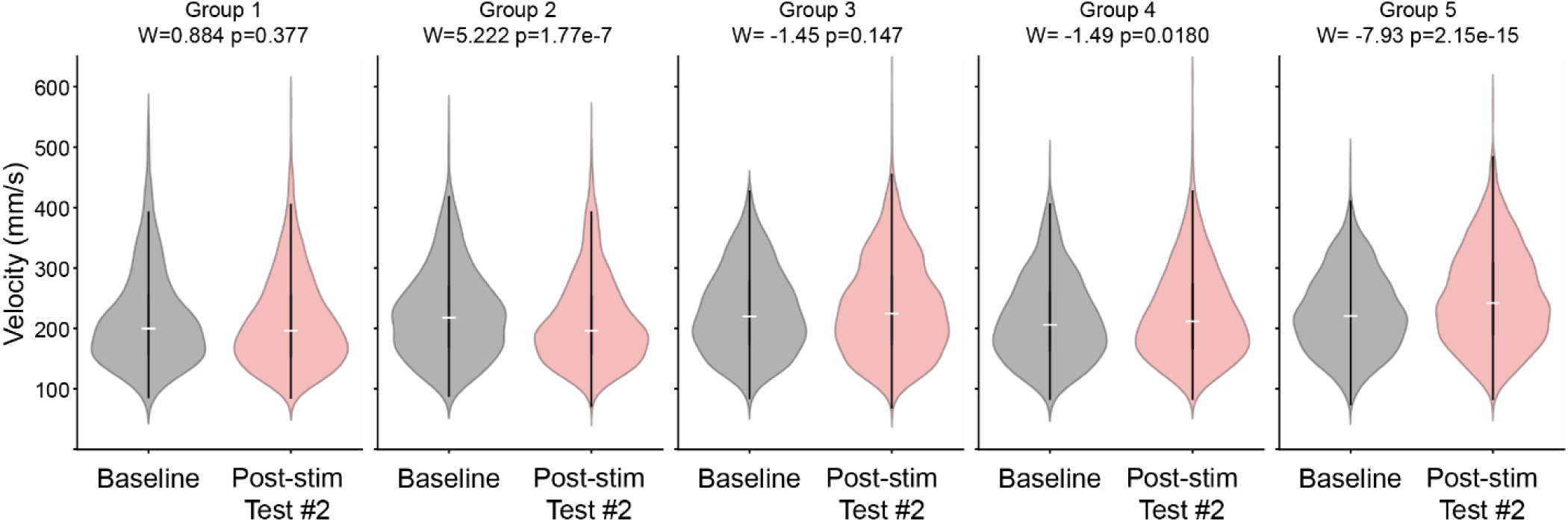
Velocity changes from baseline to Post-stimulation at Test #2. Baseline velocity was measured at Day 1 of Test #1 – the pre-stimulation session. The statistical analysis was performed via Wilcoxon Rank Sum Test.

**Supplementary Fig. 17.**
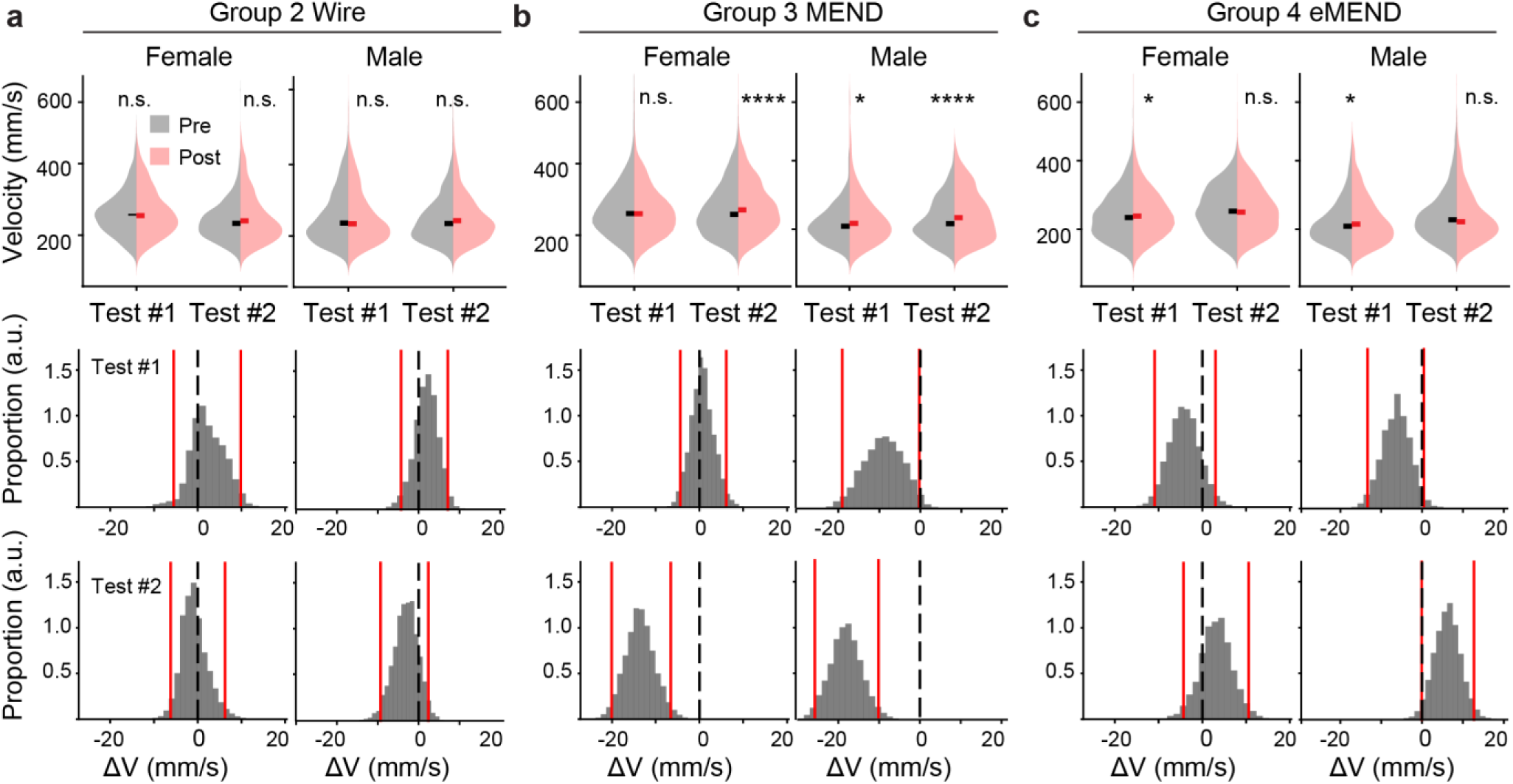
Female-male separate analysis. Velocity pre- and post-stimulation at Test #1 and Test #2 of **a** Group 2, **b** Group 3, and **c** Group 4. The statistical analysis for the histogram was performed via Wilcoxon rank sum test; Group 2 Female Test #1: W= −0.461, p=0.645, Test #2: W=-1.31, p=0.190, Male Test #1 W=-0.11, p=0.914 Test#2 W=-1.89, p=0.0585; Group 3 Female Test #1: W= .774, p=0.439, Test #2: W=-4.44, p=8.84×10^-6^, Male Test #1 W=-3.22, p=0.00129, Test#2 W=-4.443, p=9.55×10^-6^; Group 4 Female Test #1: W= −2.11, p=0.0346, Test #2: W=0.487, p=0.627, Male Test #1 W=-2.26, p=0.0241, Test#2 W=1.16, p=0.248. For Bootstrap resampling and regression analysis results; Group 2 Female ΔV_m, Pre-Post_ = 2.30 mm/s, CI = [-5.51, 9.84] mm/s, mm/s, p = 0.230 (Test #1), and ΔV_m, Pre-Post_ = −0.748 mm/s, CI = [-6.18, 6.26] mm/s, p = 5.53×10^-4^ (Test #2); Group 2 Male ΔV_m, Pre-Post_ = 1.90 mm/s, CI = [-4.30, 7.17] mm/s, p = 0.712 (Test #1), and ΔV_m, Pre-Post_ = −3.15 mm/s, CI = [-9.31, 2.40] mm/s, p = 1.15×10^-2^ (Test #2); Group 3 Female ΔV_m, Pre-Post_ = 0.667 mm/s, CI = [-4.37, 6.13] mm/s, p = 0.352 (Test #1), and ΔV_m, Pre-Post_ = −13.58 mm/s, CI = [−20.08, −6.56] mm/s, p = 6.68×10^-6^ (Test #2); Group 3 Male ΔV_m, Pre-Post_ = −9.05 mm/s, CI = [−19.024, −0.235] mm/s, p = 6.87×10^-7^ (Test #1), and ΔV_m, Pre-Post_ = −18.2 mm/s, CI = [−25.78, −10.14] mm/s, p = 9.02×10^-8^ (Test #2); Group 4 Female ΔV_m, Pre-Post_ = −4.13 mm/s, CI = [-10.9, 3.02] mm/s, p = 2.08×10^-2^ (Test #1), and ΔV_m, Pre-Post_ = 3.19 mm/s, CI = [-4.30, 10.59] mm/s, p = 0.865 (Test #2); Group 4 Male ΔV_m, Pre-Post_ = −6.27 mm/s, CI = [-13.32, 0.430] mm/s, p = 9.43×10^-4^ (Test #1), and ΔV_m, Pre-Post_ = 6.22 mm/s, CI = [-0.113, 12.721] mm/s, p = 0.994 (Test #2). In the bootstrap plots, black dashed line indicates 0, and red lines indicate the 95% CI.

**Supplementary Fig. 18.**
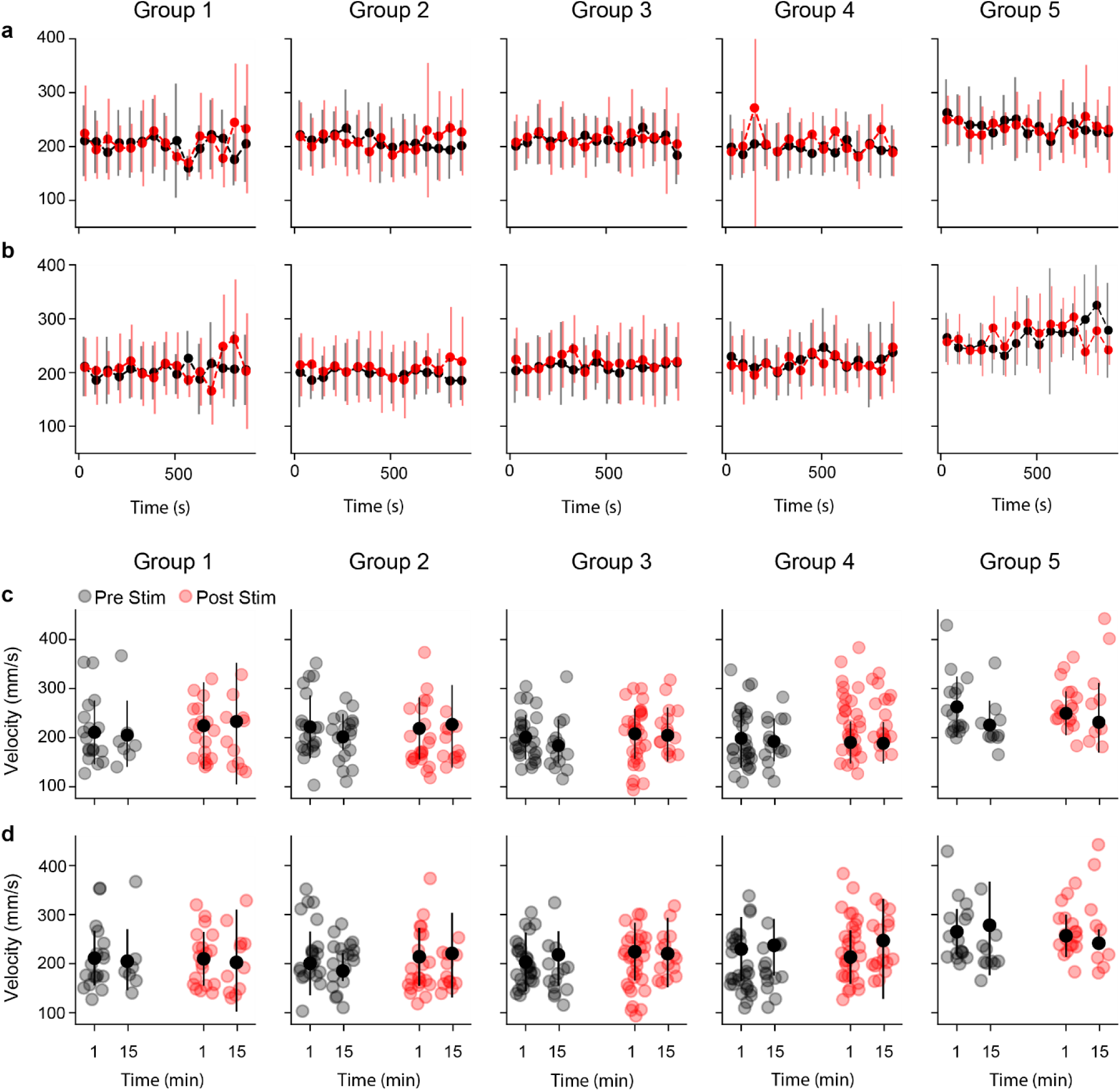
Movement velocity analyzed on per-minute basis. a-b. Average velocity in each group per minute at **a** Test#1 and **b** Test #2. **c-d** Quantitative comparison of the velocity between the first and the last 1 minute of the gait-analysis assay at **c** Test#1 and **d** Test #2. None of the pair of the first and last 1 minute have significant difference. Each data point is the average of each animal per day. Data is shown as the mean ± S.D., and black and red indicates pre- and post-stimulation, respectively.

**Supplementary Fig. 19.**
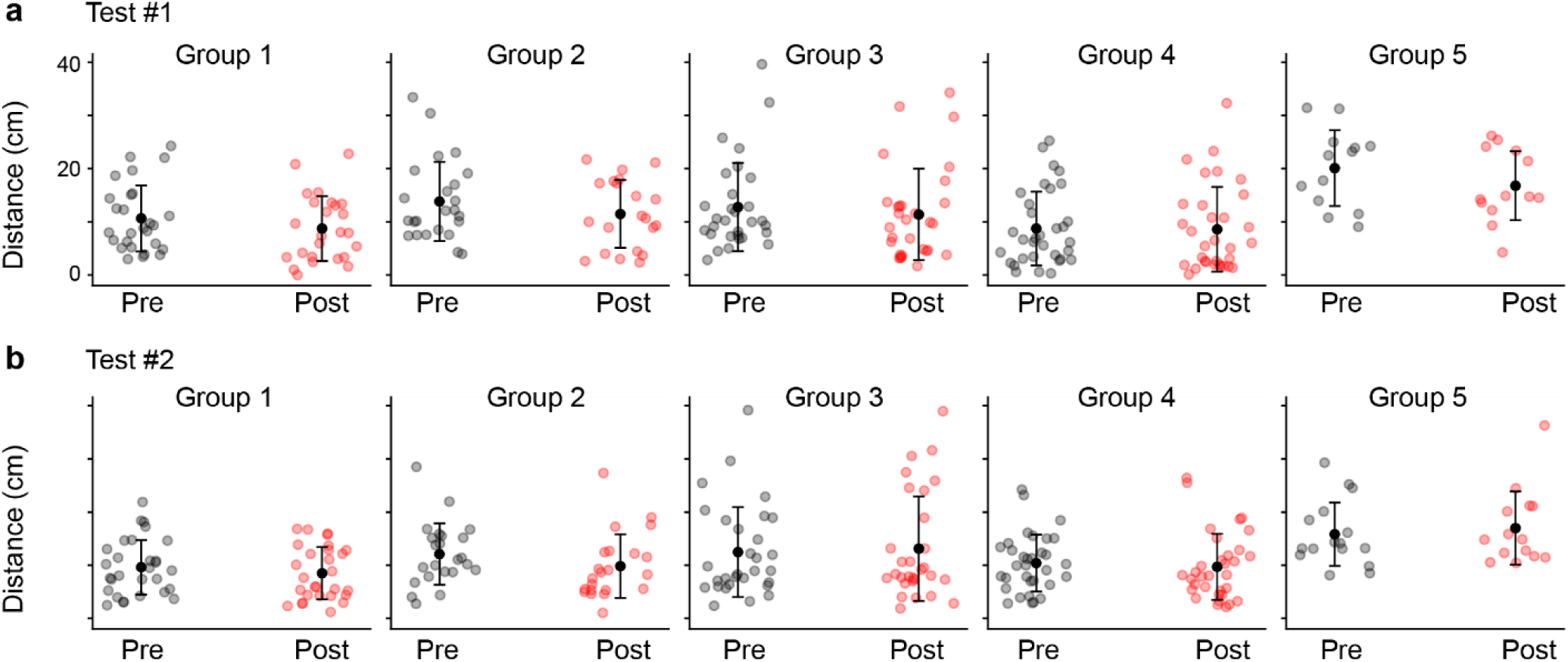
Movement distance. a-b. Movement distance of each animal per data (corresponding to each data point) shown with mean ± S.D. at **a** Test #1 and **b** Test#2. No statistical differences were identified.

**Supplementary Fig. 20.**
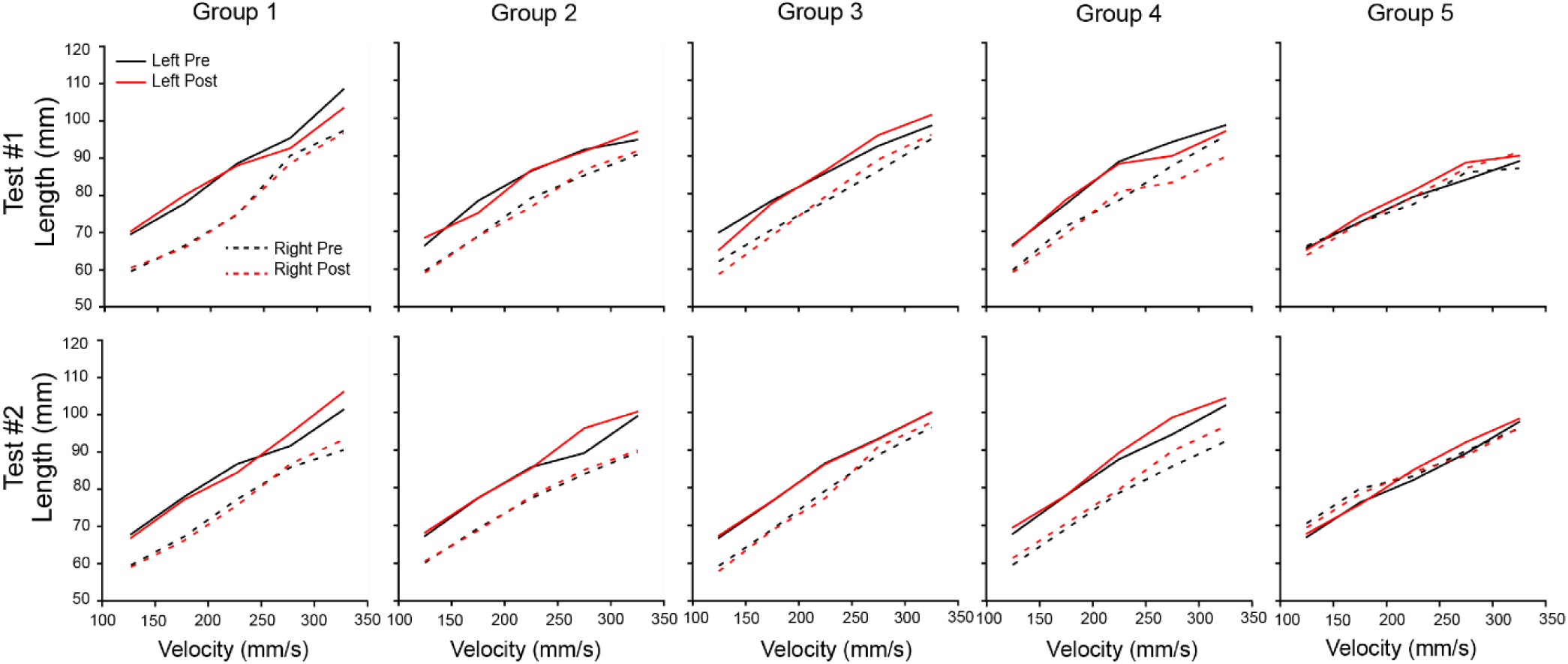
Step length. Step length for each group integrating all steps over the three days in each Test during pre- and post-stimulation sessions.

**Supplementary Fig. 21.**
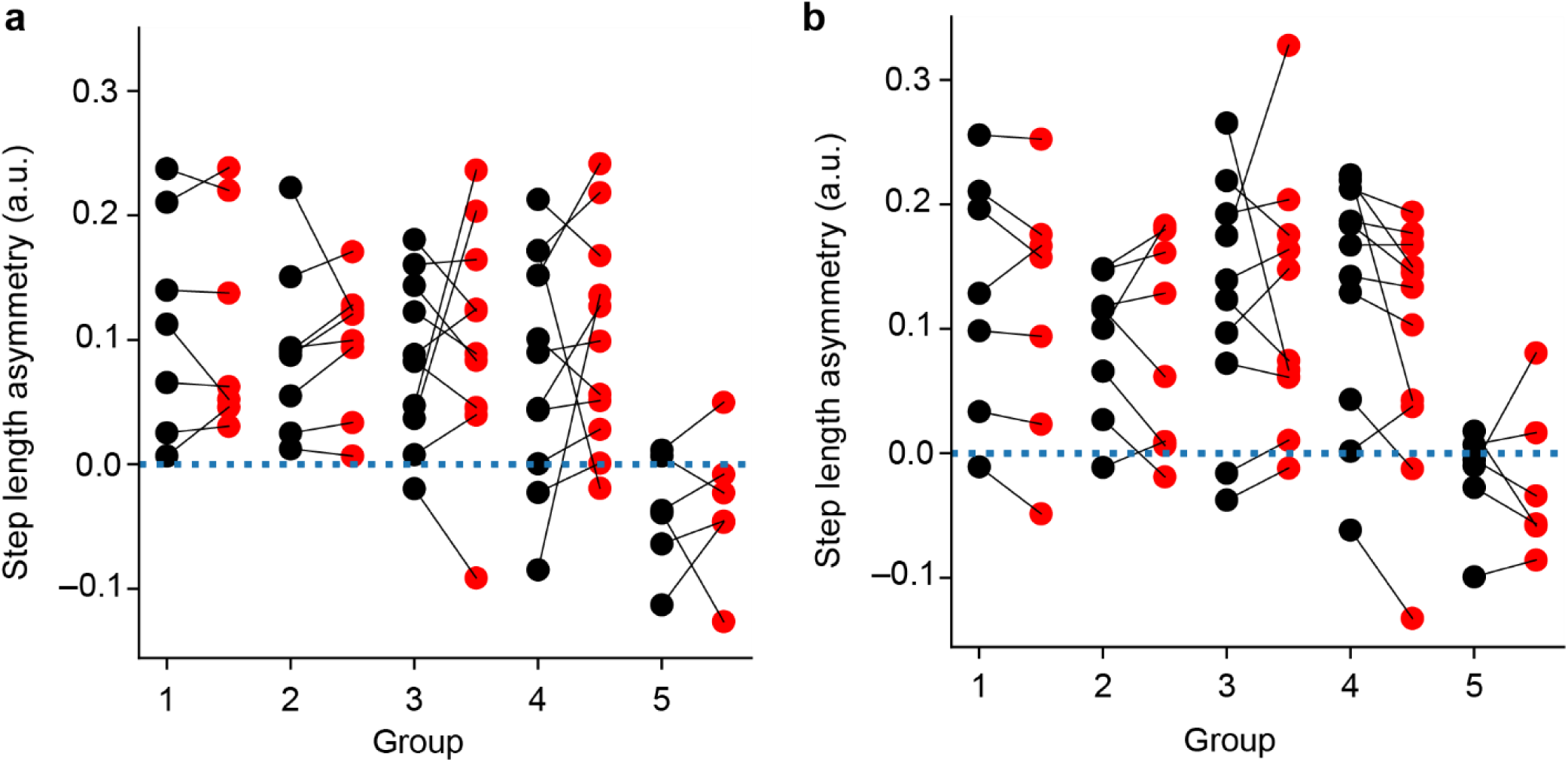
Step length asymmetry. Step length asymmetry (SLA) was calculated from each animal’s pre- (black) and post-stimulation (red) at **a** Test #1 and **b** Test #2. Paired t-test was performed for all groups to compare pre- and post-stimulation, and no significance was observed in any of the pairs.

**Supplementary Fig. 22.**
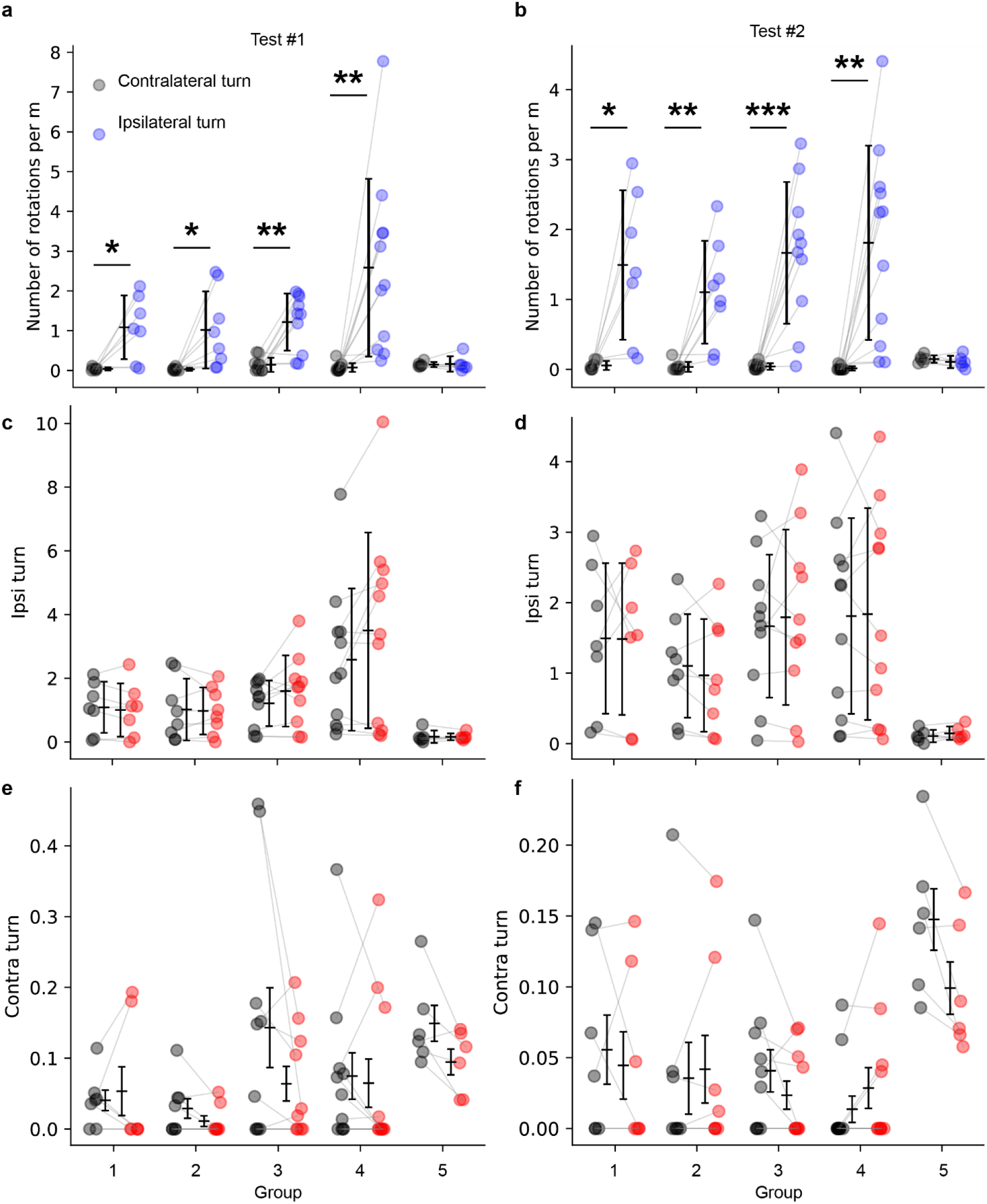
Rotations during the cat-walk assay. a-b. Number of contralateral and ipsilateral turns normalized by total movement distance pre-stimulation at **a** Test #1 and **b** Test #2. Each data point indicates each animal’s average across the three days in each test. Paired t-test was performed to compare the ipsi- and contra-lateral turns; **c-d** Number of ipsilateral turns normalized by total movement distance pre- and post-stimulation at **c** Test #1 and **d** Test #2. Each data point indicates each animal’s average across the three days in each test. Paired t-test was performed to compare the pre- and post-stimulation; **e-f** Number of ipsilateral turns normalized by total movement distance pre- and post-stimulation at **e** Test #1 and **f** Test #2. Each data point indicates each animal’s average across the three days in each test. Paired t-test was performed with Bonferroni correction to compare the pre- and post-stimulation **c-f**, none of the pair found to have significant differences. Markers and error bars represent S.D.

**Supplementary Fig. 23.**
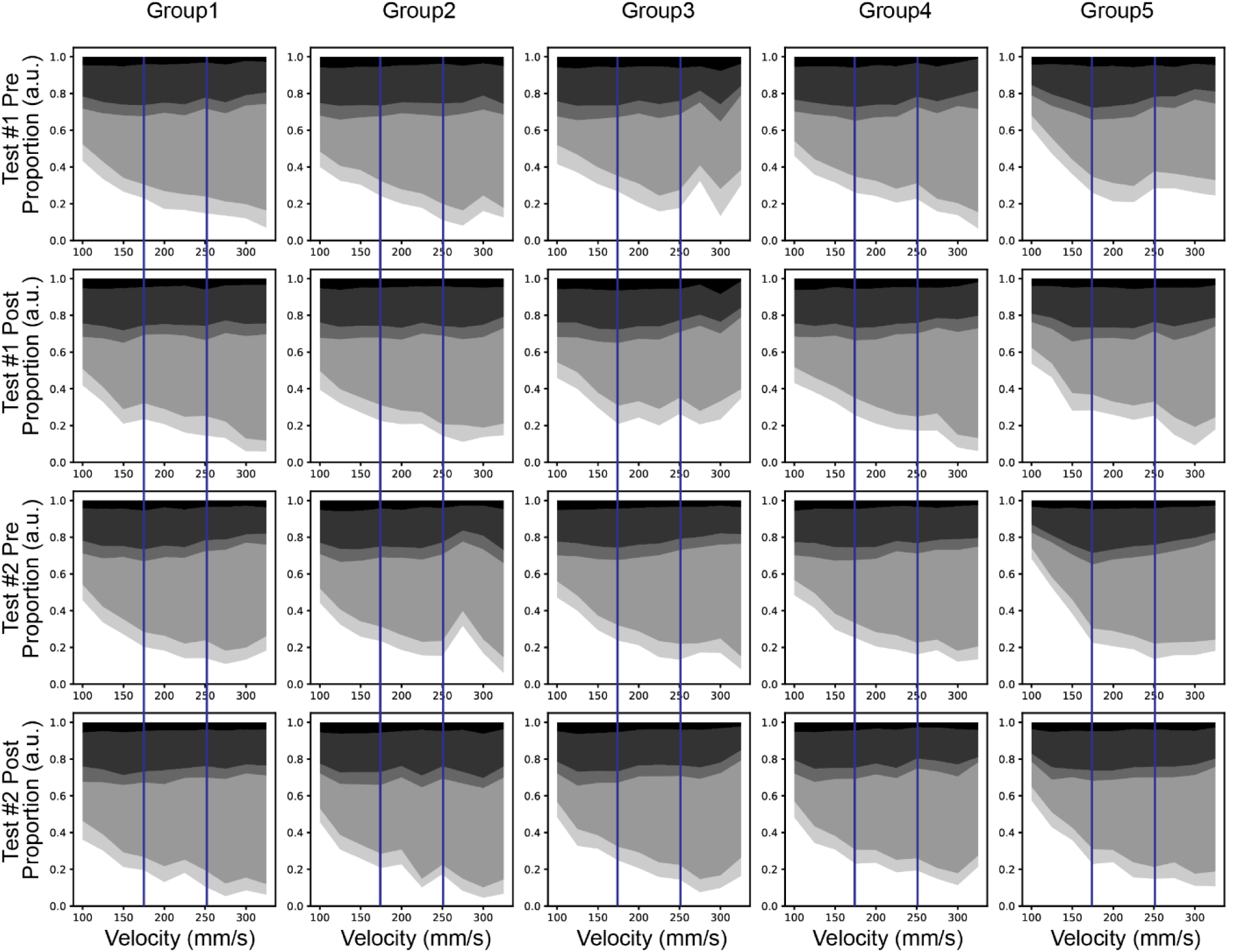
Support phase diagram. From dark to bright shades, each indicates 0 paw, 1 paw, 2 others, 2 diagonal, 3 paws, and 4 paws, respectively. The vertical blue lines are segmenting the slow, medium, and fast velocity regimes.

**Supplementary Fig. 24.**
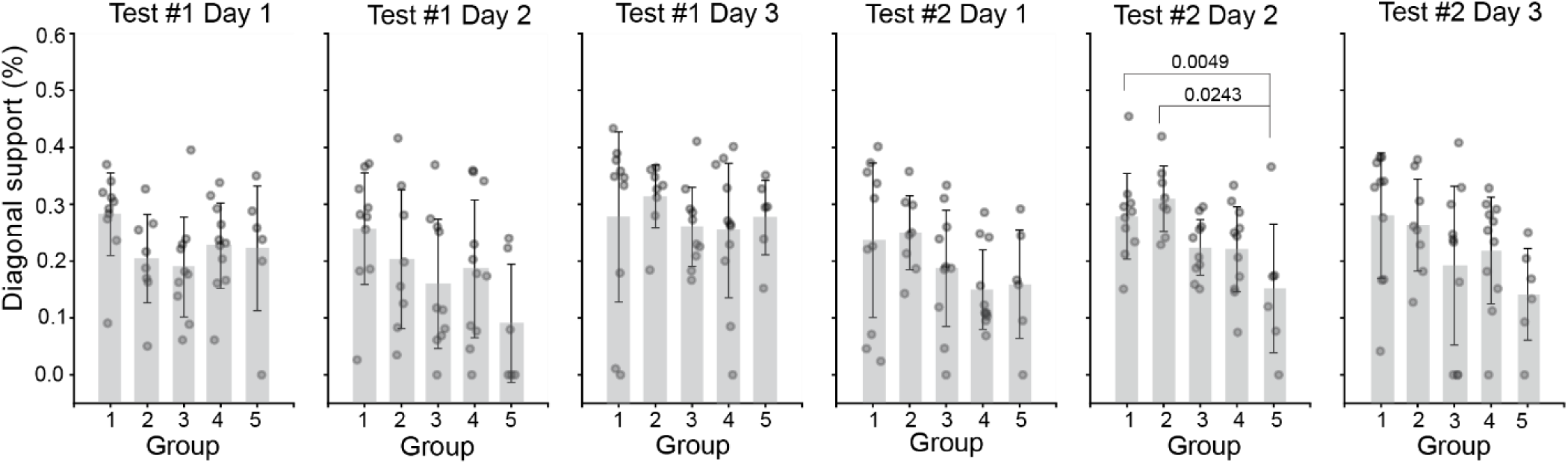
Diagonal Support. Inter-group comparison at pre-stimulation condition at each day. The statistical analysis was performed with One-way ANOVA followed by Tukey’s test; at Test #1 Day 1 F=1.56, p=0.203; at Test #1 Day 2 F=2.00, p=0.113; Test #1 Day 3 F=0.402, p=0.806; Test #2 Day 1 F=1.67, p=0.177; Test #2 Day 2 F=4.40, p=0.00484; Test #2 Day 3 F=1.88, p=0.133. Bars and error bars represent mean ± S.D.

**Supplementary Fig. 25.**
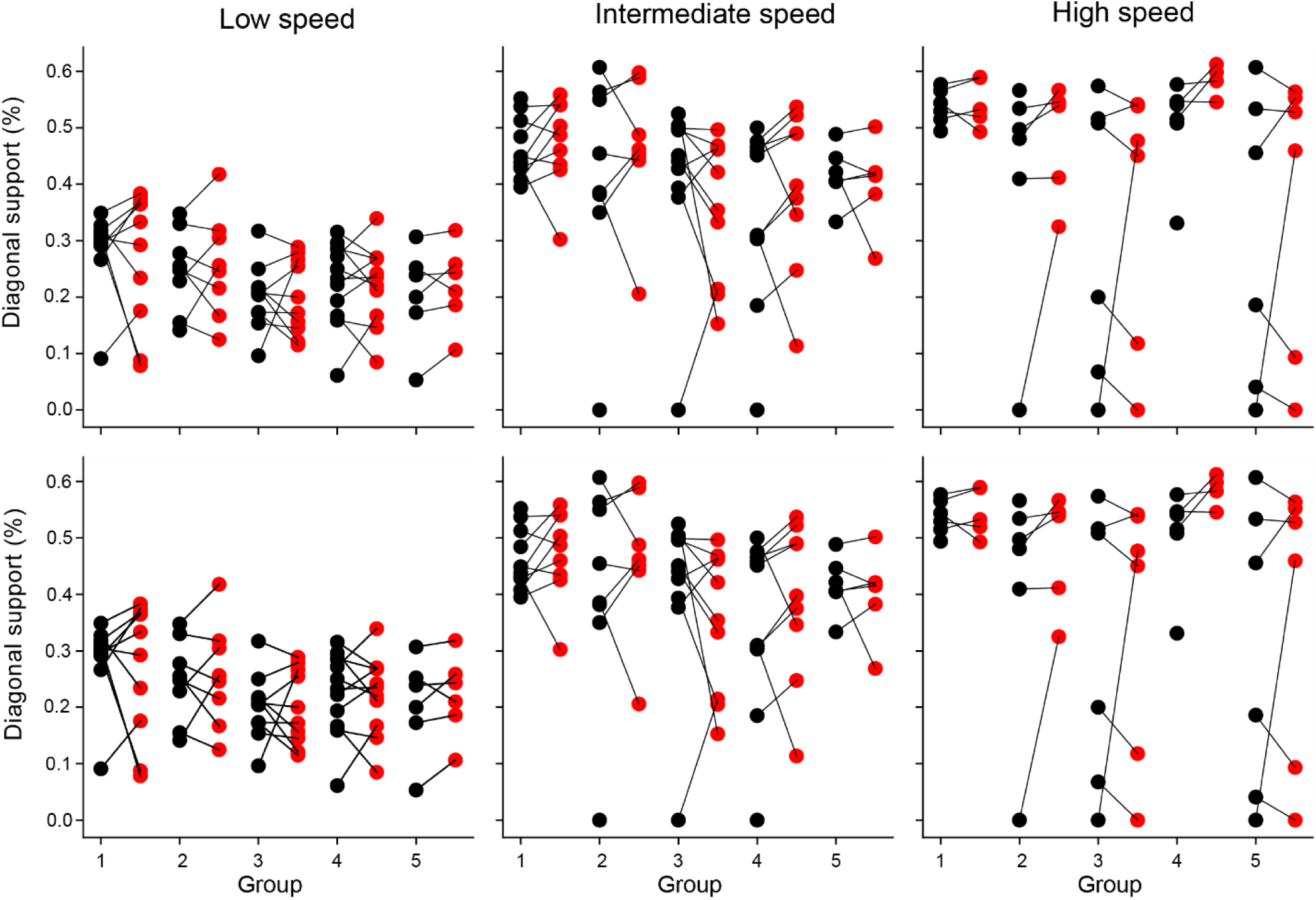
Diagonal support immediately after the stimulation. Pre- (black) and post-stimulation (red) average per animal at Test #1 (top row) and Test #2 (bottom row). Paired t-test was performed with Bonferroni correction for statistical analysis, and no significance was observed in any pair.

**Supplementary Fig. 26.**
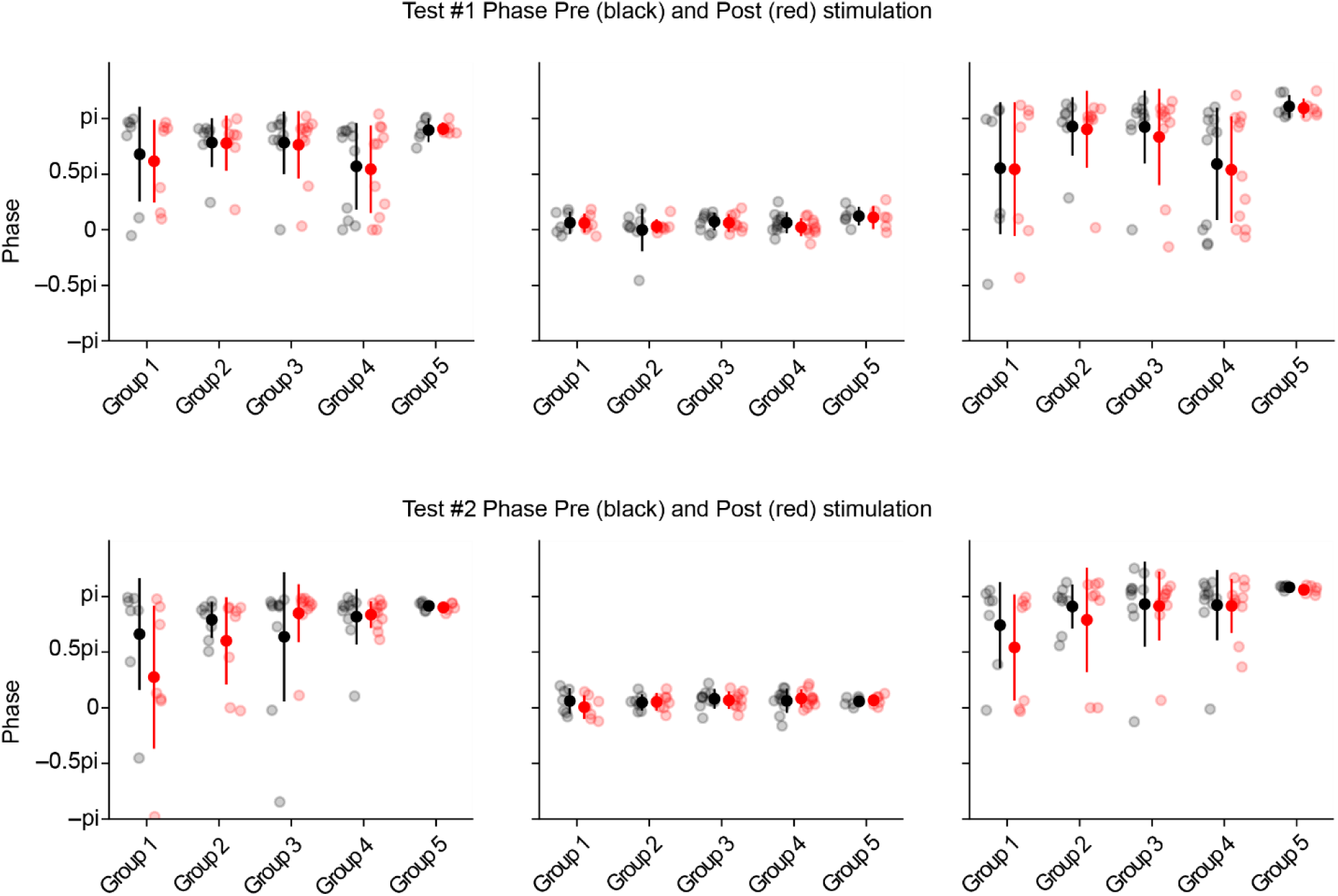
Interlimb coordination pre- and post-stimulation. Black and red, respectively, indicate pre- and post-stimulations. None of the pair showed significant difference. Data is shown as mean ± S.D.

**Supplementary Fig. 27.**
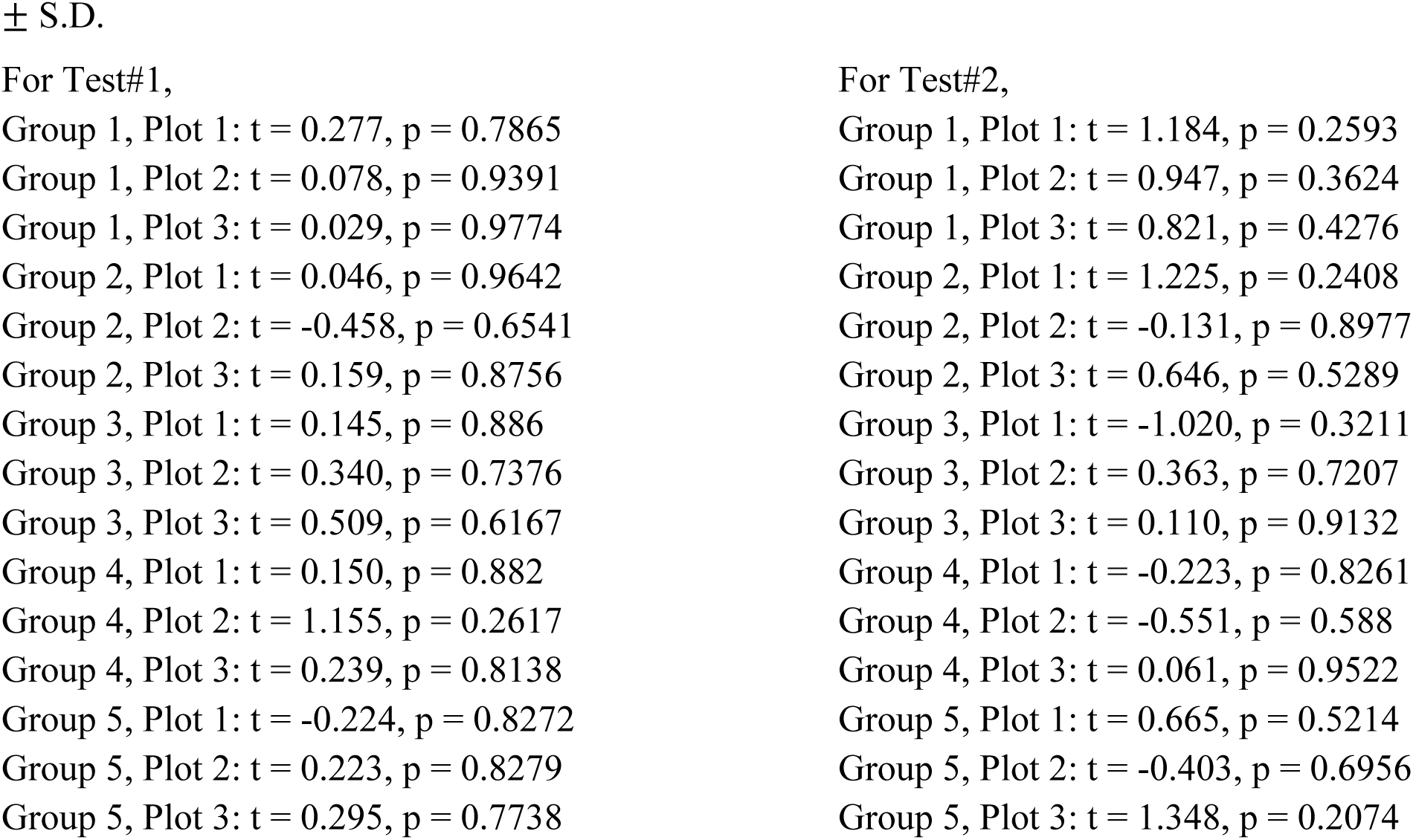
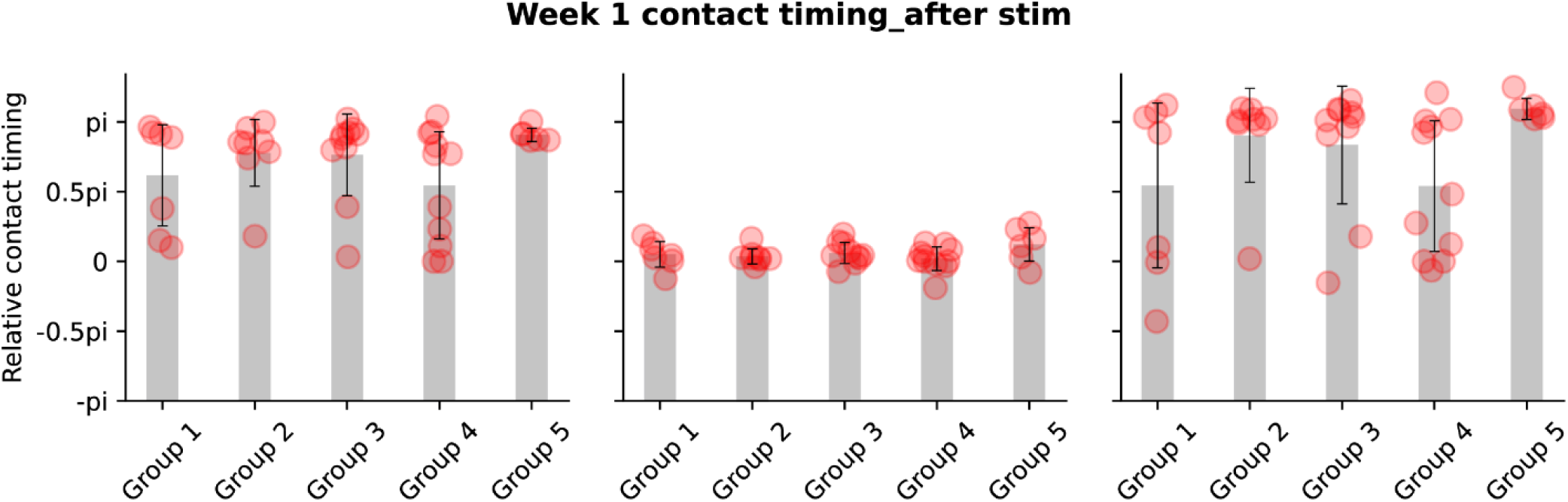
Test 1 post-stimulation gait-phase pattern. Averaged phase at speed 200-300 mm/s for each paw (FL first column, HL second column, and HR third column). The the statistical analysis was performed with One-way ANOVA followed by Tukey’s test; FL F(4,37)=1.57, p=0.20 HL F(4,37)=1.34, p=0.27; HR F(4,37)=2.13, p=0.097. Bars and error bars represent mean ± S.D.

**Supplementary Fig. 28.**
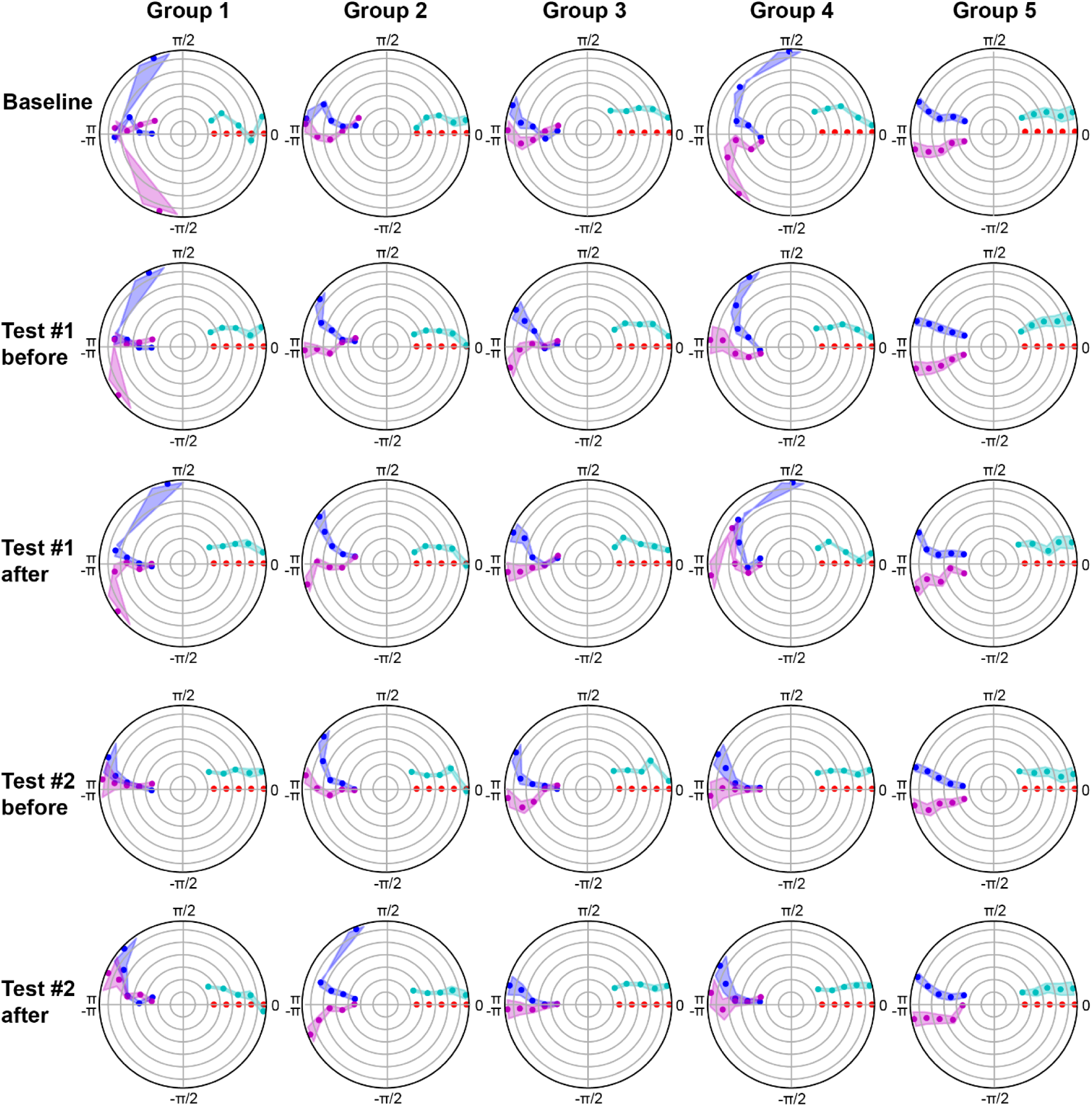
Gait-phase pattern. Polar plots indicating the phase of the step cycle in which each limb enters stance relative to stance onset of front right paw (red): blue is front left, cyan is hind left, and purple is hind right. The baseline (first row) represents pre-stimulation measurements on Day 1 of Test 1. Subsequent rows show values integrated across 3 days for pre- and post-stimulation conditions in Tests 1 and 2, respectively. Columns correspond to Groups 1–5. Markers and shaded areas represent circular mean±s.e.m.

**Supplementary Fig. 29.**
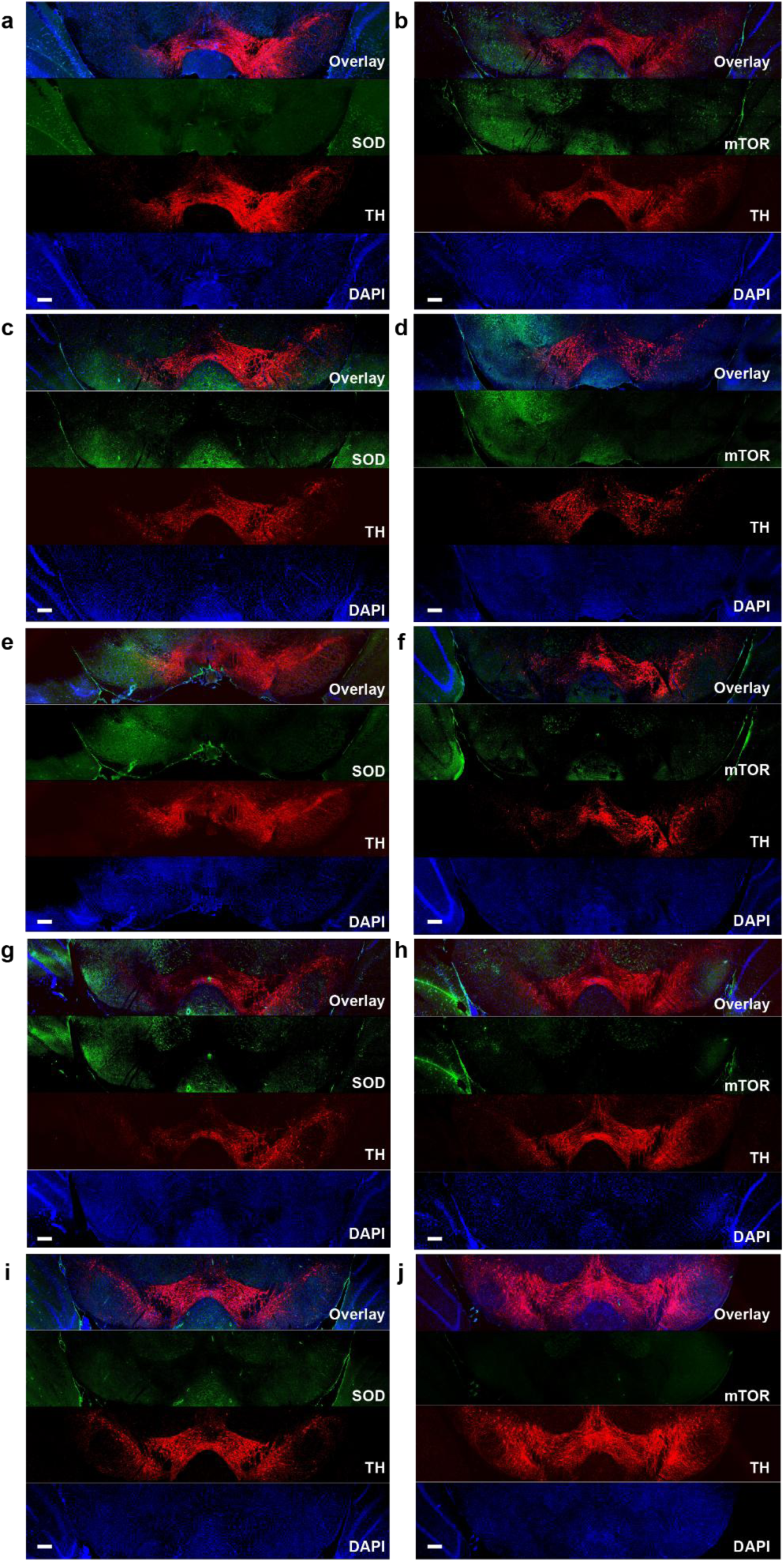
SOD, mTOR, and TH staining. Confocal microscope images for overlay (first row in each panel), **a,c,e,g,i** SOD or **b,d,f,h,j** mTOR (green, second row in each panel), TH (red, third row in each panel), and DAPI (blue, fourth row in each panel) staining in the SN of (**a,b**) Group 1, (**c,d**) Group 2, (**e,f**) Group 3, (**g,h**) Group 4, (**i,j**) Group 5. Scale bars = 100 µm.

**Supplementary Fig. 30.**
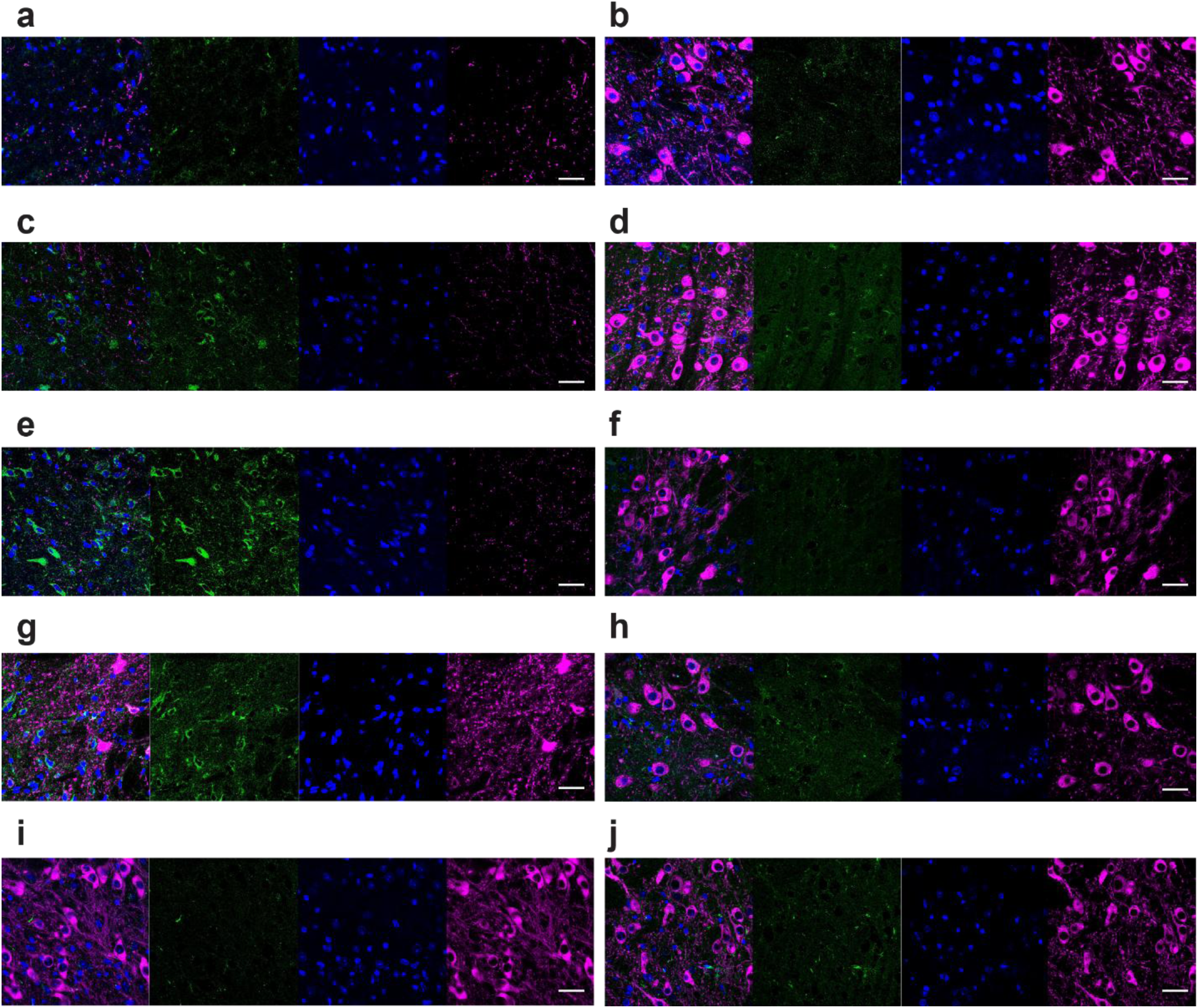
SOD and TH staining. Confocal microscope images for overlay (first column in each panel), SOD (green, second column), DAPI (blue, third column), and TH (magenta, fourth column) staining on SNpc of **a,b** Group 1, **c,d** Group 2, **e,f** Group 3, **g,h** Group 4, **i,j** Group 5. (**a,c,e,g,i**) Left SNpc and (**b,d,f,h,j**) Right SNpc. Scale bars = 25 µm.

**Supplementary Fig. 31.**
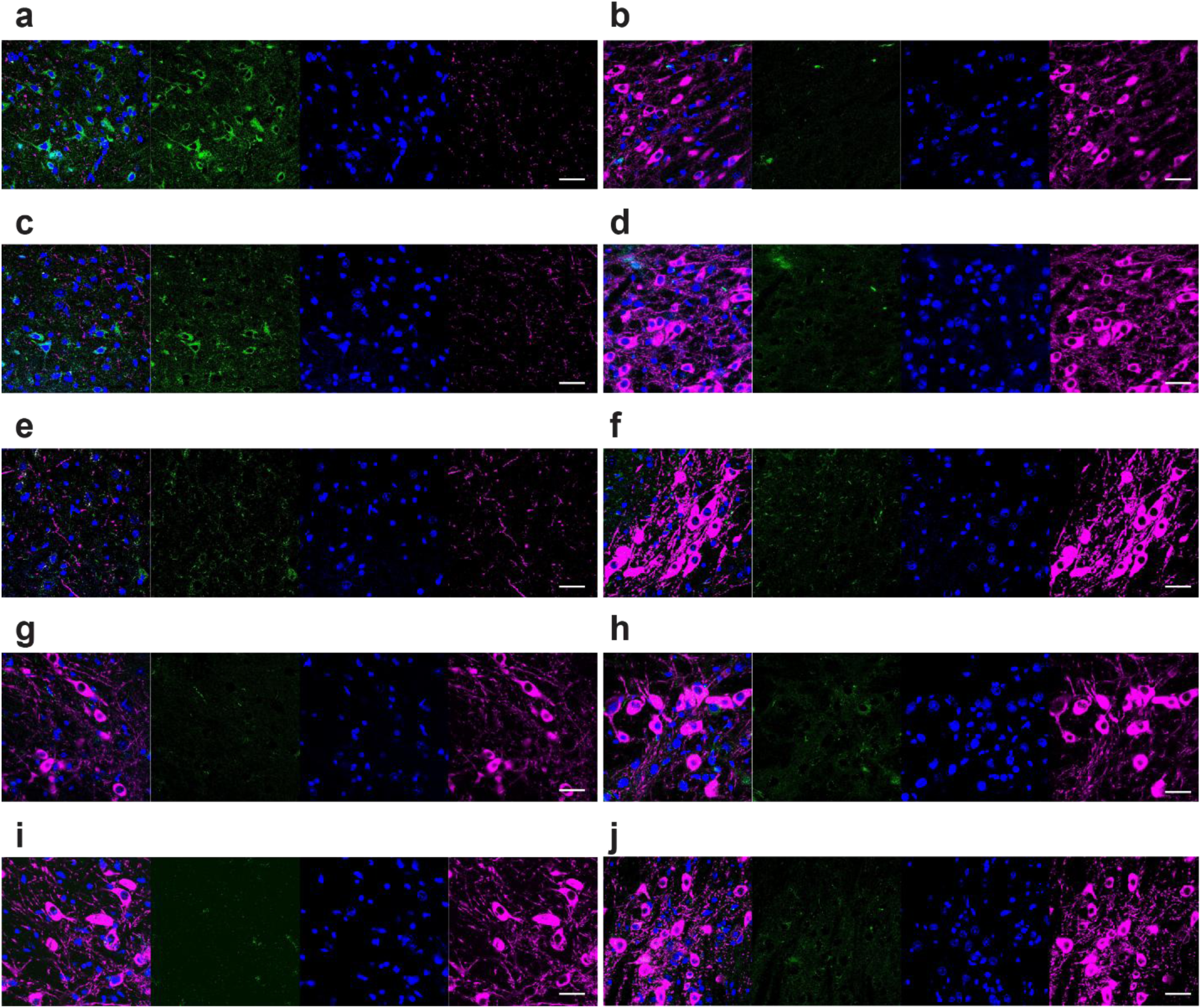
mTOR and TH staining. Confocal microscope images displaying overlays (first column in each panel), along with mTOR (green, second column), DAPI (blue, third column), and TH (magenta, fourth column) stainings in the SNpc. Panels represent different experimental groups: **a,b** Group 1, **c,d** Group 2, **e,f** Group 3, **g,h** Group 4, and **i,j** Group 5. (**a,c,e,g,i**) Images from the left (injected/implanted) SNpc. (**b,d,f,h,j**) Images of the right (unmanipulated) SNpc; 6-OHDA was injected in left MFB and MEND, PBS, or electrode was administered in left STN. Scale bars = 25 µm.

**Supplementary Fig. 32.**
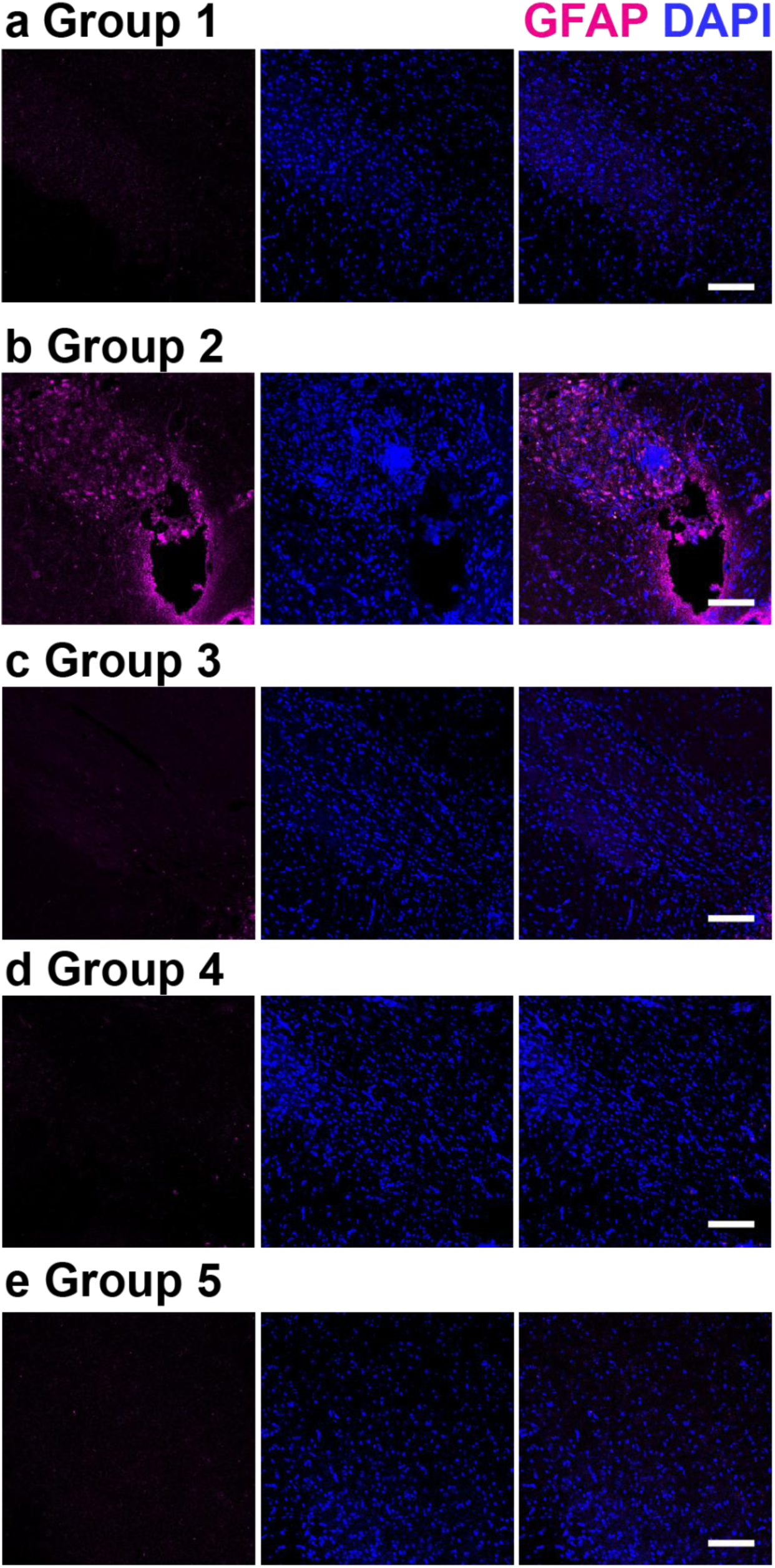
Immunostainig for GFAP. (**a**) Group 1, (**b**) Group 2, (**c**) Group 3, (**d**) Group 4,

**Supplementary Fig. 33.**
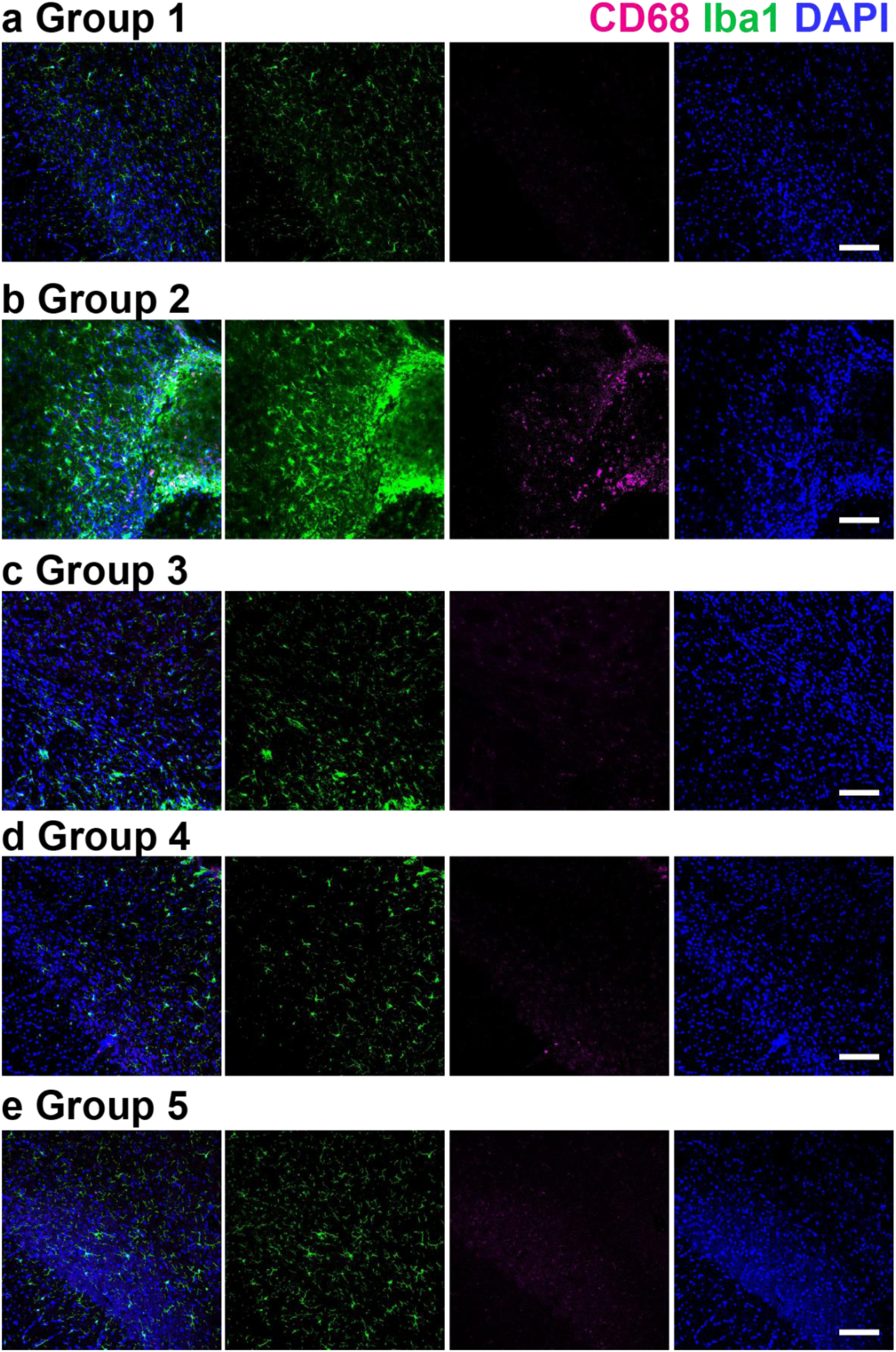
Immunostainig for CD68 and Iba1 for (**a**) Group 1, (**b**) Group 2, (**c**) Group 3, (d) Group 4, and (e) Group 5. Scale bars = 100 μm.

**Supplementary Fig. 34.**
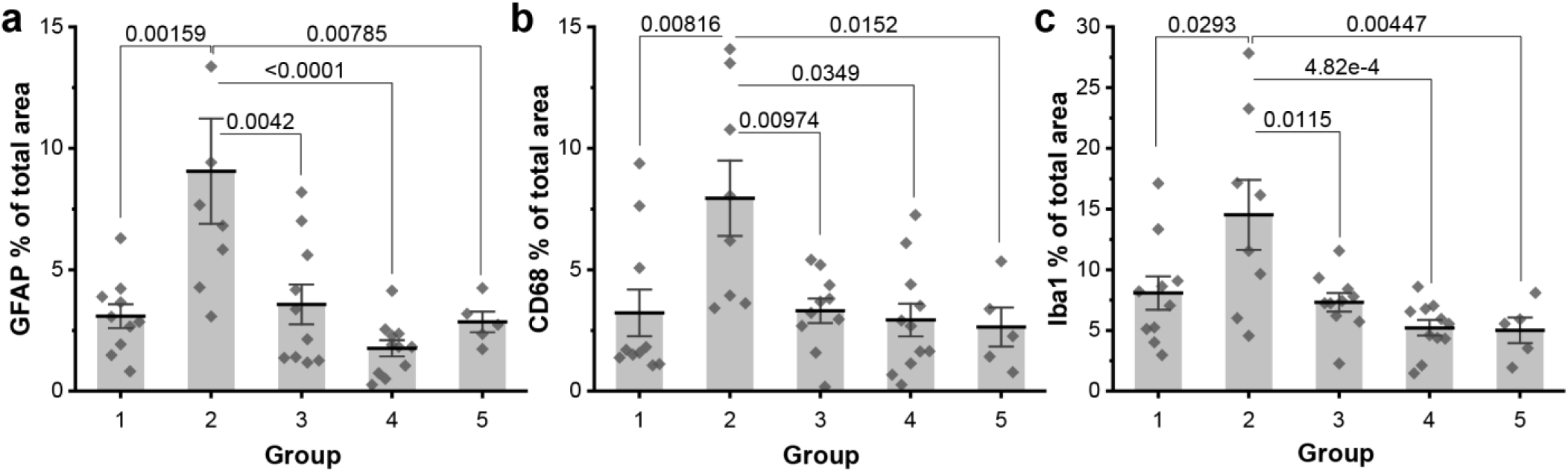
Immune response in the brain. Average percentages of the image areas covered by immunofluorescence of (**a**) GFAP, (**b**) CD68, and (**c**) Iba1 in Supplementary Fig.32 and 33. All analyses were performed after the completion of all behavioral assays. Scale bars = 100 µm. Bars and error bars represent the mean ± S.D.. The Kolmogorov-Smirnov test was performed to test the normality of data distribution. As all datasets were found to conform to normal distribution, the statistical analysis was performed with the One-way ANOVA followed by Tukey’s test. (K) F(4,37)=7.470, p=1.44e-4, (L) F(4,37)=5.048, p=0.00224, (M) F(4,37)=6.105, p=6.491e-4.

**Supplementary Fig. 35.**
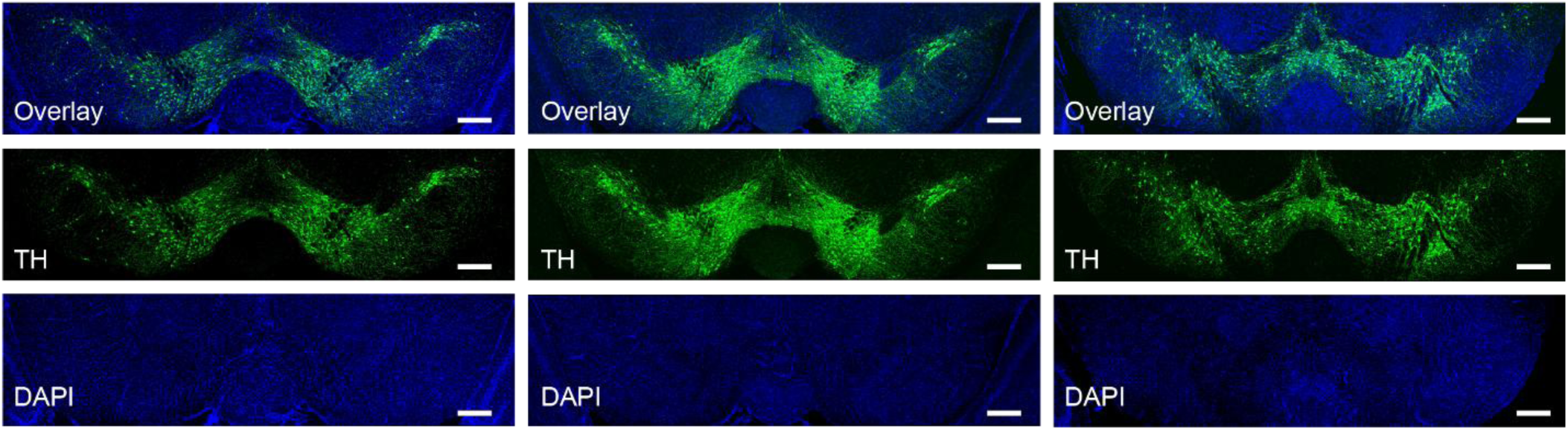
Dopaminergic neurons in the SNpc of mice that have undergone 6-OHDA injections with rapid syringe withdrawal. TH (green) and DAPI (blue) staining in the SNpc of three distinct subjects, analyzed five weeks post-injection of 6-OHDA into the left MFB. The injection procedure involved syringe withdrawal at a rate exceeding 5 mm/min, which led to a backflow of 6-OHDA, manifesting in the lack of observable dopaminergic degeneration. Scale bars = 150 µm.

### Supplementary Movie 1

Example video depicting mouse gait tracing using the computer vision approach.

